# Early Medieval Genetic Data from Ural Region Evaluated in the Light of Archaeological Evidence of Ancient Hungarians

**DOI:** 10.1101/2020.07.13.200154

**Authors:** Veronika Csáky, Dániel Gerber, Bea Szeifert, Balázs Egyed, Balázs Stégmár, Sergej Gennad’evich Botalov, Ivan Valer’evich Grudochko, Natalja Petrovna Matvejeva, Alexander Sergejevich Zelenkov, Anastasija Viktorovna Slepcova, Rimma D. Goldina, Andrey V. Danich, Balázs G. Mende, Attila Türk, Anna Szécsényi-Nagy

## Abstract

The ancient Hungarians originated from the Ural region of Russia, and migrated through the Middle-Volga region and the Eastern European steppe into the Carpathian Basin during the 9th century AD. Their Homeland was probably in the southern Trans-Ural region, where the Kushnarenkovo culture disseminated. In the Cis-Ural region Lomovatovo and Nevolino cultures are archaeologically related to ancient Hungarians. In this study we describe maternal and paternal lineages of 36 individuals from these regions and nine Hungarian Conquest period individuals from today’s Hungary, as well as shallow shotgun genome data from the Trans-Uralic Uyelgi cemetery. We point out the genetic continuity between the three chronological horizons of Uyelgi cemetery, which was a burial place of a rather endogamous population. Using phylogenetic and population genetic analyses we demonstrate the genetic connection between Trans-, Cis-Ural and the Carpathian Basin on various levels. The analyses of this new Uralic dataset fill a gap of population genetic research of Eurasia, and reshape the conclusions previously drawn from 10-11th century ancient mitogenomes and Y-chromosomes from Hungary.

## Introduction

The Ural region was involved in numerous migrations, which events also shaped the history of Europe. The archaeological imprint of these events can be witnessed among others on the early medieval cemeteries of the South-Ural region. Compact cemeteries with few hundred tombs are typical of this territory, which have provided rich archaeological findings first in the last 10-15 years^1–5^. According to archaeological, linguistic and historical arguments, the ethnogenesis of modern Hungarian population can be traced back to the Ural region^1,6,7^.

Based on linguistic evidences, the Hungarian language, belonging to the Ugric branch of the Uralic language family, was developed at the eastern side of Ural Mountains between 1000-500 BC^8,9^. According to the written and linguistic sources and archaeological arguments, after the 6^th^ century AD, part of the predecessors of Hungarians moved to the Western Urals (Cis-Ural region) from their ancient homeland. Around the first third of 9^th^ century AD a part of this Cis-Uralic population crossed the Volga-river and settled near to the Khazarian Khaganate in the Dnieper-Dniester region^1–5,10^ (Fig. 1). Early Hungarians lived in Eastern Europe (forming the so-called Subbotsy archaeological horizon) until the conquest of the Carpathian Basin that took place in 895 AD. The material traits of 10^th^ century AD Carpathian Basin was rapidly transformed after the conquest, its maintained cultural connections with East-European regions have numerous doubtless archaeological evidence^2,4,11^.

**Figure 1.**
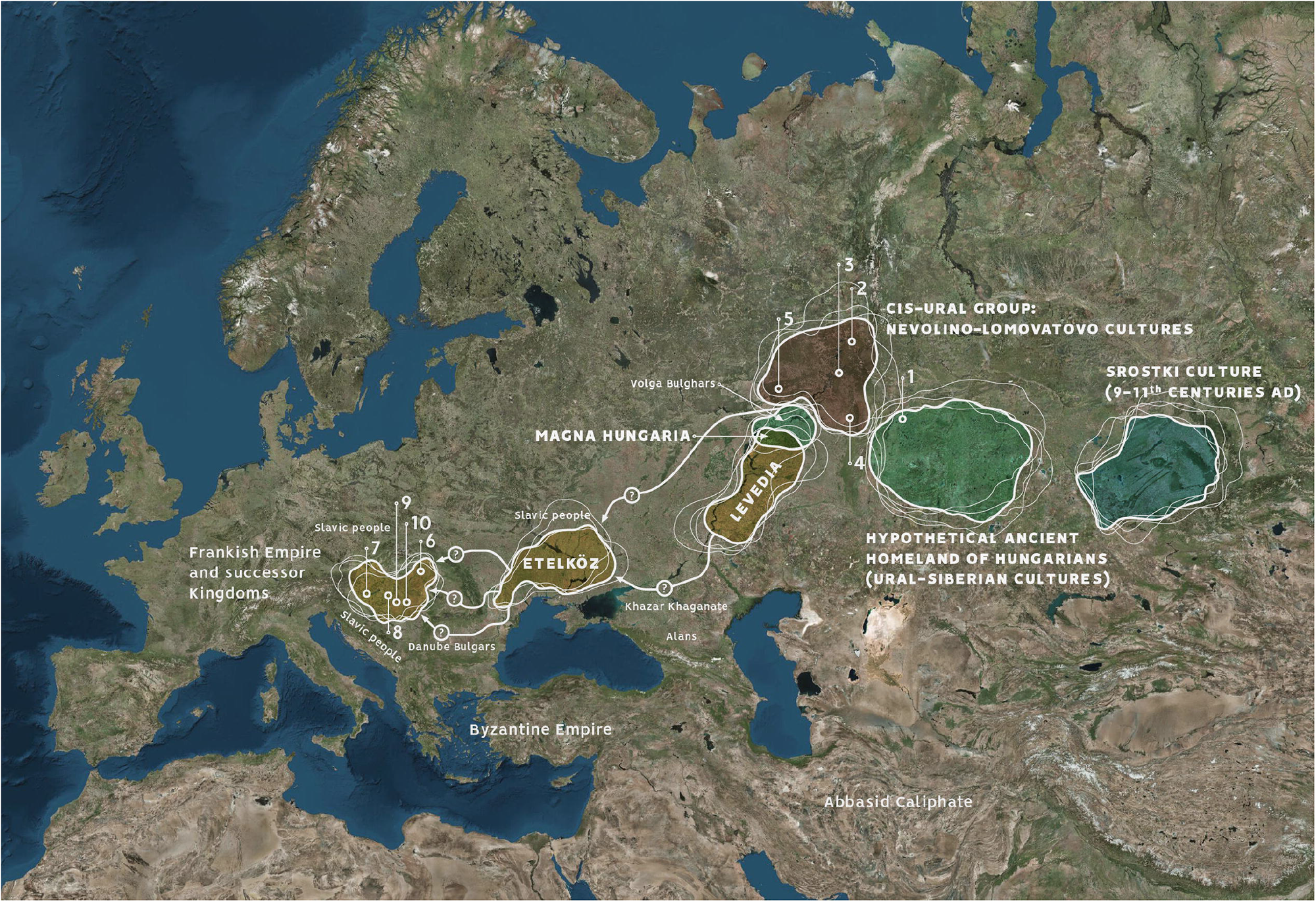
Location of investigated early medieval archaeological sites from South-Ural region and Carpathian Basin, with the possible migration routes and hypothetical Homeland of ancient Hungarians. Trans-Ural region: Uyelgi cemetery (Kushnarenkovo-Karayakupovo culture) (1); Cis-Ural group: Bayanovo (Late Lomovatovo culture) (2), Brody (Early Nevolino culture) (3), Bartym (Nevolino culture, Phase II) (4), Sukhoy Log (Late Nevolino culture) (5); Hungarian conquerors in the Carpathian Basin: Nyíregyháza-Oross, Megapark (6), Balatonújlak-Erdő-dűlő (7), Harta-Freifelt (8), Kiszombor-Tanyahalom (9), Makó-Igási járandó (10). The map of Europe is owned by the IA RCH, and was modified in Adobe 846 Illustrator CS6.

Genetic history of prehistoric to medieval populations of the Ural region have been scarcely investigated to date. On the other side, the populations of the medieval Carpathian Basin have been intensively studied from the perspective of uniparental markers^12,13^. Recently, Neparáczki et al. have published 102 whole mitogenomes from early Conquest period cemeteries in Hungary^14^. Authors have suggested that the mixed population of steppe nomads (Central Asian Scythians) and descendants of the East European Srubnaya culture’s population among other undescribed populations could have been the basis of genetic makeup of Hungarian conquerors. Their results furthermore assume Asian Hunnic-Hungarian conqueror genetic connections^14^. It is important to note, that the investigated medieval sample set does not represent the conqueror population as a whole, hence 76% of the samples originated from a special site complex Karos-Eperjesszög from northeast Hungary, which is one of the most important sites of the Hungarian Conquest period with many findings of eastern characteristics as well. The conclusions are large-scale, but the most highlighted connection with the population of the Srubnaya culture is vague, because it existed more than 2000 years before the appearance of the first traces of ancient Hungarians’ archaeological heritage. Additionally, further mentioned relations such as the Xiongnu (Hunnic) genetic dataset is bare from Eurasia, and Huns’ genetic heritage is basically unknown, as well.

Two recent articles have investigated the Y-haplogroup variability of Hungarian conquerors describing the conqueror’s elite population as heterogenous, with significant proportion of European, Finno-Permic, Caucasian and Siberian (or East Eurasian) paternal lineages^15,16^. Fóthi et al. have claimed that the Hungarian conquerors originated from three distant sources: Inner Asia (Lake Baikal – Altai Mountains), Western Siberia – Southern Urals (Finno-Ugric peoples) and the Black Sea – Northern Caucasus (Northern Caucasian Turks, Alans, and Eastern Europeans)^15^. Both studies^15,16^ pointed out the presence of the Y-haplogroup N-Z1936 (also known as N3a4-Z1936 under N-Tat/M46), which is frequent among Finno-Ugric speaking peoples^17^. This lineage also occurs among modern Hungarians in a frequency up to 4%. Post et al. have reconstructed the detailed phylogeny of N-Z1936 Y-haplogroup showing that specific sublineages are shared by certain ethnic groups, e.g. N-Y24365/B545 by Tatars, Bashkirs and Hungarians, which connect modern-day Hungarians to the people living in the Volga-Ural region^17^.

Earlier mitochondrial DNA (mtDNA) studies of modern populations speaking Uralic languages suggest that the distribution of Eastern and Western Eurasian mtDNA lineages are determined by geographic distances rather than linguistic barriers^18–20^, e.g. Finno-Ugric populations from Volga-Ural region seem to be more similar to their Turkic neighbours than to linguistically related Balto-Finnish ethnic groups^18^. The recent study of 15 Uralic-speaking populations describes their similarities to neighbouring populations as well, however they also share genetic component of possibly Siberian origin^21^. In spite of the unambiguously Central-European characteristics in mtDNA makeup^12,22^, this statement also can be applied to modern day Hungarians^23^.

The main goal of this study is to expand the current set of archaeological knowledge about the early medieval populations of the Ural region by archaeogenetic methods. During the collection of 36 samples from Ural region processed in this study, the most important intention was to collect samples exclusively from such professionally excavated and appropriately documented cemeteries from the South-Ural region, which are culturally and temporally (directly or indirectly) connected to the ancestors of Hungarians (Fig.1 and Supplementary Figs. S1a-h).

The sampled Uyelgi cemetery from Trans-Ural region presented the greatest similarity to the archaeological traits of the tenth-century Carpathian Basin (Figs. 1-2, Supplementary Figs. S1e-h). This cemetery of the late Kushnarenkovo culture was used between the end of 8^th^ century to 11^th^ century^2,24^.

**Figure 2.**
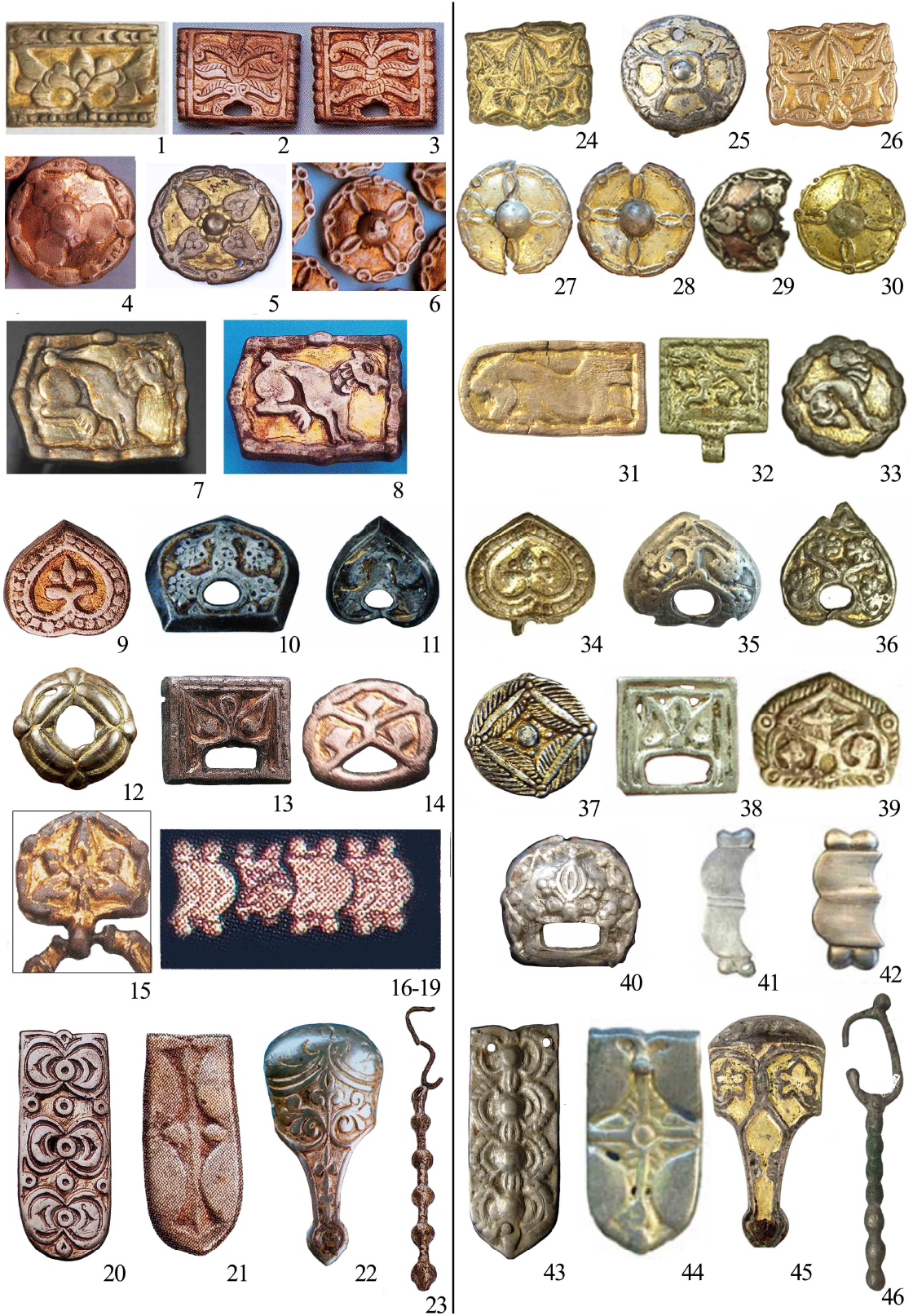
Similar type of the archaeological finds in the heritage of the Hungarian Conquest period (10^th^ century AD) in the Carpathian Basin (left 1–23) and in the material of Uyelgi and Sineglazovo cemetery in Trans-Ural region (right 24–46). The photos were taken by Sergei G. Botalov and Attila Türk.

As the archaeological and historical theories are slightly diverse, we aimed to cover a wide range of early medieval archaeological cultures located in the middle course of the Kama river in the west side of the Ural Mountains (Cis-Ural region). Scholars connect the termination of the Nevolino Culture in 8-9^th^ centuries AD to the westward migration of ancestors of Hungarians^1–3^, hence the sampling was carried out in all three phases of this culture: Brody (3^rd^-4^th^ centuries), Bartym (5-6^th^ centuries) and Sukhoy Log (7-8^th^ centuries)^25^ (Fig. 1). Furthermore, we investigated the Bayanovo cemetery (9-10^th^ centuries AD), which represents the southern variant of Lomovatovo culture^3^ that shows close cultural connection to its southern neighbour Nevolino culture. The sampling of the richly furnished graves of Bayanovo was limited by the poor preservation of bone samples (see Supplementary text, Figs S1b-d)^6^. Additionally, we reanalysed nine samples from tenth-to twelfth-centuries ancient Hungarians for whole mitogenomes from the Carpathian Basin, who were chosen from the previous study Csősz et al.^13^ based on identical hypervariable I region (HVRI) haplotypes of mtDNA with some of investigated Uralic individuals.

In this paper, our main purpose was to characterize the maternal and paternal genetic composition of populations from the third-to eleventh-centuries South-Ural region and compare the results with the available ancient and modern genetic datasets of Eurasia. We also aimed to describe possible genetic connections between the studied Uralic populations and the Conquest period populations of the Carpathian Basin.

## Results and Discussion

The sample-pool consisted of 29 males and 16 females. We performed whole mitochondrial DNA and 3000 nuclear SNP target-enrichment combining with shallow shotgun sequencing. With the latter we obtained autosomal and Y-chromosomal SNPs, as well as sex-determination of 45 individuals that originated from five different cemeteries in Ural region and six burial sites in present-day Hungary (Carpathian Basin). Furthermore, we investigated the Y-STR profiles of 20 male individuals from the Ural-region. For detailed information see Supplementary Tables S1, S2. For the radiocarbon dating and stable isotope data see the Supplementary Information chapter 2 and Supplementary Table S1.

### Primary observations

45 high coverage mitochondrial genomes were obtained (sequencing depth from 8.71× to 154.03×), with mean coverage of 71.16× and an average contamination rate of 0,2%. The new dataset consists of the mixture of nine macrohaplogroups (A, C, D, H, T, U, N, R, Z) (Fig. 3a). Haplogroups of presumably west Eurasian origin are represented by U (U2e1, U3a1, U4a1d, U4b1a1a1, U4d2, U5a1a1, U5b2a1a1, N=12), H (H1b2, H3b, H40b, N=9), N (N1a1a1a1a, N=5) and T (T1a1, T1a2, T2b4h, N=5), although phylogeographic analyses show eastern origin for some of them, see Table1 and Supplementary Figs. S4a-s. Eastern Eurasian lineages are represented by A (A+152+16362, A12a, N=4), C (C4a1a6, C4a2a1, N=6), D (D4j, D4j2, N=2), along with R11b1b and Z1a1a by one individual each (Fig. 3a).

**Figure 3.**
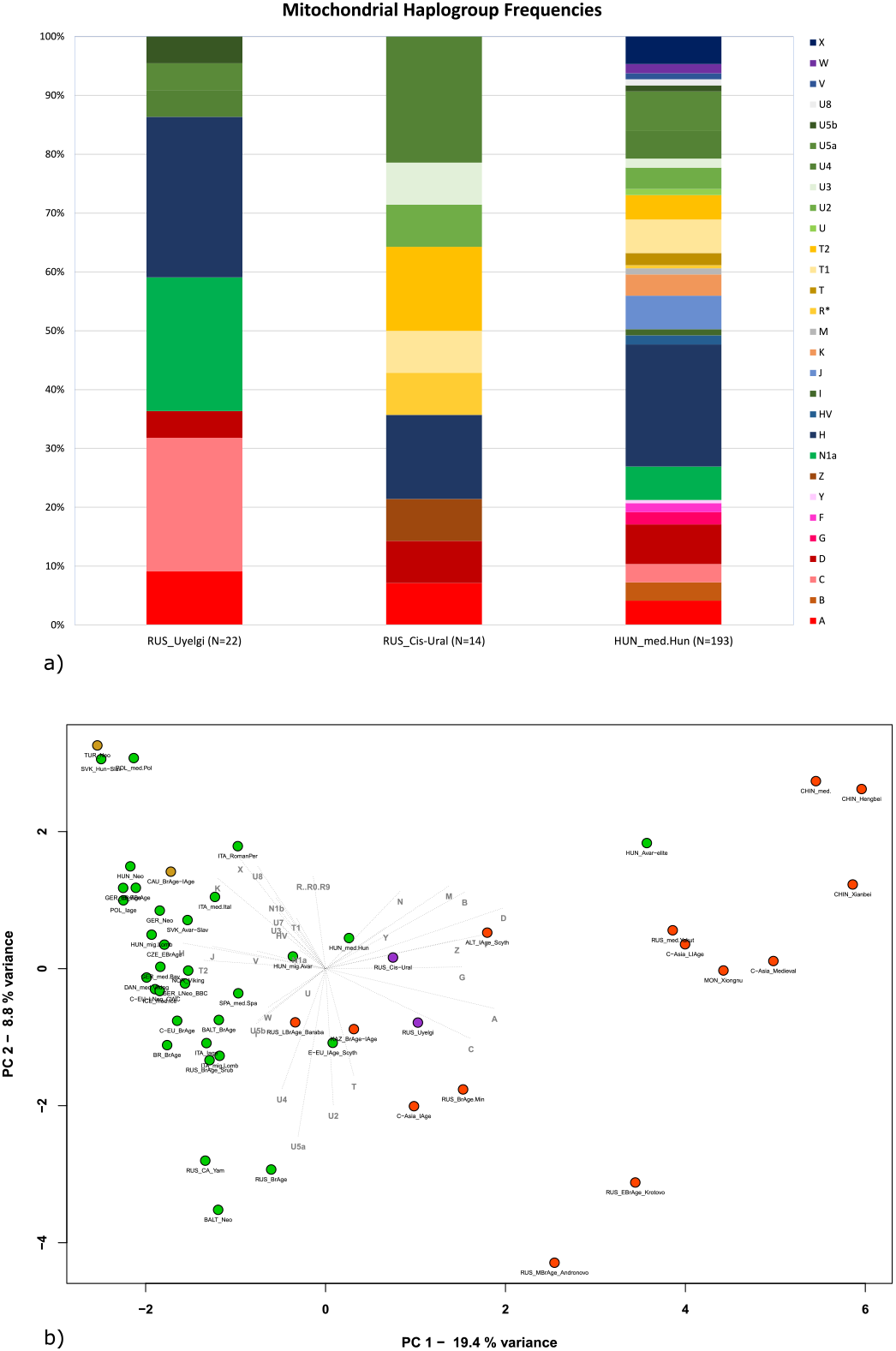
Mitochondrial haplogroup frequencies of the three investigated populations (a) and PCA plot with 50 ancient populations, representing first and second principal components (b). a) The characterized populations: Hungarian conquerors (HUN_med.Hun), data replenished by previously studies (Neparáczki et al. 2017^14,57^, Csősz et al. 2017^13^, Tömöry et al. 2007^12^); cemeteries Bayanovo, Sukhoy Log, Bartym and Brody (Nevolino-Lomovatovo cultures) grouped into “Cis-Ural” (RUS_Cis-Ural); cemetery Uyelgi (RUS_Uyelgi) from the Trans-Ural region. (see Supplementary Tables S2 and S4). b) PCA analyses based on haplogroup frequencies in Eurasian ancient populations. Clear separation of Asian (red) and European (green) populations is visible on the plot the investigated Cis-Ural and Uyelgi (violet-coloured) sites are located between them: the Cis-Ural (RUS_Cis-Ural) near to the Hungarian conquerors (HUN_med.Hun) and the Uyelgi (RUS_Uyelgi) is positioned between the Iron Age population from Central Asian Steppe (C-Asia_IAge), Russian Bronze Age population from Minusinsk Depression (RUS_BrAge.Min), Bronze Age and Iron Age populations from Kazakhstan (KAZ_BrAge-IAge) and the East European Iron Age Scythians (E-EU_IAge_Scyth).

**Table1.** Summary of the phylogeographic origin of the mitochondrial subhaplogroups detected by investigated samples.

| Sites where maternal lineage was found | MtDNA haplogroup | Most probable geographic origin of corresponding lineage according to phylogeography |
| --- | --- | --- |
| Uyelgi, Sukhoy Log | A+152+16362 (N=2) | Kazakh steppe (central-east) |
| Uyelgi | C4a1a6 (N=4) |  |
| Sukhoy Log | R11b1b (N=1) |  |
| Uyelgi | D4j (N=1) | Kazakh steppe |
| Uyelgi, Harta-Freifelt | A12a (N=2) | Kazakh steppe (central-east) or Baraba steppe (south-east) |
| Uyelgi, Makó-Igási járandó | C4a2a1 (N=2) |  |
| Brody | D4j2 (N=1) | Pontic-Caspian steppe |
| Kiszombor | T1a2 (N=1) | Pontic-Caspian steppe or Caucasus |
| Sukhoy Log, Nyíregyháza | H1b2 (N=3) | East European Plain |
| Uyelgi | H40b (N=5) |  |
|  | N1a1a1a1a (N=5) |  |
| Bayanovo, M3 161. lelőhely | T1a1 (N=2) |  |
| Bayanovo, Bartym | T2b4h (N=2) |  |
| Bartym | U2e1 (N=1) |  |
| Sukhoy Log | U3a1 (N=1) |  |
| Bartym | U4b1a1a1 (N=1) |  |
| Uyelgi, Bartym | U4d2 (N=2) |  |
| Uyelgi, Balatonújlak-Erdő-dűlő, Makó-Igási járandó | U5a1a1 (N=4) |  |
| Uyelgi | U5b2a1a1 (N=1) |  |
| Harta-Freifelt | U5a1a1+152 (N=1) | Northern Europe |
| Bartym | U4a1d (N=1) | Ural region |
| Bayanovo | Z1a1a (N=1) | Central-Southwestern Ural region |

Even though that the Hungarian conquerors were selected based on mtDNA HVRI matches with certain ancient individuals from the Ural region, they have not proved to be identical on whole mitogenome level, but remained phylogenetically close to the associated samples (see Supplementary Figs. S4a-s).

A few mitochondrial lineage relations connect Trans-Ural and Cis-Ural regions: e.g. samples from Uyelgi and Sukhoy Log clustered together in one main branch of the A+152+16362 haplogroup tree (Supplementary Fig. S4b), furthermore samples from Uyelgi and Bartym (with haplogroup U4d2) are located on the same main branch as well (Supplementary Fig. S4p).

The sole investigated sample from Brody cemetery with haplogroup D4j2 neither show close maternal genetic connection to other Uralic samples nor to Hungarian conquerors.

In contrast to the mitochondrial lineages, the Y-chromosomal gene pool based on STR and/or SNP data show homogenous composition in our dataset: 83.3% is N-M46, 5.5% G2a (G-L1266), 5.5% J2 and 5.5% is R1b of the typed male individuals (Supplementary Table S2). 13 male samples out of 19 from Uyelgi cemetery carry Y-haplogroup N with various DNA preservation-dependent subhaplogroup classifications, while in the Cis-Ural we detected three N-M46 Y-haplogroups (samples from Brody, Bartym and Bayanovo cemeteries). The overall poor preservation of further Cis-Uralic samples from Sukhoy Log and Bartym disabled further Y-chromosome-based analyses (Supplementary Table S2).

### Comparative population genetic analyses of maternal lineages and genomic data

We performed population genetic statistical analyses as well. The principal component analysis (PCA) and Ward clustering of 50 ancient and 64 modern populations were performed separately (Fig. 3b, Supplementary Figs. S5-S8), based on haplogroup frequencies (Supplementary Tables S3 and S4). The Hungarian conquerors are the closest population to the Cis-Ural group on the PCA (along PC1 and PC2 components, see Fig. 3b) and this population is relatively near to the Uyelgi among the Iron Age population from Central-Asia and the East European Scythians along PC1 and PC3 components (Supplementary Fig. S5), because these ancient populations have mixed pool of western and eastern Eurasian macrohaplogroups, which is unusual in European and Asian populations that are separated along the PC1. The nearby position of Cis-Ural and Uyelgi to the Hungarian conquerors is displayed on the mtDNA haplogroup-based Ward type clustering tree too, where they appear in the same main branch (Supplementary Fig. S6). Some of Central-South Asian and Finno-Ugric modern populations (e.g. Khanty and Mansi) show close connections to the investigated Cis-Ural and Uyelgi populations based on Ward-type clustering and PCA (Supplementary Figs. S7 and S8). The haplogroup frequencies of three highlighted populations are displayed on the Fig.3a diagram. The mitochondrial haplogroup pool of the Hungarian conquerors’ large sample-set is the most diversified and contains nearly all haplogroups obtained in two populations from Ural region with a similar proportion of haplogroups with western and eastern Eurasian origin. This phenomenon causes their relatively nearby positions on the PCA and Ward clustering tree.

Pairwise FST values of populations indicate non-significant differences of the Cis-Ural from 13 ancient populations (Supplementary Table S5), among them the Hungarian conquerors^14^ show the lowest genetic distance (F_ST_ = 0.00224) (for further F_ST_ values, p values, and references see Supplementary Table S5). According to the MDS plot of 28 ancient populations based on linearized Slatkin F_ST_ (Supplementary Fig. S9a), the Cis-Ural population shows affinities among others to the populations of medieval Hungarian conquerors along coordinates 1 and 2, and is situated between European and Asian populations, which reflects the raw F_ST_ values. The Uyelgi is standing on the Asian part of the plot relatively far from all ancient populations, which is most likely due to its significant and larger genetic distances from ancient populations (except the Late Iron Age population from Central Asia^26^) and the scarcity of Asian comparative mitogenome datasets. The rank correlation heatmap (Supplementary Fig. S9b) of the F_ST_ values of ancient populations supports the MDS plot, where the Uyelgi and Cis-Ural populations cluster with the same ancient populations that are close to them on the MDS plot.

The genetic connection of Cis-Ural population and Hungarian conquerors^14^ is obvious based on pairwise F_ST_ calculation and is visible on the PCA and MDS plots as well, where they are the closest, although direct phylogenetic connections are scarce. This indicates geographical proximity of their former settlement area, rather than a direct connection. Neparáczki et al.^14^ have described the Hungarian conqueror mitogenome diversity in essence as a mixture of Srubnaya and Asian nomadic populations. Their analyses and interpretation were restricted by the lack of ancient samples from the Ural region, whereas new data now refine such previous conclusions^14^. Furthermore, it is notable, that the previously studied Hungarian conqueror population is a pool of mixed origin including not only immigrants but also local admixed lineages from the Carpathian Basin.

The Cis-Ural population reveals non-significant genetic distances from four modern populations of Central Asian Highlands, furthermore seven populations of Near East and Caucasus region and six European populations (see Supplementary Table S6) indicating a mixed character of this population, which is also visible on the MDS plot.

Interestingly, the mitogenome pool of Uyelgi shows significant differences in genetic distances among nearly all prehistoric and modern populations including Hungarian conqueror population in spite of the extensive phylogenetic connections, which might be explained by high amount of related lineages within the population, as well as by their mixed character of Eastern- and Western-Eurasian haplogroups.

We performed genomic PCA of five Uyelgi samples consisting of 10,828 nuclear genomic SNPs on average gained from 3000 SNP capture and shallow shotgun sequencing data (from 598,094 called SNPs). The five samples are plotted together on the genomic PCA and they also appear close to the modern Bashkir and Siberian Tatar individuals as well as to the Altaian Bronze Age Okunevo population^27^, to a hunter-gatherer individual from Tyumen region^28^ and Iron Age Central Sakas from Kazakhstan^26^ (see Supplementary Information, chapter 3 and Supplementary Figs. S3a-c) in line with the uniparental makeup. Since PCA may not reveal population stratification we performed unsupervised ADMIXTURE (K=16) on an enlarged sets of SNPs (SI, chapter 3). The five Uyelgi samples with an average calling of 22,540 SNPs show the most similar ancestry cluster proportions to present-day Mansis and Irtysh-Barabinsk Tatars and to a set of various populations lived in the Central Steppe region^27^.

To disentangle the connections between these populations and possible population genetic events of thousands of years between populations under study, more ancient reference samples and deeper sequencing for more detailed analyses are needed.

### The genetic continuity between the horizons of the Uyelgi cemetery (Trans-Ural region)

The kurgan burials at Uyelgi site can be divided into at least three chronological horizons: I.) the oldest ninth-century, II.) ninth- and tenth-centuries and III.) tenth- and eleventh-centuries according to the archaeological records (see Supplementary text chapter 1 and Supplementary Figs. S1e-h). Uniparental genetic markers show genetic continuity between these horizons suggesting maternally rather endogamous population, which could not be observed in archaeological findings due to high number of disturbed burials in the cemetery. Mitochondrial phylogenies of N1a1a1a1a, C4a1a6 and H40b provide identical or monophyletic lineages within and between the three horizons (see Figs. 4-5 and Supplementary Figs. S4h-g), which trend is more pronounced by haplotype and network analysis of paternal lineages (Fig. 6., Supplementary Figs S11-12).

**Figure 4.**
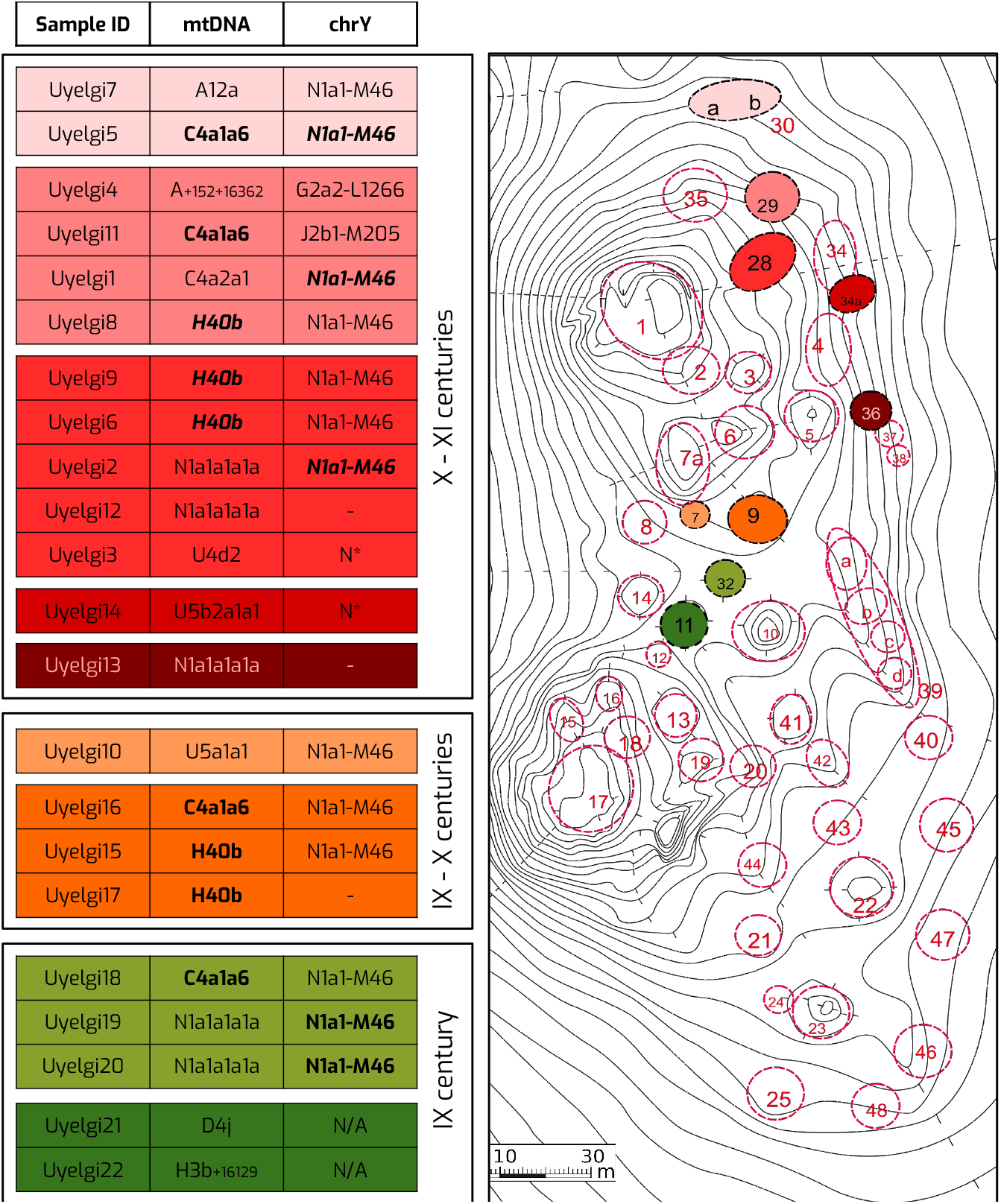
Mitochondrial haplotype and Y-chromosomal similarities between the kurgans of the three horizons of Uyelgi cemetery. Three chronological horizons were defined in the cemetery: an oldest horizon from 9^th^ century (marked with green), a middle horizon from 9-10^th^ centuries (marked with orange) and the youngest horizon from 10-11^th^ centuries (marked with red colour). The bold and *italic* highlighted letters indicate different mitochondrial and/or Y-STR haplotype matches within and between the kurgans whose localization is visible on the right part of the figure. Four identical mitogenome haplotypes belonging to C4a1a6 haplogroup appear in all three horizons in four different kurgans, furthermore two identical haplotypes of H40b haplogroup are from the middle horizon and three also identical but different from the previous haplotypes of H40b come from two kurgans of the youngest horizon. Additionally, the five individuals with mtDNA haplogroup N1a1a1a1a are distributed in three kurgans from the oldest and youngest horizons.

**Figure 5.**
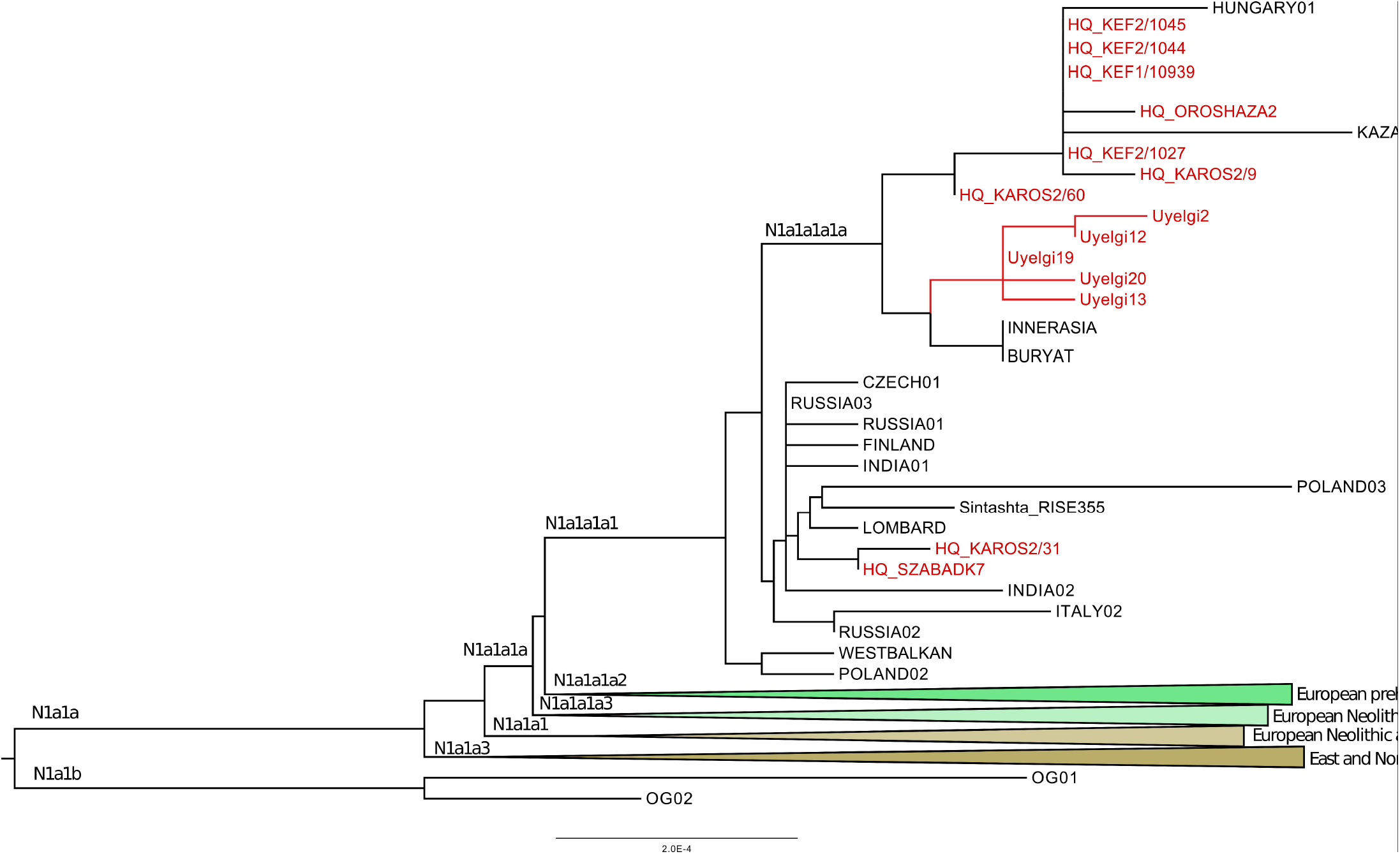
Phylogenetic tree of mitochondrial haplogroup N1a1. The subhaplogroup N1a1a1a1a was detected in five individuals assigned to two horizons of Uyelgi cemetery: Uyelgi19 and Uyelgi20 from the oldest (9^th^ century) horizon and the Uyelgi2, Uyelgi12 and Uyelgi13 from the youngest horizon (10-11^th^ centuries), furthermore, in nine 10^th^ centuries Hungarian conqueror graves from various cemeteries in Hungary, and in one modern Hungarian individual. The Uyelgi branch of the tree is very compact, clearly connects the oldest and youngest horizons together, however, the maternal lineages of the populations from Uyelgi and the Hungarian conquerors are separated (for the abbreviation and further information see Table S8).

**Figure 6.**
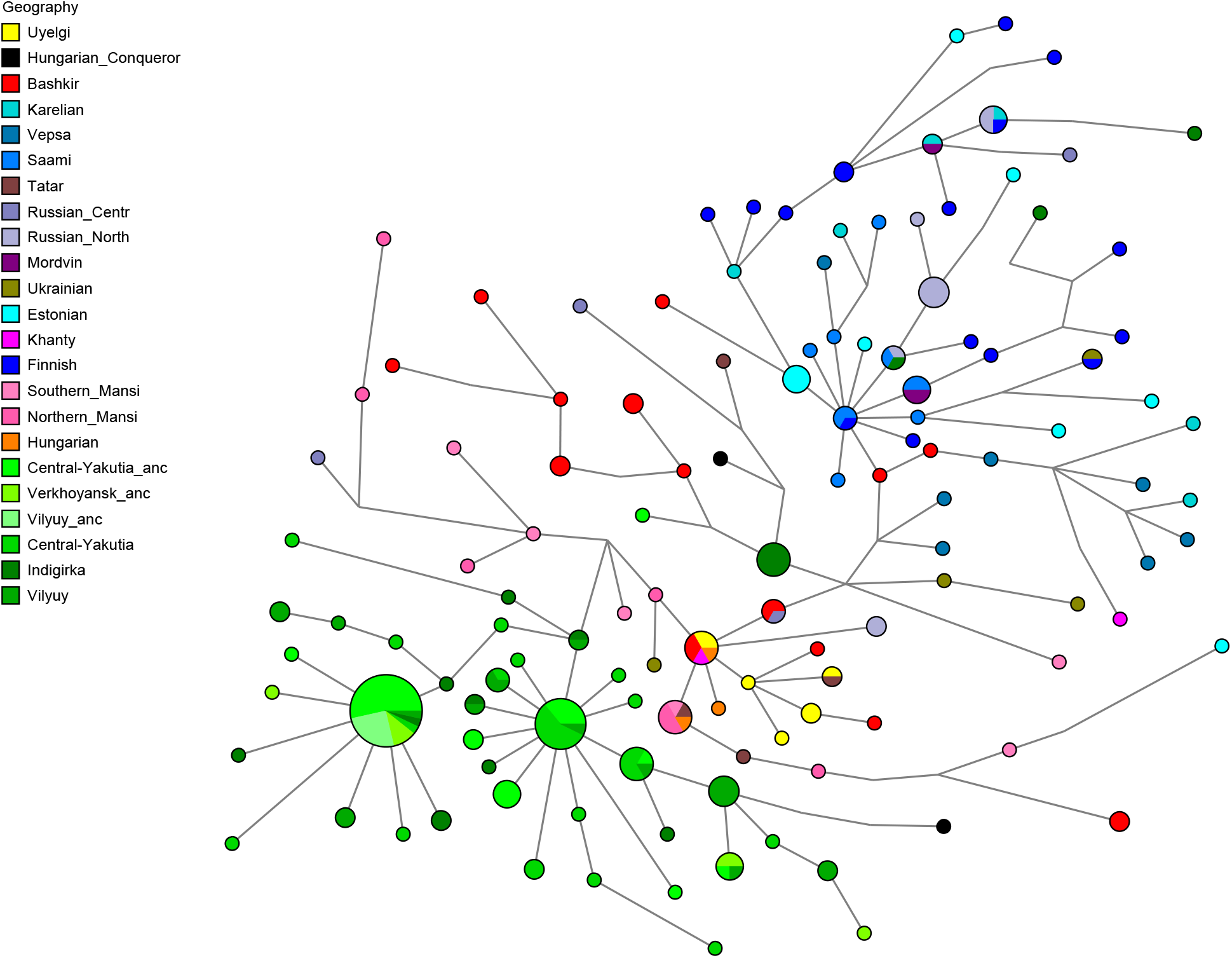
Median Joining Network analysis of the N-M46 Y-chromosomal haplogroup based on 17 STRs in 238 samples. All seven samples from Uyelgi (marked with yellow) besides two-two identical samples (Uyelgi1-Uyelgi5 and Uyelgi19-Uyelgi20) are one-step-neighbours to each other, as well as to five Mansi, four Bashkir, two Hungarian samples, one Tatar sample from Volga-Ural region and one Central Russian sample. The Uyelgi1 and Uyelgi5 share identical STR haplotype with two present-day Bashkir individuals from Volga-Ural region, one Khanty individual from Western-Siberia and one Hungarian individual. The sample Uyelgi16 has identical STRs with a Tatar individual from Volga-Ural region (see Supplementary Table S10).

The haplotypes of N-M46 Y-haplogroup are presented in all three horizons, however with little differences in STR profiles (Supplementary Table S2). The oldest and the middle horizons contain only N-M46 haplotypes including two identical STR profiles in Kurgan 32 (9^th^ century). Three identical Y-STR profiles are detected among individuals of Kurgans 28, 29 and 30 (Fig. 4 and Fig. 6). Probably further identical Y-haplotypes could have been in this cemetery, but the preservation has not let us reconstruct whole Y-STR profiles of seven males (see Supplementary Table S2). Based on these results we suggest that Uyelgi cemetery was used by a patrilocal community.

The genetic continuity between the 9–11^th^ centuries is also supported by genomic data (Supplementary Figs. S3a-c). The Uyelgi2 sample of the youngest horizon (10– 11^th^ centuries) has high proportion of shared drift with the Uyelgi10 of the 9–10^th^ centuries.

### The possible maternal genetic connection of South-Ural region’s populations and the Hungarian conquerors

The genetic connection of Uyelgi cemetery in the Trans-Ural and 10^th^ century Hungarian conquerors in the Carpathian Basin is supposed by close maternal relationships of the following individuals: Uyelgi3 from Kurgan 28 of the youngest horizon and three Hungarian conquerors from Karos II cemetery^14^ have identical U4d2 mitogenome haplotype (Supplementary Fig. S4p). Furthermore, the mtDNA A12a lineage of Hconq3 (30-40 years old woman from Harta cemetery dated to the first half of 10^th^ century AD) is an ancestor of the mtDNA lineage of Uyelgi7 (from Kurgan 30 of the youngest horizon of the cemetery) based on the A12a haplogroup tree (see Supplementary Fig. S4a).

The mentioned graves from Uylegi show the characteristic of the Srostki culture, where the gilt silver mounts with plant ornaments were typical, and which was disseminated from the Siberian Minusinsk Depression and the Altai region through the Baraba Steppe and North-Kazakhstan to the Trans-Ural region (Fig. 1). Moreover, it is notable that the archaeological findings in these kurgans are dated not earlier then the10^th^ century AD, i.e. after the Hungarian conquest of the Carpathian Basin. The Hungarian conquerors from Karos cemetery appearing on these phylogenetic trees could represent the first generation of conquering populations based on their grave material, therefore identical mitogenome sequences can point out close biological connections or common source population of the Uyelgi population and the Hungarian conquerors.

The D4j phylogenetic tree contains one interesting phenomenon: the mitochondrial lineage of the sample Uyelgi21 from the Kurgan 11 located in the oldest horizon of Uyelgi cemetery clusters only with one modern-day Hungarians, whose lineage is ancestral to the lineage of Uylegi21. The findings of this Kurgan 11 (belonging to the Srostki culture) show similarities to the typical findings of the Hungarian conquerors from the Carpathian Basin as well (see Fig. 2 and Supplementary Fig. S1h).

The mitogenome of individual Uyelgi10 and three identical lineages of two Hungarian conquerors (Hconq1 and Hconq6) from Balatonújlak-Erdő-dűlő and Hconq9 from Makó-Igási járandó cemetery clustered together in one branch on the phylogenetic tree of haplogroup U5a1a1 (Supplementary Fig. S4q). The Uyelgi10 from Kurgan 7 of the middle horizon of the cemetery shows mixed character from archaeological point of view: the findings can be connected to the 9^th^ century AD as well as to the cultural influences of the Srostki culture (for the detailed information see Supplementary information)^29,30^. The samples of adult women from Balatonújlak-Erdő-dűlő buried with gilt silver hairpins could be dated (based on archaeological findings) to the middle third of the 10^th^ century AD^31^. One of their burials had a grave with a sidewall niche of eastern origin. The grave from Makó-Igási járandó without findings is dated to the middle third of 11^th^ century AD, i.e. to the Árpádian Age, when conquerors and the local population presumably admixed already. Interestingly, the 25-30 years old man shows some Asian cranial traits as the most men buried in this cemetery^32^.

The connection of Uyelgi cemetery and Hungarian conquerors is visible on the N1a1a1a1a branch of the tree of haplogroup N1a1 too, that was prevalent among the ancient Hungarians (Fig. 5). Here seven Hungarian conqueror samples from cemeteries Kenézlő-Fazekaszug, Orosháza-Görbicstanya and Karos-Eperjesszög clustered together on one branch, while the five Uyelgi samples from the earliest and latest horizons are located together next to this branch. These results signalize indirect connection between these two populations and don’t speak for their direct successiveness but rather for their common source in agreement with the archaeological chronology of Uylegi site.

The maternal genetic connection of the Cis-Ural region and the Hungarian conquerors is apparent especially on the phylogenetic tree of mitochondrial haplogroup T2b4h, where Bartym2, Bay3 and Hungarian conqueror from Karos site^14^ are located on the same branch, moreover, the individuals from Bartym and Karos share the same lineage that is ancestral to the mtDNA lineages of individual from Bayanovo (Supplementary Fig. S4k). The lineage of Karos (K1/3286) sample was determined as of possibly Asian origin by Neparáczki et al.^14^, nevertheless, their assumption is revisited by our data, not only by actual phylogenetic connections but due to the recurrent western presence of eastern lineages even from pre-medieval times. The burial of this adult male in Karos was without findings because disturbance of the Karos I cemetery’s burials by agricultural activity.

### Ancient paternal lineages of the South-Ural region

Majority of Uyelgi males belonged to Y chromosome haplogroup N, and according to combined STR, SNP and Network analyses they belong to the same subclade within N-M46 (also known as N-tat and N1a1-M46 in ISOGG 14.255). N-M46 nowadays is a geographically widely distributed paternal lineage from East of Siberia to Scandinavia^33^. One of its subclades is N-Z1936 (also known as N3a4 and N1a1a1a1a2 in ISOGG 14.255), which is prominent among Uralic speaking populations, probably originated from the Ural region as well and mainly distributed from the West of Ural Mountains to Scandinavia (Finland). Seven samples of Uyelgi site most probably belong to N-Y24365 (also known as N-B545 and N1a1a1a1a2a1c2 in ISOGG 14.255) under N-Z1936, a specific subclade that can be found almost exclusively in todays’ Tatarstan, Bashkortostan and Hungary^17^ (ISOGG, Yfull).

Median Joining (MJ) network analysis is performed using 238 N-M46 Y-haplotypes including seven samples from Uyelgi detected with 17 STR loci (Fig. 6, Supplementary Table S8) as well as 335 N-M46 Y-haplotypes with 12 STR loci (Supplementary Fig. S12, Supplementary Table S8). Based on MJ of 17 Y-STR loci, certain samples show identical or one-step neighbour profiles to Bashkirs, Khantys^17^, Hungarians^34^, Tatars from Volga-Ural region and a Central Russian sample^17^ (Fig. 6). The MJ based on 12 Y-STR data show one-step neighbour connection of Uylegi with two Hungarian conquerors from Bodrogszerdahely-Bálványhegy and Karos-Eperjesszög^15^ (Supplementary Fig. S12). YHRD online database show further affinities or identities among Finnish, Ural region (Sverdlovsk Oblast) or European Russian region (Penza and Arkhangelsk Oblasts) samples, notably either from territories of Uralic language affinities or along the supposed migration route of early Hungarians. It is noteworthy that the seventh-century Avar elite from the Carpathian Basin^35^, in spite of the similar N-M46 frequency to Uyelgi, had a distant subtype (N-F4205, N1a1a1a1a3a in ISOGG 14.255), which is prominent in present-day Mongolic speaking populations around Lake Baikal^33^. Furthermore they had a fairly different population history than populations of this study, therefore they shall not be confused with each other^35^.

Uyelgi11 from Kurgan 29 belongs to J2 Y-haplogroup. The Y-haplogroup J is widespread nowadays descended from the Near East^36^. Interestingly, a Hungarian conqueror from Sárrétudvari-Hízóföld (SH/81) carries the J2a1a subgroup^16^, however Uyelgi11 could not be typed downstream to J2 and therefore further assumptions cannot be made at this level.

Uyelgi4 belongs to G-L1266 (G2a2b2a1a1a1b in ISOGG 14.255), which sublineage is confirmed to be present outside of Europe within the European G-L140 branch of G. Among Hungarian conquerors the presence of G-L30 (G2a2b in ISOGG 14.255) was attested by Neparáczki et al.^16^ from Karos II (K2/33) without further classification or STR data, but recently G-L1266 is confirmed by Fóthi et al.^15^ which sample could also be included in our STR analysis. By using 14 STR markers in this case, due to the limitations of the database, MJ network shows a Caucasian affinity of both Hungarian conqueror (RP/2) and Uyelgi individuals (Supplementary Fig. S11, Supplementary Table S9), however, neither identity nor monophyly can be observed between them.

Both studies^15,16^ indicate Caucasian origin for part of the Hungarian conquerors based on the prevalence of this specific G2a Y-haplogroup. This hypothesis cannot be confidently excluded by our data nor our network analysis, however its presence in Uyelgi site could reshape this theory in the future.

In the Cis-Ural sample set the DNA preservation was insufficient for proper paternal lineage analyses, the only obtained N-M46 Y-haplotype of Bay2 sample and the R1b haplotype of Bartym3 do not have direct matches in the worldwide YHRD database, however, we found four one-step-neighbours of Bay2 from Sverdlovsk Oblast (Ural region) and Lithuania.

## Conclusions

The Ural region had an important role in ancient Hungarians’ ethnogenesis based on archaeological, linguistic and historical sources, although the results of these research fields exhibit differences of chronological and cultural aspects. The here presented new mitogenome, Y-chromosome and shallow shotgun autosomal DNA sequence data from the South-Urals confirms the region’s relevance from population genetic perspective too.

The overall maternal makeup of the investigated 36 samples from the Ural region in a phylogenetic and phylogeographic point of view suggests a mixed characteristic of rather western and rather eastern components, although the paternal lineages are more homogenous with Y-haplogroups typical for the Volga-Ural region. The exact assignment of each mitochondrial haplotype of the Trans-Uralic Uyelgi population to the eastern and western Eurasian components is impossible, but comprehensive representatives are present. Mitochondrial haplogroups of European origin N1a1a1a1a and H40b provide a horizon-through success of maternal lineages with inner diversification, which suggests a base population of a rather western characteristics. On the other hand, identical (C4a1a6) or single (A, A12a, C4a2a1) haplotypes with strong eastern phylogeography, highly pronounced in the third horizon, suggest a relatively recent admixture to this population. The apparent co-occurrence of genetic and archaeological shift is however contradicted by the homogeneity of ancestry components, nuclear genomic PCA positions, homogeneity of paternal makeup (although this one itself can be explained by patrilocality), and presence of eastern component (C4a1a6) in all horizons. Despite the fact that the genetic contribution of a population related to the Srostki culture cannot be excluded at this level, it is more likely that the majority of eastern components admixed before the usage of the Uyelgi cemetery. The uniparental genetic composition of Uyelgi population signals them as a chronologically and/or geographically related population to the possible genetic source of the Hungarian conquerors. Furthermore, their preliminary autosomal results show that they shared their allele frequency makeup with modern Uralic and West Siberian populations that are linguistically or historically related to Hungarians, which provide a good standpoint for future studies.

The maternal phylogenetic connections of Uyelgi with Hungarian conquerors can be divided to indirect (monophyletic but not successive) and direct (identical or one-step neighbour) relationships. Interestingly, indirect connections can be genetically assigned to the western-characteristic base population, whereas direct connections are almost exclusive to the admixed eastern component. One possible explanation for this phenomenon is that Hungarian conquerors and Uyelgi shared common ancestry in the past that separated prior eastern admixture, latter which provided genetic components subsequently to both groups. The exact origin or identification of the eastern component yet to be described, however, nuclear admixture proportions and loose phylogenetic connections points towards Central Asia, but further and deeper analyses with extended dataset is required for firm our statements.

The phylogenetic makeup of Cis-Ural region questions their compactness or successiveness; however, the scarce data does not allow extensive analysis for this group. Hungarian conqueror connections here are sporadic, but regional affinity is observable, which is more pronounced in MDS and PCA. Earlier studies based solely on the genetic makeup of Hungarian conquerors tend to connect the non-European lineages to various eastern regions, but especially the presence of rare Far East haplotypes in the Late Iron Age and Early Medieval Cis-Ural group may reshape these conclusions in the future.

## Material and Methods

### Sampling

Our aim was to collect samples from all available anthropologically well characterised human remains from five cemeteries of the Ural region: from Uyelgi 22 samples, from Bayanovo (Boyanovo) three samples, from Sukhoy Log five samples, from Bartym five samples and from Brody one sample, as well as nine comparative samples from Carpathian Basin (for more information see Supplementary Table S1).

Owing to the pressure of insufficient amount of samples from each cemetery, aside from the large chronological difference between cemeteries of Bayanovo, Sukhoy Log, Bartym and Brody, individuals deriving from there were grouped as “Cis-Ural” in the mtDNA population genetic analyses, indicated by the relative geographical proximity (~400 km) and archaeological similarities (see Supplementary text). Furthermore, these cemeteries are connected to Hungarian prehistory through various archaeological evidences and historical sources as well^1,3^.

### Sample preparation

All procedures leading to Next Generation Sequencing of entire mitochondrial DNA were performed in a dedicated ancient DNA laboratory according cleanness recommendations at Laboratory of Archaeogenetics, Institute of Archaeology, Research Centre for the Humanities in Budapest, Hungary. After photo documentation and bleach, samples were UV-C irritated for 30 minutes per side. Therefore, samples were abraded by using bench-top sandblaster machine with clean sand, followed by additional UV-C exposure procedure for 20 minutes per side. Cleaned bone samples were grinded into fine powder (types of bone samples are given in Supplementary Table S1). Approximately 100 mg (80 – 120 mg) of powder was collected and processed^13,37^.

### DNA Extraction, library preparation and NGS sequencing

DNA extraction was performed according the protocol of Dabney et al.^38^ with minor changes pointed also by Lipson et al.^37^

For verifying the result of DNA extraction, a test PCR reaction was performed^37^. DNA library preparation with partial uracil-DNA-glycosylate treatment was performed as described at Rohland et al. ^39^ with minor modifications. Partially double-stranded and barcoded P5 and P7 adapters were used for T4 ligation reaction. Each DNA extract was assigned unique barcode combination. No barcode combination was used more than once in one batch. After fill-in reaction, 13.2 μL of product was amplified using TwistAMP (TwistDX) in 34.3 μL final volume. Amplification reaction products were purified by AMPure Beads Purification (Agilent).

To capture the target sequences covering whole mitochondrial genome and autosomal SNPs, in solution hybridisation method was used as described by Csáky et al.^35^, Haak et al.^40^ and Lipson et al.^37^. Captured samples as well as raw libraries for shotgun sequencing were indexed using universal iP5 and unique iP7 indexes^41^.

Next generation sequencing was performed on Illumina MiSeq System (Illumina) using V3 (2 × 75 cycles) sequencing kits and custom sequencing setup.

### Pre-processing of the Illumina sequence data

Customized in-house analytic pipeline was run on the Illumina sequence data. Paired reads were merged together with SeqPrep master (John JS. SeqPrep. https://github.com/jstjohn/SeqPrep), requiring an overlap at least 10 base pairs for capture, and 5 base pairs for shotgun data. For one mismatch, the one with higher base quality was accepted, the overlapping reads with two or more mismatches were discarded. Cutadapt^42^ were used to remove barcodes as well as to discard fragments too short (<15 bp for shotgun and <20 bp for capture) or/and without barcode. The pre-processed reads were mapped to the reference sequence (GRCh37) using BWA v.0.7.5^43^, with MAPQ of 20, and gap extension of 3 base pairs. These permissive options were considered due to the frequent occurrence of low quality and/or amount of reliable fragments in the data pool. Samtools v.1.3.1^44^ were utilized for further data processing, such as indexing or removing PCR duplications. BAM files uploaded to the ENA repository contain both single and paired end reads. Damage pattern estimations were performed by MapDamage v.2.0.6 (https://ginolhac.github.io/mapDamage/).

BAM files imported into Geneious 8.1.7 (https://www.geneious.com/) were re-assembled against either rCRS and RSRS using 5 iteration steps. The automatic variant caller of Geneious was used with a minimum variant frequency of 0.8 and minimum coverage of 3× to collect SNPs to a database. In this step, the known troublesome sites (309.1C(C), 315.1C, AC indels at 515-522, 16182C, 16183C, 16193.1C(C) and 16519) were masked. Remaining ambiguous sites were inspected by eye. Consensus FASTA files were created by Geneious 8.1.7 software (https://www.geneious.com/). Mitochondrial haplogroup determinations were performed in HaploGrep^45^, which utilizes Phylothree mtDNA tree build 17 (https://www.phylotree.org/). The Y-haplogroup were assigned based on Y-STR data using nevgen.org, as well as based on Y-SNP capture and shallow shotgun sequencing data by Y-leaf v1 and v2^46^. Terminal Y-SNPs were verified on the Y tree of ISOGG version 15.34 (https://isogg.org/tree/).

### Estimates of contamination

The contamMix 1.0.10 was used to estimate the level of human DNA contamination in the mitochondrial DNA^40,47^. All of our samples show 99%< endogenous content, which makes them eligible for whole genome analyses. For the results see Supplementary Table S2.

### Population genetic analyses

The different size of populations used in sequence-based analyses is caused by absence of whole mitogenomes of some populations.

Standard statistical methods were used for calculating genetic distances between investigated populations from Ural region (Uyelgi and Cis-Ural) and 26 ancient and 43 modern populations. Even the Uyelgi population was composed of sample-pools from two distinct sampling events with approximately one hundred years between the collected samples, it was considered together in further analyses. Nine samples of conquerors from Carpathian Basin were excluded from any population analyses because of the possible sample bias due to selected haplogroups. The whole mitochondrial genome alignment of the samples were performed in SeaView by ClustalO^48^ with default options, and later regions with poor alignment quality were discarded. Population pairwise F_ST_ values were calculated based on 4015 modern-day and 1132 ancient whole mitochondrial sequences using Arlequin 3.5.2.2^49^. The Tamura and Nei substitution model was used^50^ with gamma value of 0.62, 10,000 permutations and significance level of 0.05 in case of comparison between two investigated populations from Ural region and 43 modern-day Eurasian populations (for the references see Supplementary Table S6). For the comparison of 28 ancient populations the F_ST_ calculation was performed with Tamura and Nei DNA evolution model with gamma value of 0.599, 10,000 permutations and significance level of 0.05. The genetic distances of linearized Slatkin F_ST_ values^51^ were used for multidimensional scaling (MDS) and visualized on a two-dimensional plot (Supplementary Fig. S9a and Fig. S10) using metaMDS function based on Euclidean distances implemented in the vegan library of R 3.4.1^52^.

Spearman rank correlation matrix of FST values was calculated in Pandas (Python) and visualized in seaborn package by clustermap function using Euclidean metric.

Principal component analysis was performed based on mtDNA haplogroup frequencies of 64 modern and 50 ancient populations. 32 mitochondrial haplogroups were considered in PCA of ancient populations, while in PCA of modern populations and two ancient populations from Ural region we considered 36 mitochondrial haplogroups (Supplementary Tables S3 and S4). The PCAs were carried out using the prcomp function in R 3.4.1 and visualised in a two-dimensional plot with first two (PC1 and PC2) or the first and third principal components (PC1 and PC3) (Fig. 3b, Supplementary Figs. S5 and S7).

For hierarchical clustering, Ward type algorithm^53^ and Euclidean measurement was conducted based on haplogroup frequencies of ancient and modern populations as well, and displayed as a dendrogram in R3.4.1 (Supplementary Figs S6 and S8). The same population-pool was used as in PCAs.

Shallow shotgun and captured nuclear DNA sequences were 1bp trimmed on both ends by trimBam function of bamUtil (https://genome.sph.umich.edu/wiki/BamUtil:_trimBam).

Genotypes from shotgun data were called for the Human Origin SNP panel by samtools mpileup command (-q30 and –Q30) and by pileupCaller (which is designed to sample alleles from low coverage sequence data, see https://github.com/stschiff/sequenceTools). Prior to the ADMIXTURE analysis, we filtered for missing SNPs in the dataset (“--geno 0.999 parameter”) and pruned SNPs in strong linkage disequilibrium with each other using the parameters “--indep-pairwise 200 25 0.4” in PLINK^54^, leaving 1,146,167 SNPs. We run unsupervised ADMIXTURE with K=16 on a “1240k” worldwide dataset (https://reich.hms.harvard.edu/datasets) of published ancient captured/shotgun sequenced and modern deep sequenced genomes^55^. The plotted samples’ sources are seen in Supplementary Table S10.

### Phylogenetic and network analysis

All available mitochondrial genome sequences in NCBI (more than 33,500) were downloaded and sorted according to their haplogroup assignments. Then multiple alignments for each haplogroup were performed with ClustalO within SeaView^48^. Neighbour Joining (NJ) trees were generated by PHYLIP version 3.6^56^. The phylogenetic trees then were drawn by Figtree version 1.4.2 (http://tree.bio.ed.ac.uk/software/figtree). We decided to omit median joining network (MJN) to avoid unresolvable ties and bootstrap calculation due to the low number of substitutions.

To analyse the Y-STR variation within the Y chromosomal haplogroups N1a1-M46 and G2a, Median Joining (MJ) networks were constructed using the Network 5.0 software (http://www.fluxus-engineering.com). For the N1a1-M46 Y-haplogroup MJ Network calculation with 17 STR loci 238 samples, and for MJ network calculation with 12 STR loci of the same haplogroup, 335 samples of 27 ancient and modern population were included (Supplementary tables S8). The MJ network analysis of G2a Y-haplogroup was calculated based on 14 STR data using 120 samples of 27 populations (Supplementary tables S9). Post processing MP calculation was used, creating network containing all shortest tree. Repeats of the locus DYS389I were subtracted from the DYS389II.

## Acknowledgements

The reported study was carried out with the financial support of the Russian Foundation for Basic Research in the framework of project No. 18-59-23002: “The origins of the formation of the culture of ancient Hungarians. Archaeological paleoanthropological and paleogenetic aspect of the study of medieval monuments of the Southern Urals and Western Siberia”, furthermore by RFBR and FRLC research project No. 19-59-23006: „The problem of cultural transformations of Magyars on the way of Hungarian Conquest”.

We thank Sufija Renatovna Gazizova from the South Ural State University (national research university) in Cheljabinsk and Aleksej Vladimirovich Parunin from Community foundation “South-Ural” Cheljabinsk for providing archaeological information about the Uyelgi cemetery and bone materials. Furthermore, we are grateful to Olga Evgenevna Poshekhonova from the Tyumen Scientific Centre SB RAS (Institute of the problems of Northern development), Elizaveta M. Chernykh from the Department of History, Archaeology and Ethnology of Udmurtia of the Institute of History and Sociology at the Udmurt State University Izhevsk, Andrey M. Belavin from the Perm State Humanitarian-Pedagogical University for the archaeological data and providing bone material from Cis-Ural region for DNA analyses.

We thank Viktor Szinyei for preparing the base maps. We are grateful to István Major from the Isotopte Climatology and Environmental Research Centre, Institute for Nuclear Research, Debrecen, Hungary for performing the ^14^C data of samples within the project GINOP-2.3.2-15-2016-00009 ‘ICER’.

## Author contributions

B.G.M., A.T. and A.Sz-N. designed the study. V.Cs., D.G., B.Sz., B.E. and. B.S. performed the ancient DNA analyses. V.Cs., D.G. and A.Sz-N. performed population genetic and phylogenetic analyses. S.G.B., I.V.G., N.P.M., A.S.Z., A.V.S., R.D.G., A.V.D and A.T. performed the archaeological evaluation, provided the historical background and interpretation. V.Cs., D.G., B.Sz., B.G.M, T.A. and A. Sz-N. wrote the paper. All authors read and discussed the manuscript.

## Data availability

The NGS data were uploaded to the repository ENA (European Nucleotide Archive) under project number PRJEB39054 and are available upon the publication.

