## SUPPLEMENTARY INFORMATION for "Early Medieval Genetic Data from Ural Region Evaluated in the Light of Archaeological Evidence of Ancient Hungarians"

### 1. Studied archaeological cultures, sites and findings

#### The Uyelgi cemetery and the Kushnarenkovo-Karayakupovo culture (Trans-Ural region)

Uyelgi cemetery is located on the eastern side of the Ural Mountains (Trans-Ural region) 2.5 km far from Kanzafarova (Chelyabinsk region, Kunashaksky district) in Bashkortostan (Central Russia) and is dated to the 9–11th centuries. The excavation of the late Kushnarenkovo culture site is still ongoing leading by Sergej G. Botalov and the archaeologists of the South-Ural Federal University. The location of the cemetery situated on the west site of Uyelgi lake and surrounded with several lakes (e.g. Saygerli) provided good environmental conditions to settle, especially for the peoples dealing with livestock farming. Based on the mostly scattered artefact findings of the Uyelgi cemetery, it emerged from the Ural-region's archaeological sites as the cemetery with the largest number of parallels to the Hungarian Conquest period findings in Carpathian basin.

People buried their deceased at Uyelgi in pit graves under or between the kurgans. Unfortunately, 80% of the graves were disturbed in various level. So far, 20 kurgans and nearly 100 objects have been investigated primarily on the northeast and northwest site of the cemetery.

Sampling was carried out in the earlier discovered 10–11th century area of the cemetery and also in the 9th century section of the site that has been explored in 2015. The earliest artefact horizon of the cemetery appeared in the kurgans no. 10 and 32 located on the flat and deeper part, in the middle part between the northern and southern rise. A child´s grave and tombs of harnessed horse-burial and of an archer buried with silver belt were excavated in the kurgan no. 32. Two thoroughly disturbed burials are known from kurgan no. 10. On the basis of typochronological analysis, the finds of this horizon can be dated to the 9th century (end of 8th century): flat, smooth surfaced silver mounts sometimes with ribs or hemisphere-shaped decoration. A very archaic archery equipment typical of the 7–8th centuries was unearthed as well, which refers to the oldest time horizon of the cemetery that can be dated to the turn of the 8–9th centuries. This assumption is strengthened by the radiocarbon analysis, as it resulted in the period between 770 and 900 AD (with 95,4% probability)[1]. The younger part of the cemetery is located on a northern elevation in the northern area of the site, where the plant ornaments and niello are characteristic for the findings. Graves of the 10–11th centuries can be distinguished from the 9th-century burials on the basis of the typochronological differences of their metal finds: the appearance of plant ornament, gilt background and developed variants of double crescent-shaped mounts, although this research topic requires further research. A transition phase between 9th and 10–11th centuries was identified in the Uyelgi cemetery. Here belongs the child's grave no. 7 of the kurgan no. 9 which includes finds both from the 9th and the 10th centuries. The grave no. 5 from the kurgan no. 7 can be dated unambiguously to the 10–11th centuries according to the typochronological and radiocarbon analyses (879–1150 cal AD) as well[1], although, findings typical for the 9th century were also found in this kurgan. This phenomenon implies the usage of these kurgans in the later periods too. The typical burials of Uyelgi cemetery were found in the kurgans 1, 2, 3, 28, 29, 30, 34a, which cannot be dated earlier than 10th century[2].

The findings from the 9th century of the Uyelgi cemetery belong to the late Kushnarenkovo culture, which indicates the autochthon component in this complex population. The Gaultry cemetery and several cemeteries of Bashkortostan from 9th century (investigated by N. A. Mazhitov) show similar characteristics as well. The experts named these as "Ural component" because of the local ornament and moulding of the belts, which differs from the Srostkinsky culture from the Altai/Minusinsk region and from the Perm style of the west foreground of Ural Mountains which styles are the closest to the ancient Hungarians' culture. Their co-appearance attests to the mixed character of the site, but the other hand, the affinity of the 10th century Hungarian Conquerors needs further investigation and explanation.
One sample (Grave 5 of Kurgan 11) of this earliest horizon (720-941 cal AD) should be mentioned separately. The archaeological findings of Srostkinsky types and the secondary filling of the kurgan dated this tomb to the middle and second half of the 9th century AD. The further archaeological features, as the shallow tomb, bone buckle and gilt belt end, date the grave to the 10-11th century AD. Based on this fact, the archaeologists suppose that in this tomb the earliest newcomer of eastern origin was buried, which signals the earlier appearance of this new population from the East dated to the second half of the 9th century AD.
The large DNA sample-set from Uyelgi cemetery is justified not only by the Russian-Hungarian joint research (started in 2013) but also by the closest analogy known from Uyelgi to the burials of the Hungarian Conquest period in the Carpathian basin. The cemetery might also contribute to the investigation of those ancient Hungarians who did not participate in the westward migration but remained in the Trans-Ural.

**The Nevolino culture: Brody, Bartym and Sukhoy Log cemeteries (Cis-Ural region, Lower Kama-valley)**

The Nevolino culture was located in the Kama valley at the western foothills of the Ural Mountains and represents the most significant and well-researched culture of the 4–9th centuries history of the region. Its end was previously associated with the migration of Hungarians, therefore we investigated samples from all of its three chronological phases in accordance with archaeological chronology. The Nevolino culture (6-9th centuries) is one of the well-studied Prikamye cultures. It was investigated mainly by the archaeological expedition of the Udmurt State University (Izhevsk) under the guidance of R. Goldina[3] and occupied the Sylva basin from its start till the Tis inflow (the Sylva is a left-bank tributary of the Chusovaya the tributary of Kama). The length of the territory is over 150 km from the north to the south and over 100 km from the west to the east. Well-known cemeteries belonged to the Nevolino culture are near to the villages Nevolino, Brody, Bartym, Verkh-Saya and Sukhoy Log. Its key geographical location between the forests and the steppe, branched river system that connected the region with the south (by the Belaya River), north, west, south-west (by the Kama) and with the east to the Trans-Urals (by the Sylva and Chusovaya), helped the population to develop in a successful and dynamic way. Constant contacts with close neighbours and remote regions – Central Asia, Sasanian Iran, Byzantine,

Baltic Sea Region and others, from which the human greatest achievements came – allow the Prikamye population to establish its own expressive, distinctive and original culture.

The Brody cemetery of the Nevolino culture can be dated to the 4–5$^{th}$ centuries AD, the Bartym cemetery to the 6–7$^{th}$ centuries AD, and Sukhoy Log cemetery after which the late phase of the culture is named, is dated to the 8–9$^{th}$ centuries AD. The Nevolino culture occupied about 15,000 square km in the southern zone of the forest area. As a result of long-distance trade activity of the population, there are clear archaeological connections with the east of the Urals, the Sasanian Iran, Byzantium, Central Asia and the Baltic Sea Region. The center of the area's archaeological research is operated by the Department of Archaeology of the Udmurt State University in Izhevsk[3–5].

**The Bayanovo cemetery and the Lomovatovo culture (Cis-Ural)**

The Lomovatovo culture was situated in the western outskirts of the Ural Mountains, northeast of the area of the Nevolino culture in the Kama valley, and can be dated to the 8–10$^{th}$ centuries AD.

The Bayanovo cemetery located in Dobrjansky region (Perm region) next to Bayanovo village is not only the most significant site of the culture, but due to its rich Ugric archaeological heritage, it is also the most important archaeological site of the western territory of the Urals. Although the site has been discovered already in 1951-1953 leaded by V. A. Oborin, its intensive research began in 2005 under the guidance of the Department of History at the Perm State Humanitarian-Pedagogical University with leading by Andrey. V. Danich. The cemetery is dated from the 9$^{th}$ to the beginning of the 11$^{th}$ century as it is proven by the radiocarbon analysis as well[6].

So far, 421 graves have been excavated on the 2413 $m^2$ explored area, which resulted in the most important finding of the south variant of Lomovatovo culture.

In the investigated grave no. 277 was found a skeleton of 18-25 years old man, the bones were in anatomically position, in the north part of the grave was a mandible of fragmented skull and the long bones of lower and upper extremities. The north part of the child-grave no. 280 (10-12 years old girl) in anatomically position contains skull slightly leaned to the right, long bones and fragmented ribs and pelvic. The skeleton lain in extended supine position in northeast direction. In case of the grave no. 365, only a fragments of skull, 14 teeth and remains of vertebrae were excavated. Based on the vertebrae, she was a 20-25 years old women.

Unfortunately, the early medieval inheritance of this region is determined by the poorly preserved bones because of the bad soil conditions. The graves of the Lomovatovo culture and Bayanovo site with findings of the death-mask, women´s jewels and silver ornaments with gold background are archaeologically related with the Conquest period Hungarians in many ways.

**Investigated cemeteries of the Hungarian Conquest period from the Carpathian Basin**

The majority of the investigated burials and cemeteries from Carpathian Basin comes from the well-documented recent excavations of the last 20 years.

Cemetery *Harta-Freifelt* located in the middle of Carpathian Basin contains typical (classical) Hungarian Conqueror archaeological findings from the 10th century[7]. Interestingly, the individuals of 22 burials in this cemetery are maternally not related to each other based on previously study of HVRI region of the mtDNA[8].

The small cemetery *Balatonújlak-Erdő-dűlő* cemetery in the western part of the Carpathian Basin represents the characteristic finds from the 10th century as well[9]. Based on archaeological interpretation, the eastern genetic connection of the cemeteries Harta-Freifelt and Balatonújlak-Erdő-dűlő is reasonably expected.

The cemetery-complex *Karos-Eperjesszög II and III* in the northeast parts of the Carpathian Basin is the richest burial of Hungarian conquest period, where also the elite of the Conquest period are buried[10].

The south – southeast part of the Carpathian Basin is covered by cemetery *M43 Makó-Igási járandó*[11]. This cemetery contains besides 10th century findings also archaeological finds belonged to the 12th century.

**Fig. S1 Archaeological sites and artefacts of the studied cemeteries from Ural region.**

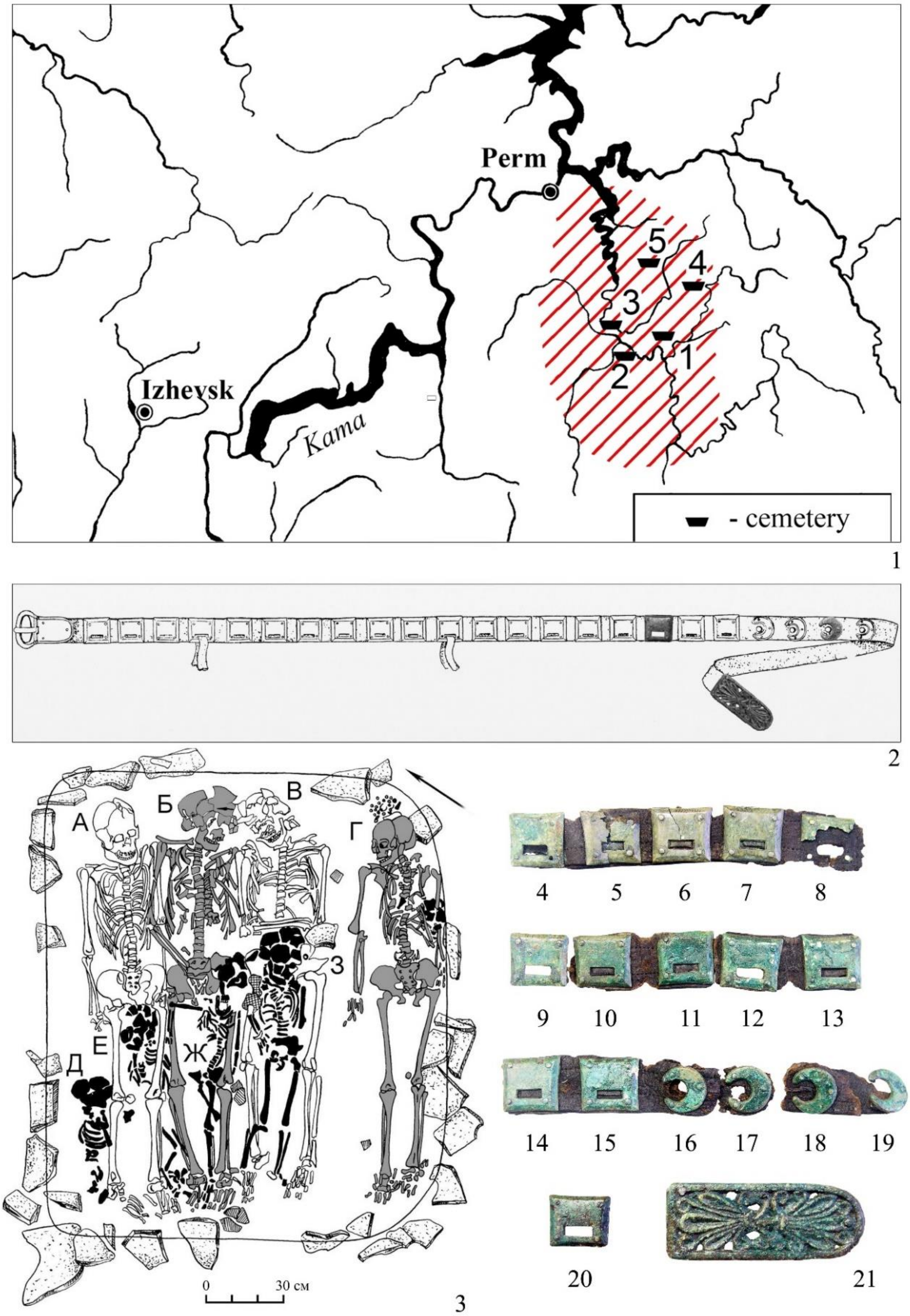

**Fig. S1a Location of cemeteries of Nevolino culture** (1: Sukhoy Log; 2: Nevolino; 3: Brody; 4: Bartym; 5: Verkh-Saya) (map by R. D. Goldina) (part 1). **The most typical finds and belt sets of the Nevolino culture** (part 2–21): Bartym cemetery, Grave 16 (illustrations by R. D. Goldina).

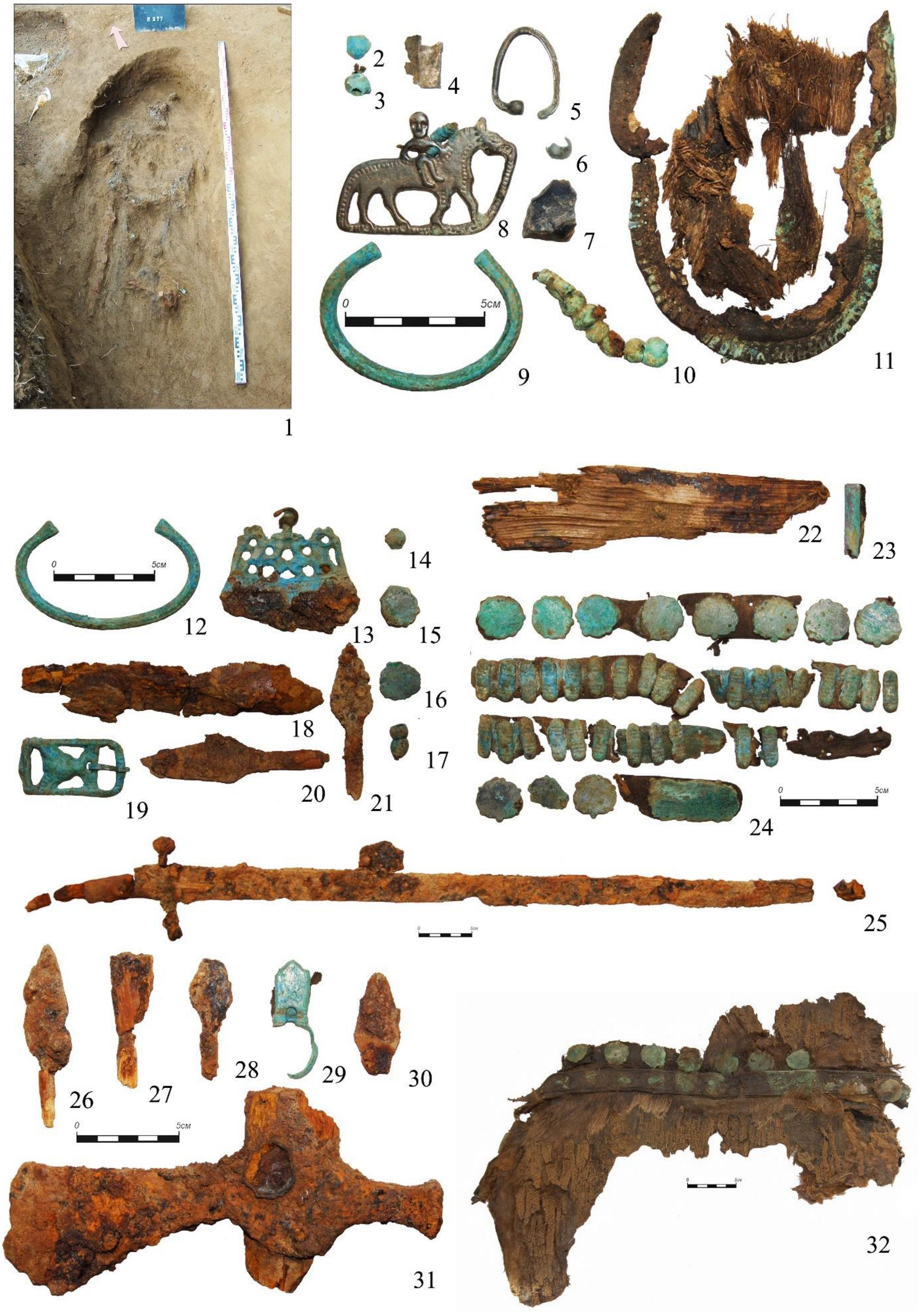

**Fig. S1b Bayanovo cemetery (Lomovatovo culture), Grave 277.** Photos were taken by A. V. Danich.

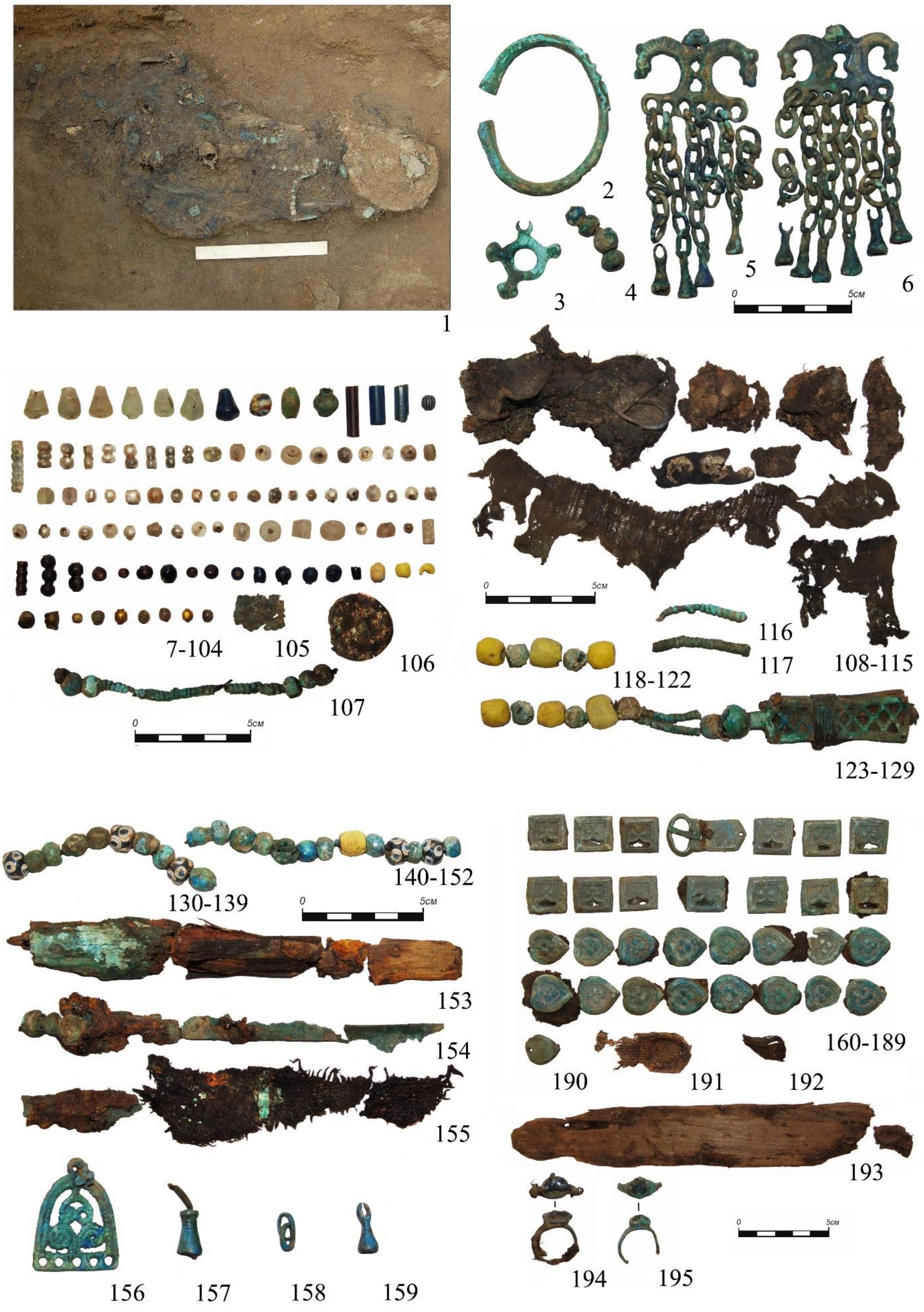

**Fig. S1c Bayanovo cemetery (Lomovatovo culture), Grave 280.** Photos were taken by A. V. Danich.

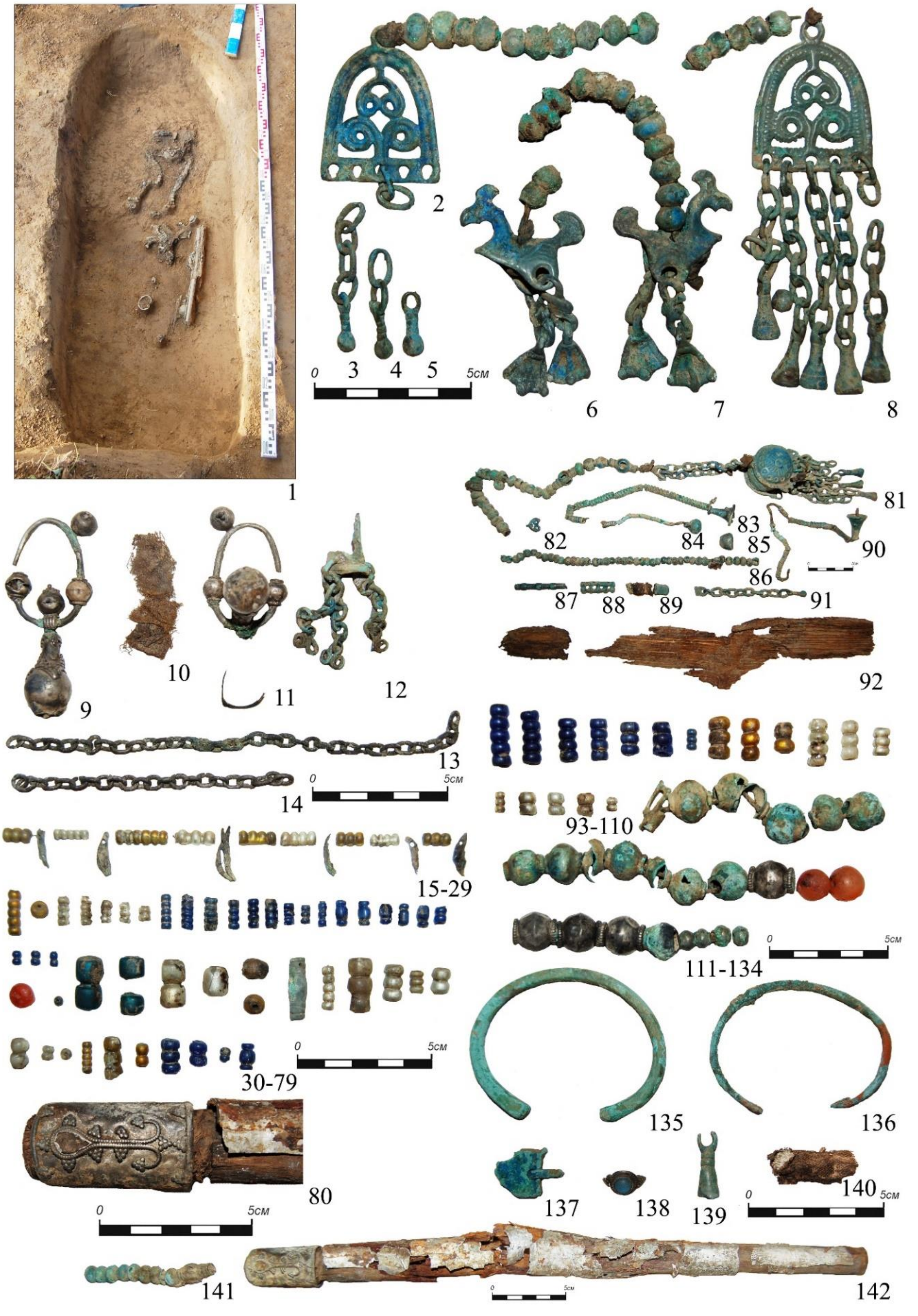

**Fig. S1d Bayanovo cemetery (Lomovatovo culture), Grave 365.** Photos were taken by A. V. Danich.

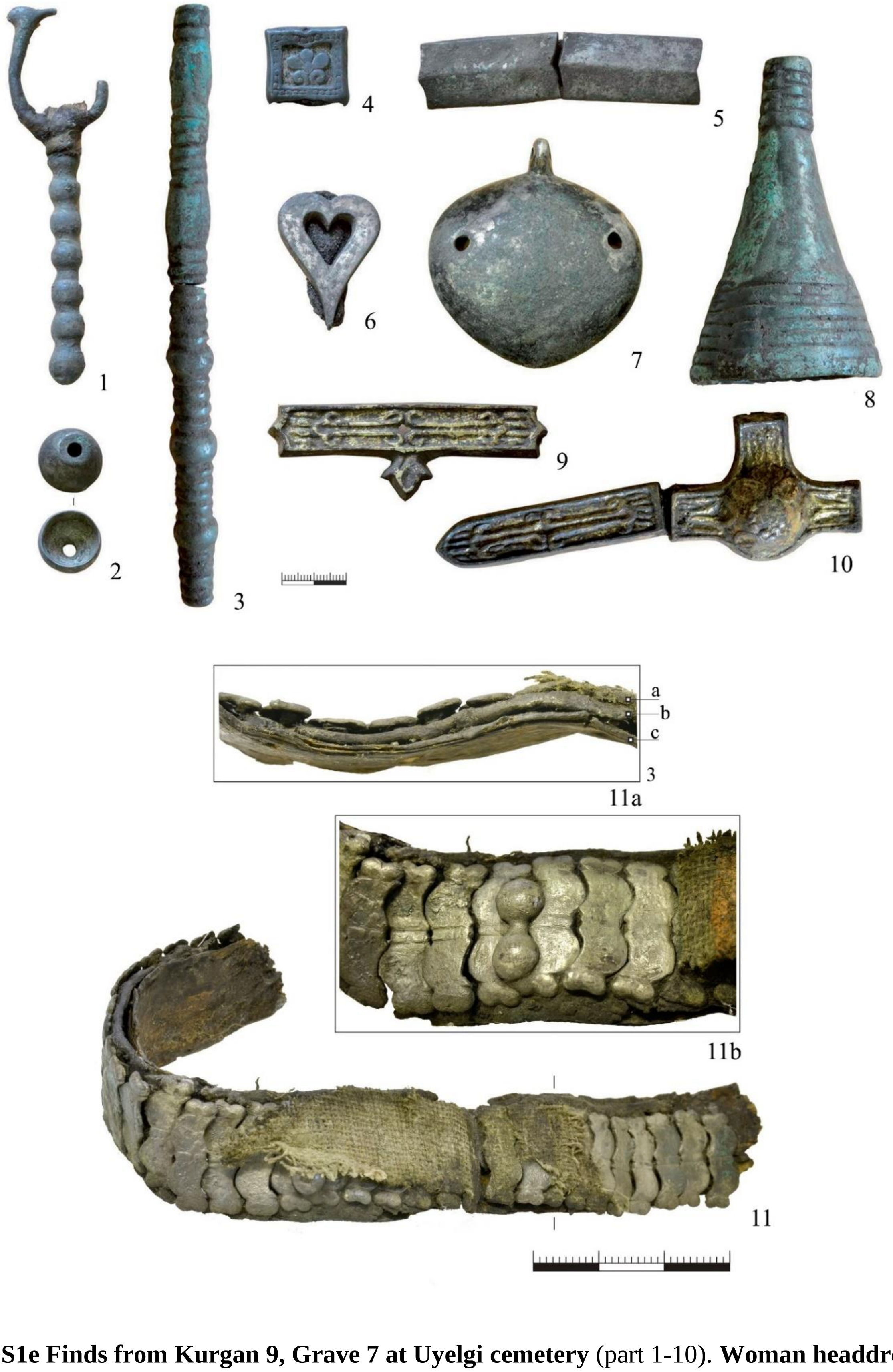

**Fig. S1e Finds from Kurgan 9, Grave 7 at Uyelgi cemetery** (part 1-10). **Woman headdress from Kurgan 32, Grave 1** (part 11; 11a: Cross section of the headdress – a: fittings; b: leather; c: birchbark; 11b: the central part of the headdress). Photos were taken by S. G. Botalov.

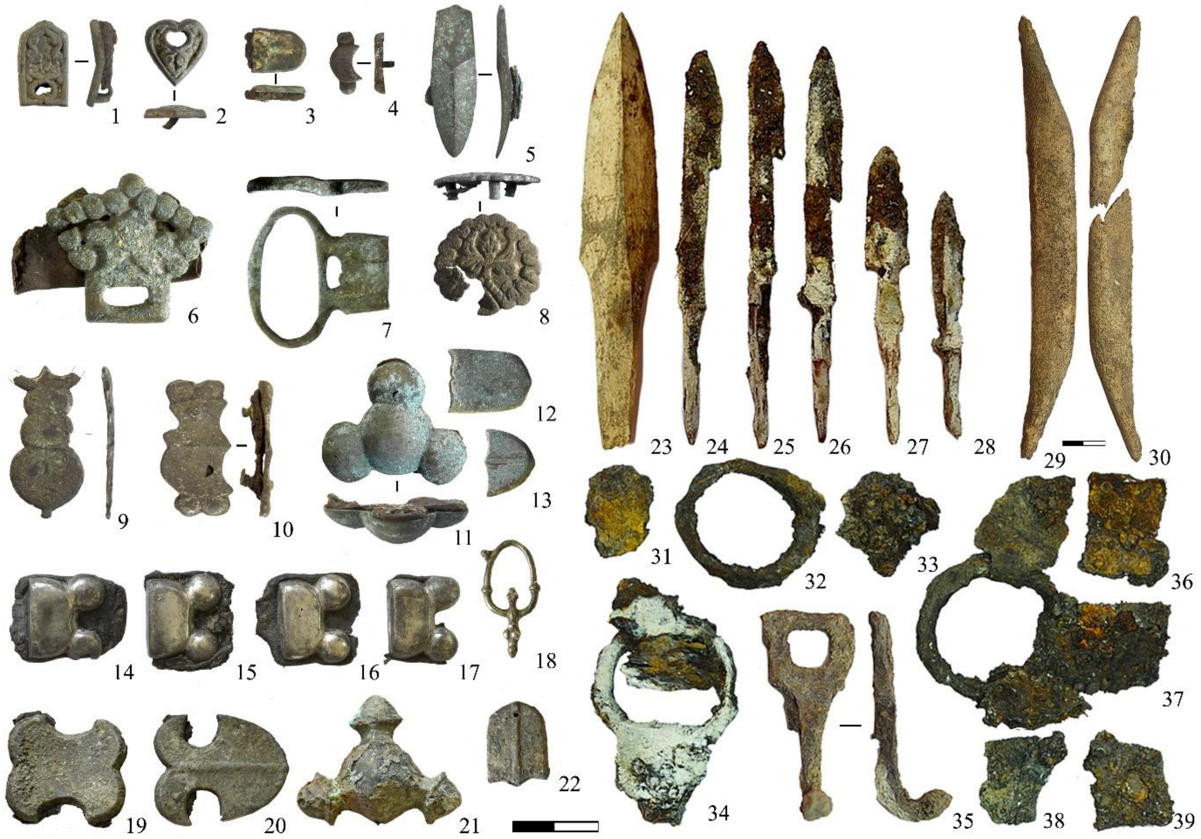

**Fig. S1f Finds from the early kurgans (9th century) of Uyelgi cemetery.** 1–9: Kurgan 32, embankment of kurgan; 10: Kurgan 32, Grave 12; 11–13: Kurgan 10, Grave 3; 14–18: Kurgan 32, Grave 14; 19–22: Kurgan 10, Grave 3; 23–39: Kurgan 32, Grave 16. Photos were taken by S. G. Botalov.

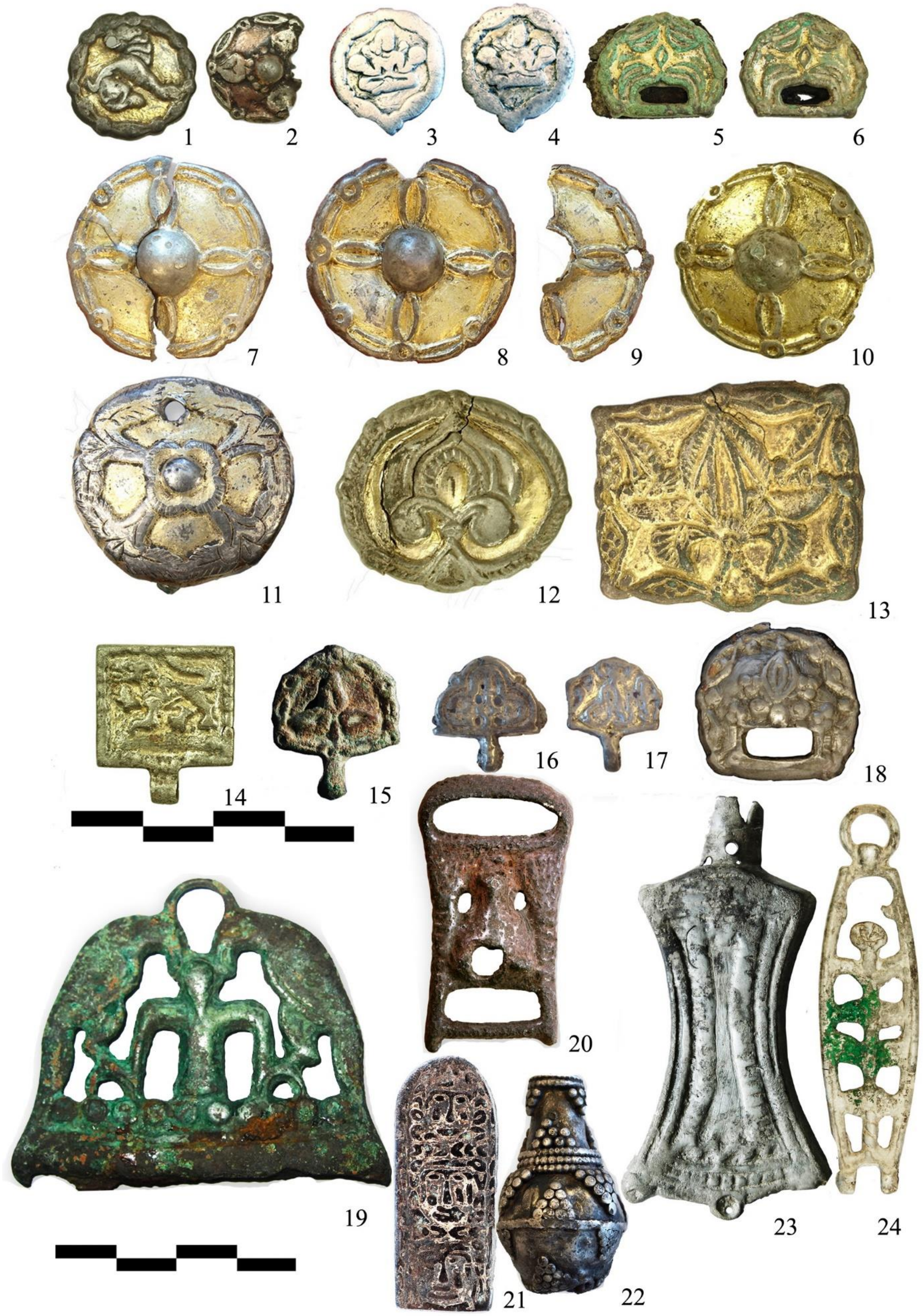

**Fig. S1g Findings from the Uyelgi cemetery.** Artefacts with ancient Hungarian character: 1-18; artefacts with Cis-Ural character: 19-24- Photos were taken by S. G. Botalov.

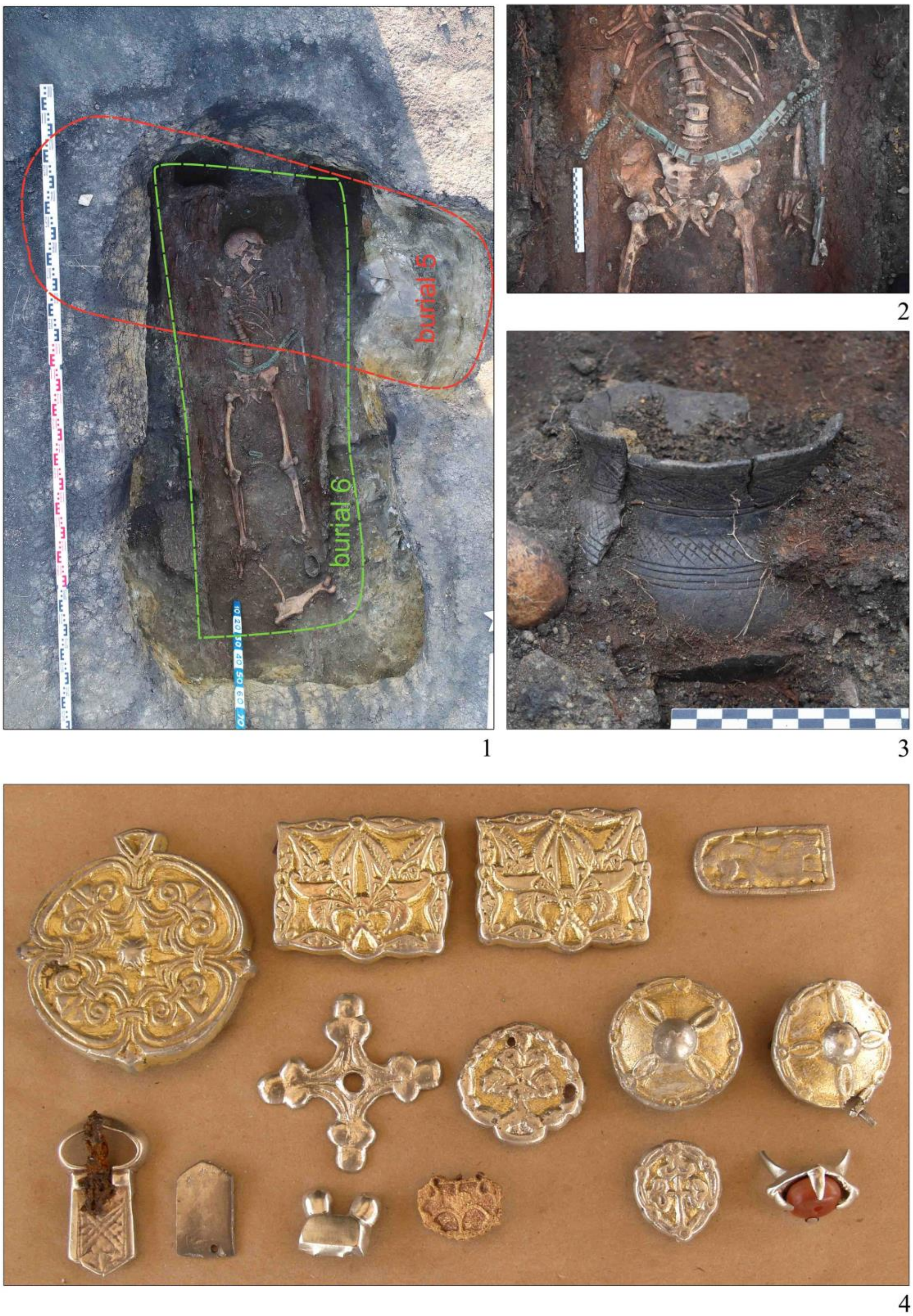

**Fig. S1h Uyelgi cemetery, Kurgan 11, Grave 5-6** (part 1-3). **Artefacts with ancient Hungarian characters from the excavation in 2019** (part 4). Photos were taken by S. G. Botalov.

#### 2. Radiocarbon dates and stable isotope data of the sampled burials

We report the 95.4% calibrated radiocarbon date confidence intervals from OxCal version 4.2 with the IntCal13 calibration curve[12,13]. The Uyelgi SPb and DeA radiocarbon samples were analysed by Grudochko et al.[1].

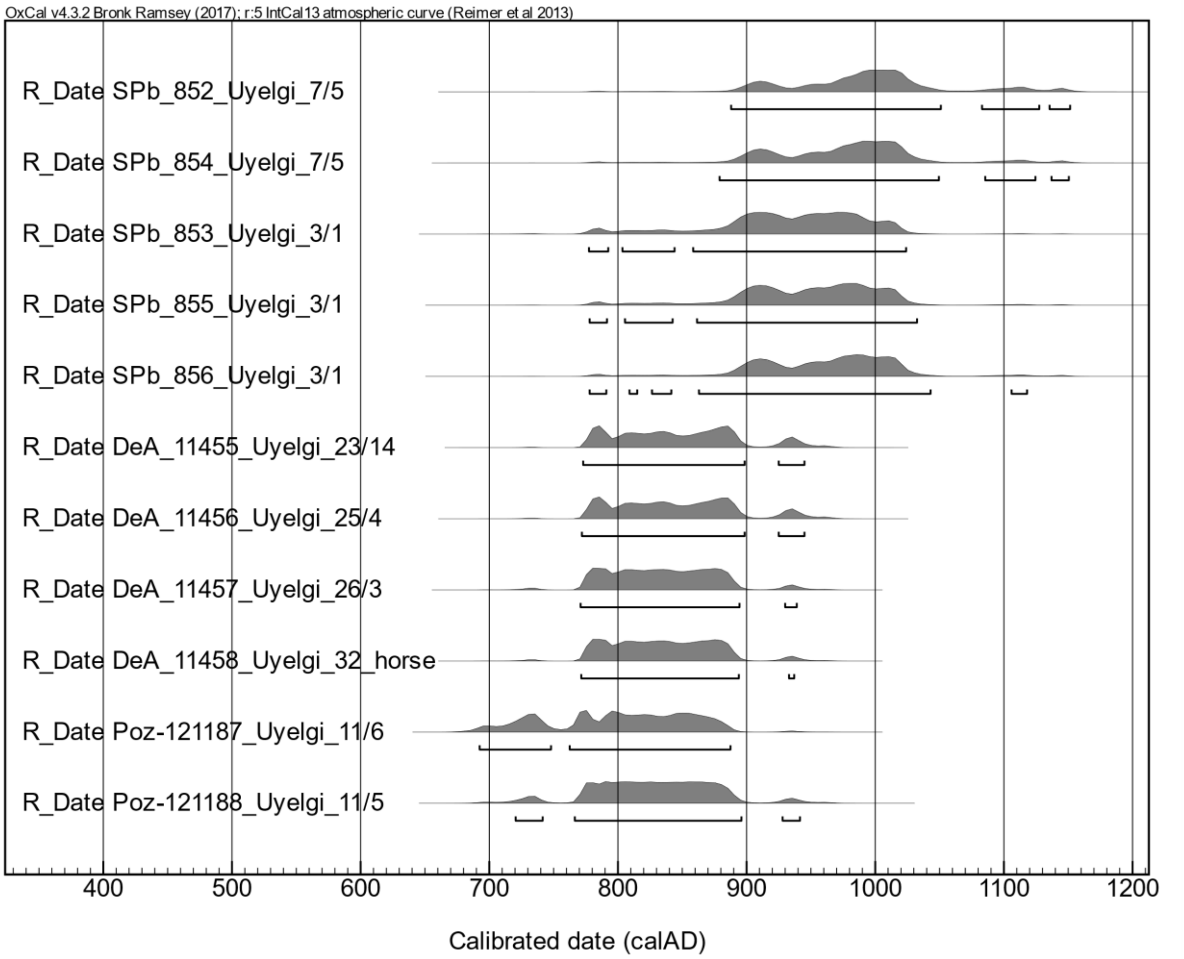

**Fig. S2a Calibrated radiocarbon dates of the samples from Uyelgi cemetery (Late Kushnarenkovo culture).** The dated samples signal well the early and middle/late horizons of this cemetery. The raw data are shown in SupplementaryTable S1.

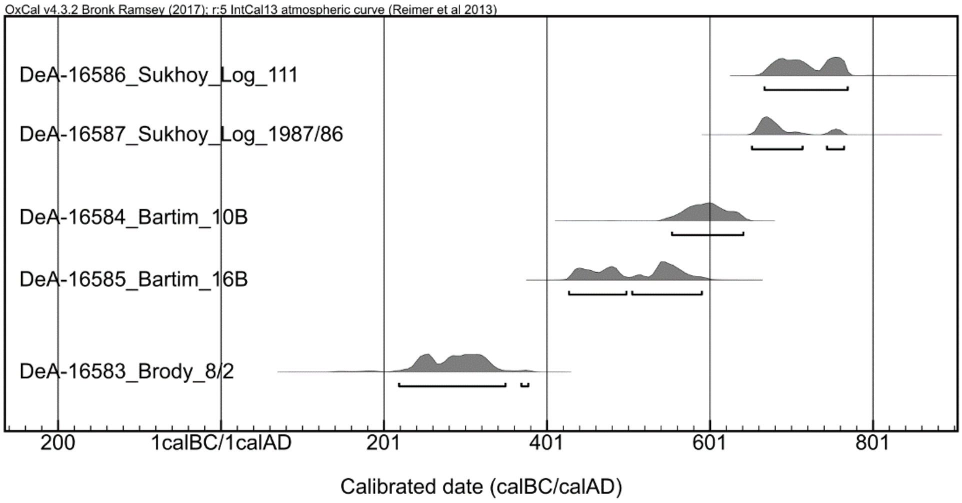

**Fig. S2b Calibrated radiocarbon dates of the samples of Nevolino culture:** Early Nevolino culture – Brody cemetery, Nevolino culture Phase II – Bartym cemetery and the Late Nevolino culture – Sukhoy Log cemetery. The raw data are shown in Supplementary Table S1.

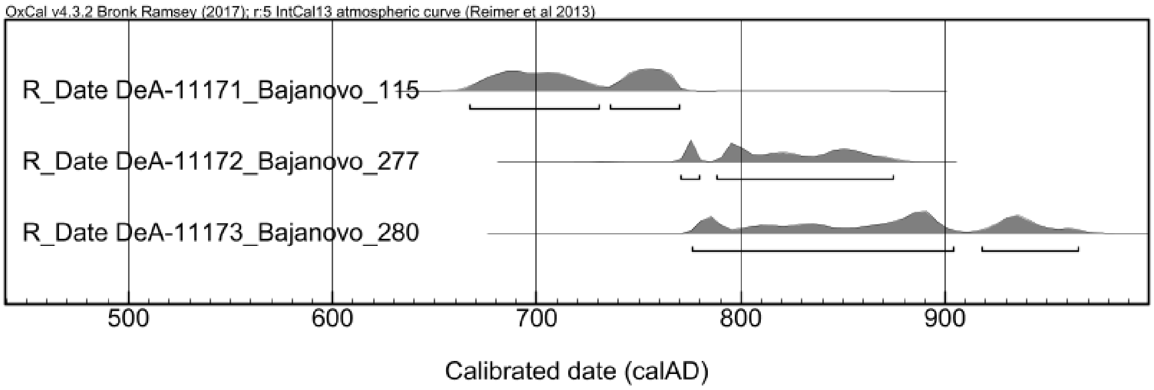

**Fig. S2c Calibrated radiocarbon dates of the samples of Late Lomovatovo culture:** Bayanovo cemetery. The sample Bayanovo Grave 115 from the earliest phase was not used for DNA analyses. The raw data are shown in SupplementaryTable S1.

| Sample Name | Nr lab. | $\delta^{13}C$ (‰) | $\delta^{15}N$ (‰) |
|---|---|---|---|
| Uyelgi21_ Uelgi ku11 gr5 KOL | Poz-0 | -21.3 | 11.7 |
| Uyelgi22_Uelgi ku11 gr6 KOL | Poz-0 | -20.1 | 10.3 |

**Supplementary Table S12.** Stable isotope data of two samples from the earliest horizon of the Uyelgi cemtery

### 3. Analyses of the shallow shotgun and captured genomic DNA data from Uyelgi cemetery

Summaries of SNP calling by pileupCaller:

| Sample Name | Total Called Sites | Non Missing Calls | average Raw Reads | average Damage Cleaned Reads | average Sampled From |
|---|---|---|---|---|---|
| Uyelgi1 | 598094 | 14180 | 2.479e-2 | 2.479e-2 | 2.479e-2 |
| Uyelgi2 | 598094 | 12418 | 2.176e-2 | 2.176e-2 | 2.176e-2 |
| Uyelgi4 | 598094 | 9032 | 1.586e-2 | 1.586e-2 | 1.586e-2 |
| Uyelgi5 | 598094 | 8980 | 1.549-2 | 1.549e-2 | 1.548e-2 |
| Uyelgi10 | 598094 | 10032 | 1.716e-2 | 1.716e-2 | 1.715e-2 |

**Supplementary Table S13a.** Genotype calling for the PCA analyses.

| Sample Name | Total Called Sites | Non Missing Calls | average Raw Reads | average Damage Cleaned Reads | average Sampled From |
|---|---|---|---|---|---|
| Uyelgi1 | 1233013 | 28748 | 2.428e-2 | 2.428e-2 | 2.427e-2 |
| Uyelgi2 | 1233013 | 26479 | 2.238e-2 | 2.238e-2 | 2.236-2 |
| Uyelgi4 | 1233013 | 19700 | 1.662e-2 | 1.662e-2 | 1.661e-2 |
| Uyelgi5 | 1233013 | 17832 | 1.4913e-2 | 1.491e-2 | 1.490e-2 |
| Uyelgi10 | 1233013 | 19943 | 1.660e-2 | 1.660e-2 | 1.658e-2 |

**Supplementary Table S13b.** Genotype calling for the ADMIXTURE analyses.

Only those samples were used for the PCA, that had over 8900 called SNPs. PCA was perform by smartpca (EIGENSOFT, https://github.com/DReichLab/EIG). The reference dataset was the published Human Origin datasets (https://reich.hms.harvard.edu/datasets) completed with Jeong et al.[14]. We calculated the principal components based on 1517 present-day west Eurasians and projected ancient individuals using lsqproject:YES and shrinkmode:YES. (The ancient populations and their references used for genomic PCA are listed in Supplementary Table S10.) PCA was plotted in conda with matplotlib.pyplot package of python.

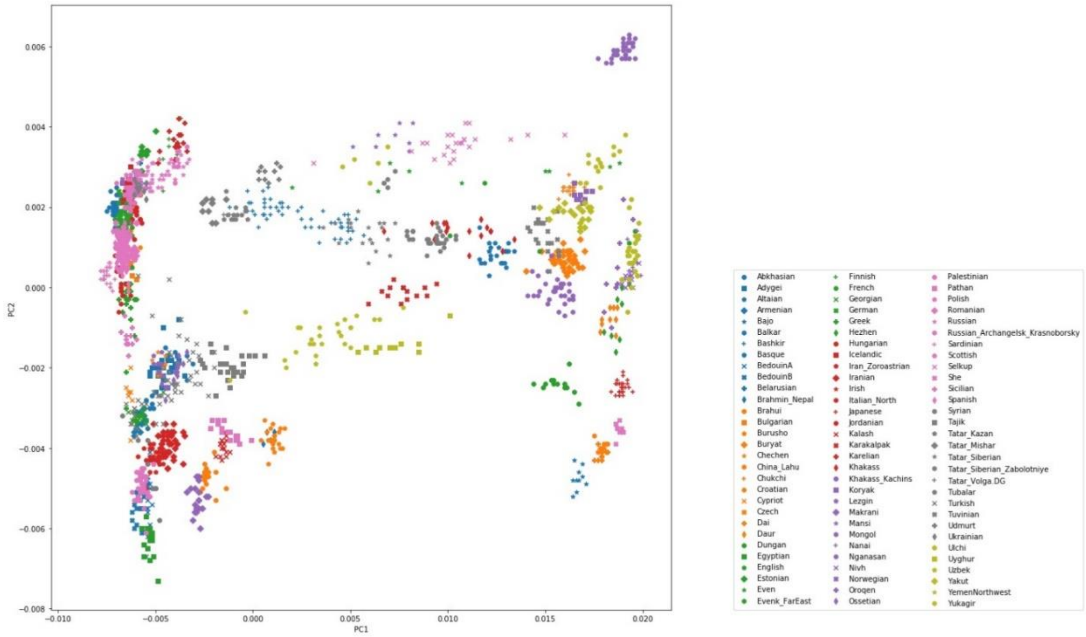

**Fig. S3a PCA with modern Eurasian individuals, used for the calculation of the principal components by smartpca.** Eigenvalues of PC1: 59.924, PC2: 8.044.

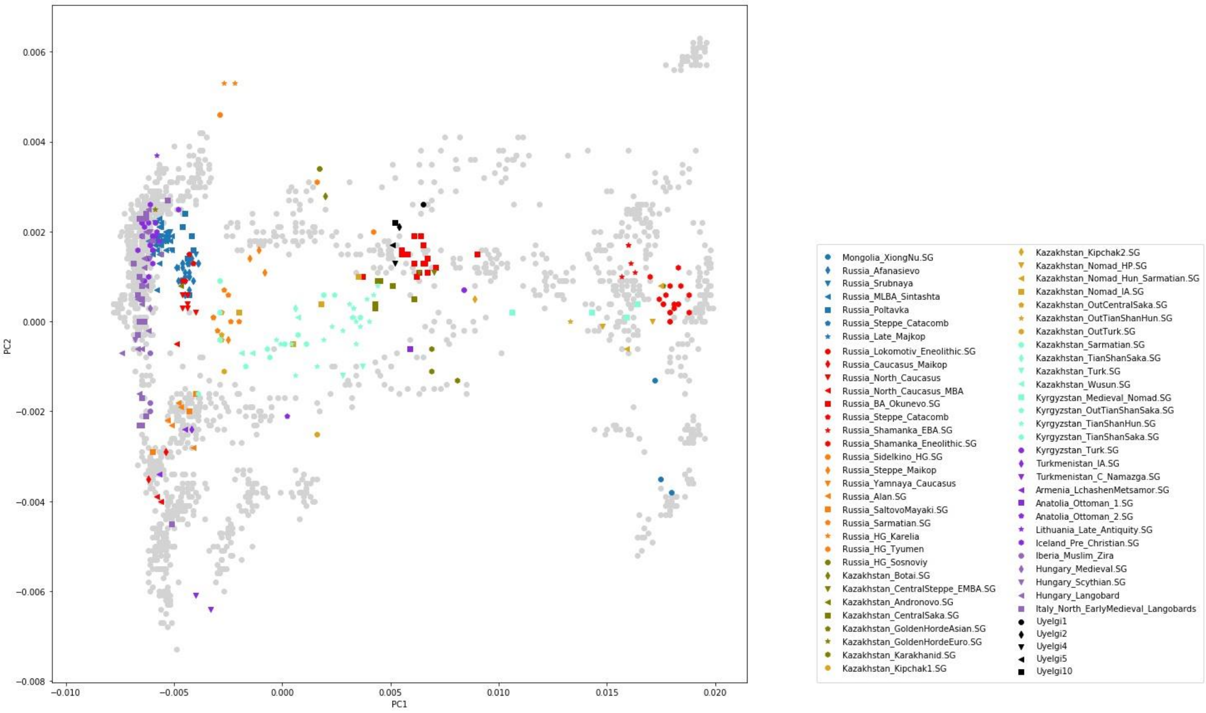

**Fig. S3b Projection of five ancient samples from Uyelgi site and other ancient samples to the modern individuals along PC1 and PC2.** PCs were computed by smartpca.

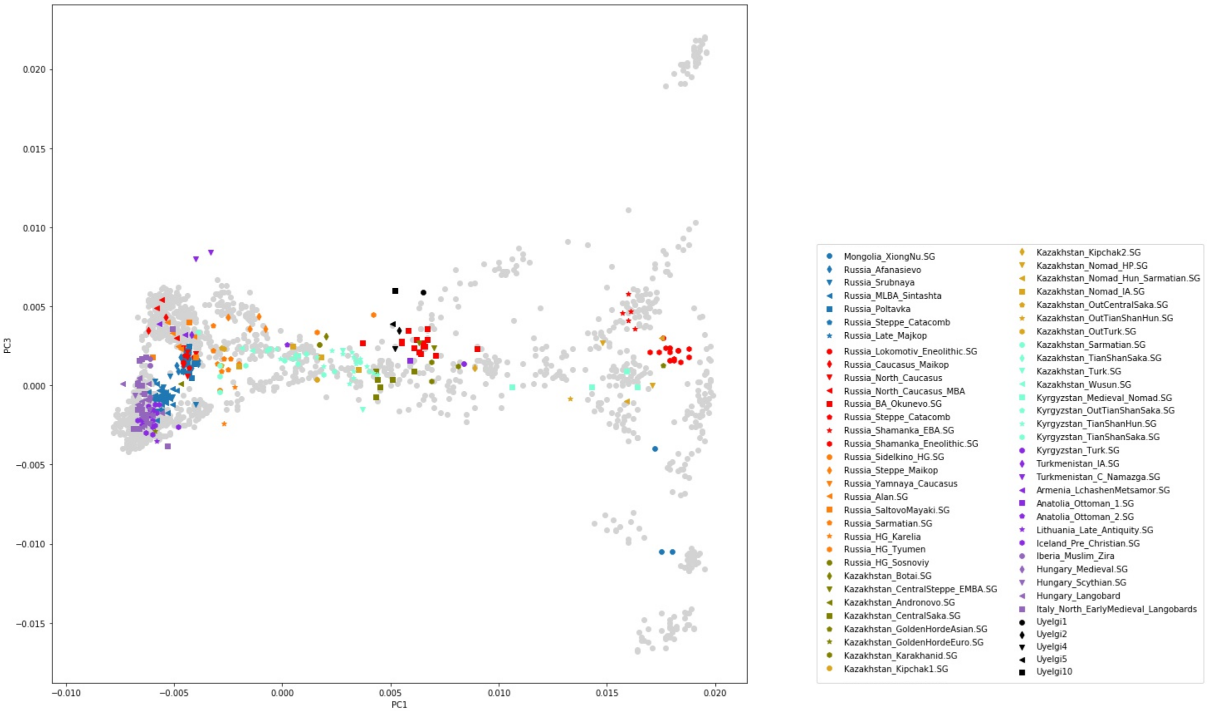

**Fig. S3c Projection of five ancient samples from Uyelgi site and other ancient samples to the modern individuals along PC1 and PC3.** PCs were computed by smartpca.

#### ADMIXTURE analysis

We can describe the affinities of ancestry components of Uylegi samples through those modern populations in that the five samples' ancestry components are maximized: Uyelgi population has modern European-related components, a modern North-Siberian-related, an East-Asian-related component and a component that is maximised in hunter gatherer samples from Tyumen and Afontova Gora3 Fig. S3d). A minor part of their ancestry components could be originated from the Caucasus/Central Asia based on modern Adygei, Pathan and Tajik genomes in the comparative study.

The five Uyelgi samples with an average of 22,540 SNPs show the most similar ancestry cluster proportions to present-day Mansis and Irtysh-Barabinsk Tatars[15]. The structure of ancestry components is different from Uylegi in case of the Okunevo that shows more North-Siberian and East-Asian related ancestry. Closest ancient proxies in the current published databases are the Iron Age Central Sakas and early medieval Kimak from present-day North Kazakhstan. The Central Sakas (Inner Asian Scythians) were modelled as a two-way mixture of Late Bronze Age pastoralists (56%) and southern Siberian hunter-gatherers (44%) by Damgaard et al.[16], which scenario is plausible for the Uyelgi population as well.

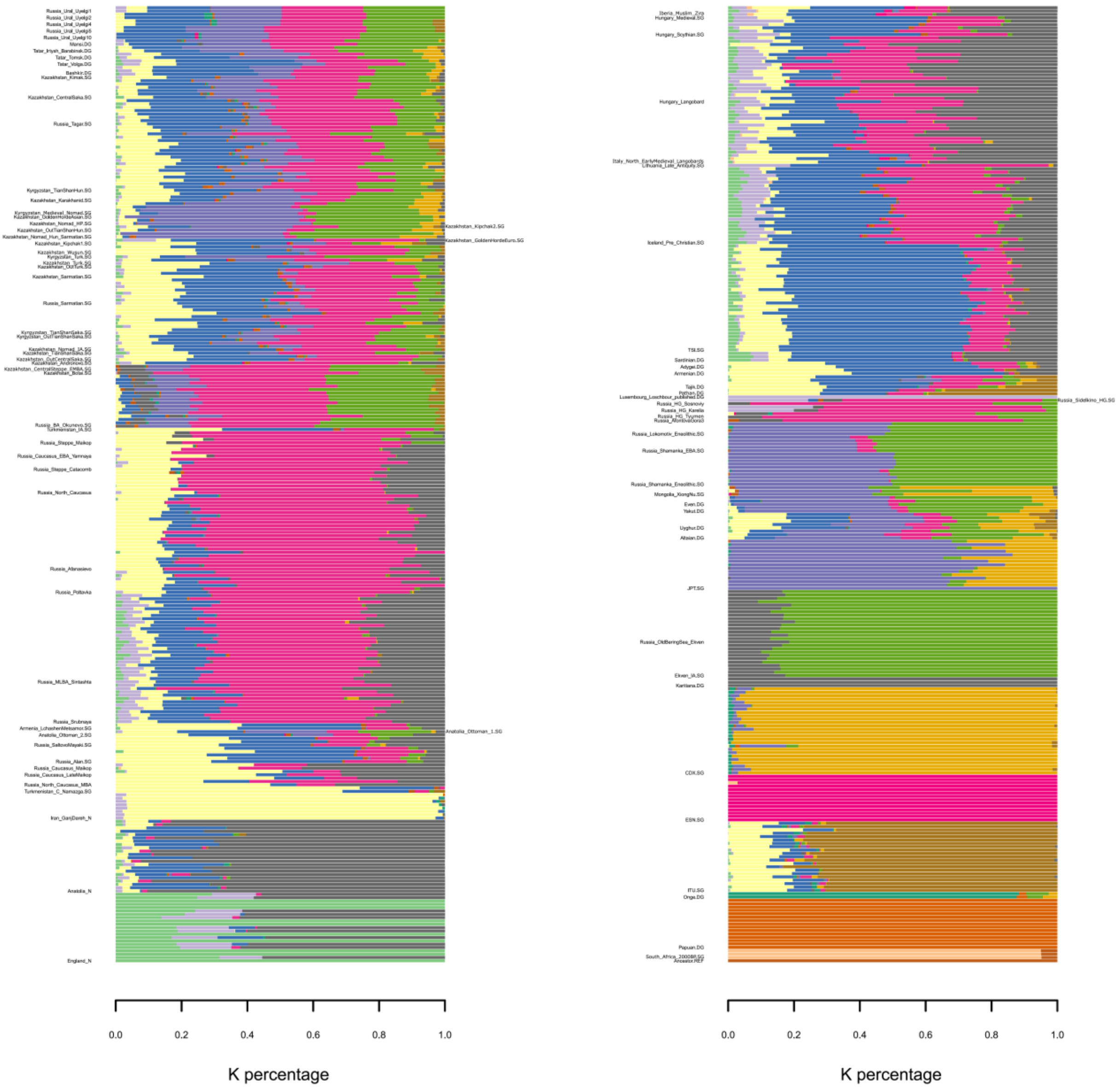

**Fig. S3d ADMIXTURE (K=16) barplot with 681 ancient and modern genomic data.** (TSI – Toscani in Italia; JPT – Japanese in Tokyo, Japan; CDX – Chinese Dai in Xishuangbanna, China; ESN – Esan in Nigeria; ITU – Indian Telugu in the UK)

#### 4. Mitochondrial phylogenetic trees and descriptions (Figures S4a-S4s)

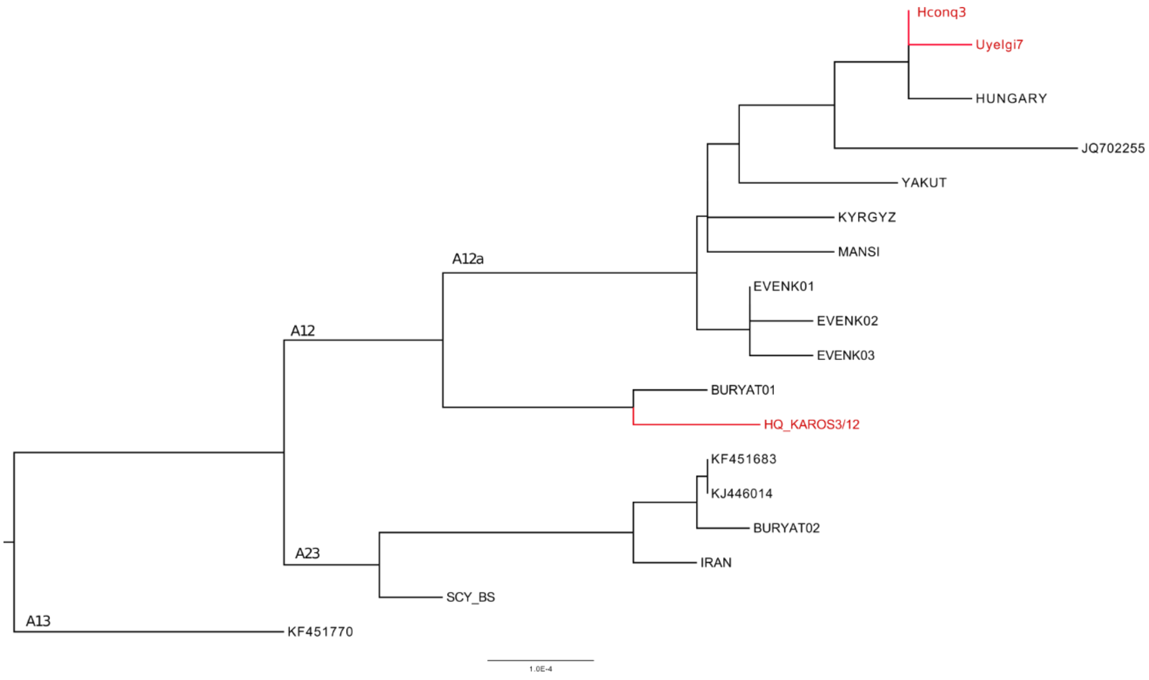

**Fig. S4a Phylogenetic tree of haplogroup A12a.**

The subhaplogroup A12a can be found in a Kurgan 30 grave (Uyelgi7) from the newest horizon of Uyelgi site (10-11th centuries), in a Hungarian Conqueror grave from Harta site, 10th century (Hconq3), and in a modern Hungarian sample from Debrecen region. In spite of the low sample coverage of this haplogroup the close proximity of the individuals in question without any outsider is apparent and presumes relatively close or direct maternal relationship between those samples (for the abbreviation and further information see Supplementary Table S7).

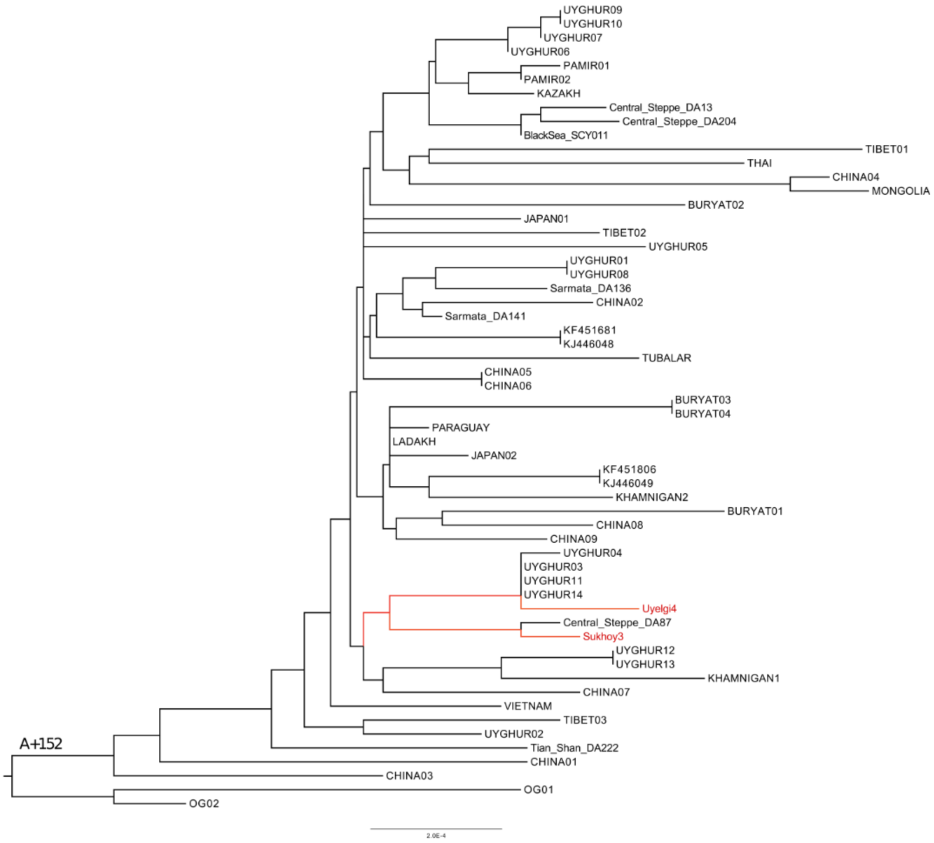

**Fig. S4b Phylogenetic tree of haplogroup A+152+16362.**

The subhaplogroup A+152+16362 can be found in a Kurgan 29 (grave 1,2) from the newest horizon of Uyelgi site, 10-11th centuries (Uyelgi4), as well as in sample from Grave 13 of Sukhoy Log cemetery (7-8th centuries) (Sukhoy3). The samples show modest proximities to each other. Within this branch, the Uyelgi4 groups together with Uyghur individuals, whereas Sukhoy3 is situated together with a medieval Kimak individual from Central Steppe[16] (for the abbreviation and further information see Supplementary Table S7). The overall structure of the phylogenetic tree suggests a Central-Eastern Asian (or at least Trans-Ural) origin and distribution of this lineage.

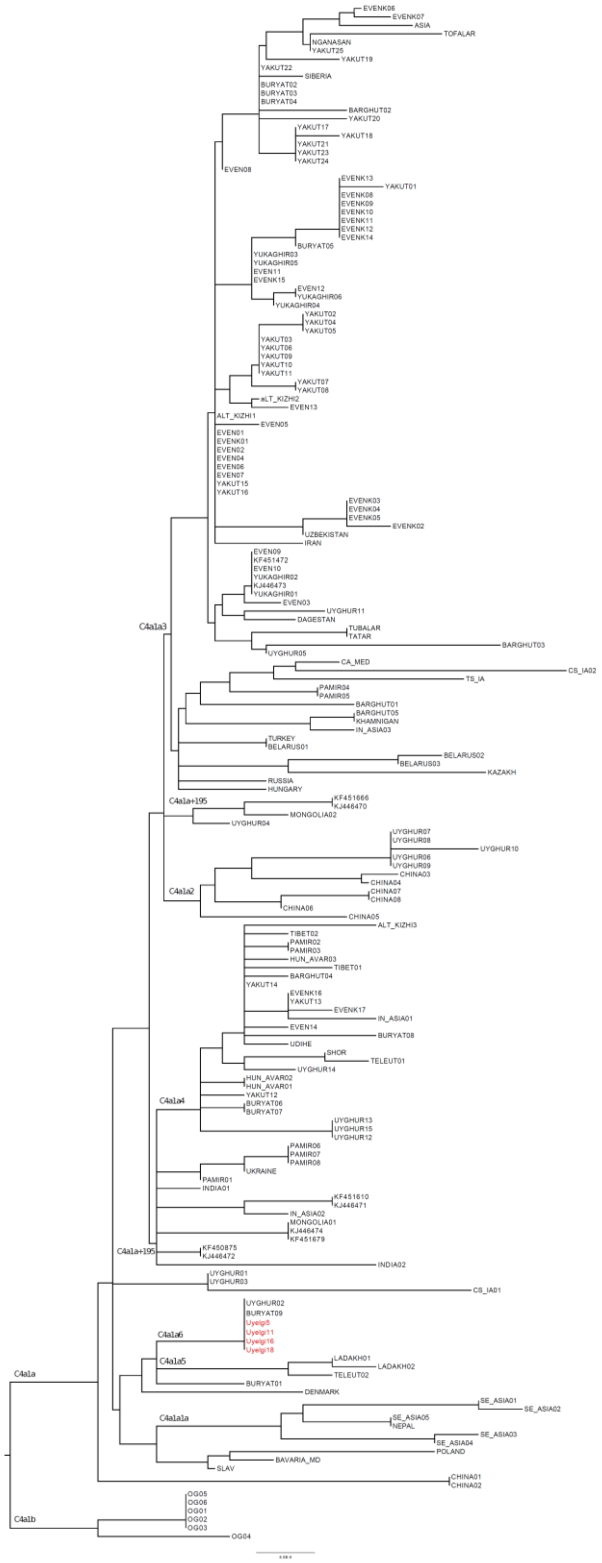

**Fig. S4c Phylogenetic tree of haplogroup C4a1a6.**

The subhaplogroup C4a1a6 can be found in four graves from all three horizons of Uyelgi site: in Kurgan 30 (Uyelgi5) and Kurgan 29 Grave 7 (Uyelgi11) from the newest horizon (10-11$^{th}$ centuries), in Kurgan 9 Grave 5 (Uyelgi16) from the middle horizon (9-10$^{th}$ centuries) and in Kurgan 32 Grave 1 (Uyelgi18) from the oldest horizon (9$^{th}$ century). These samples from

Uyelgi site are all identical presuming a close maternal relationship between them, apparently connecting the three horizons together. In addition, the Uyelgi individuals are identical to a Buryat an Uyghur sample as well, suggesting a more eastern or steppe origin for these samples (for the abbreviation and further information see Supplementary Table S7).

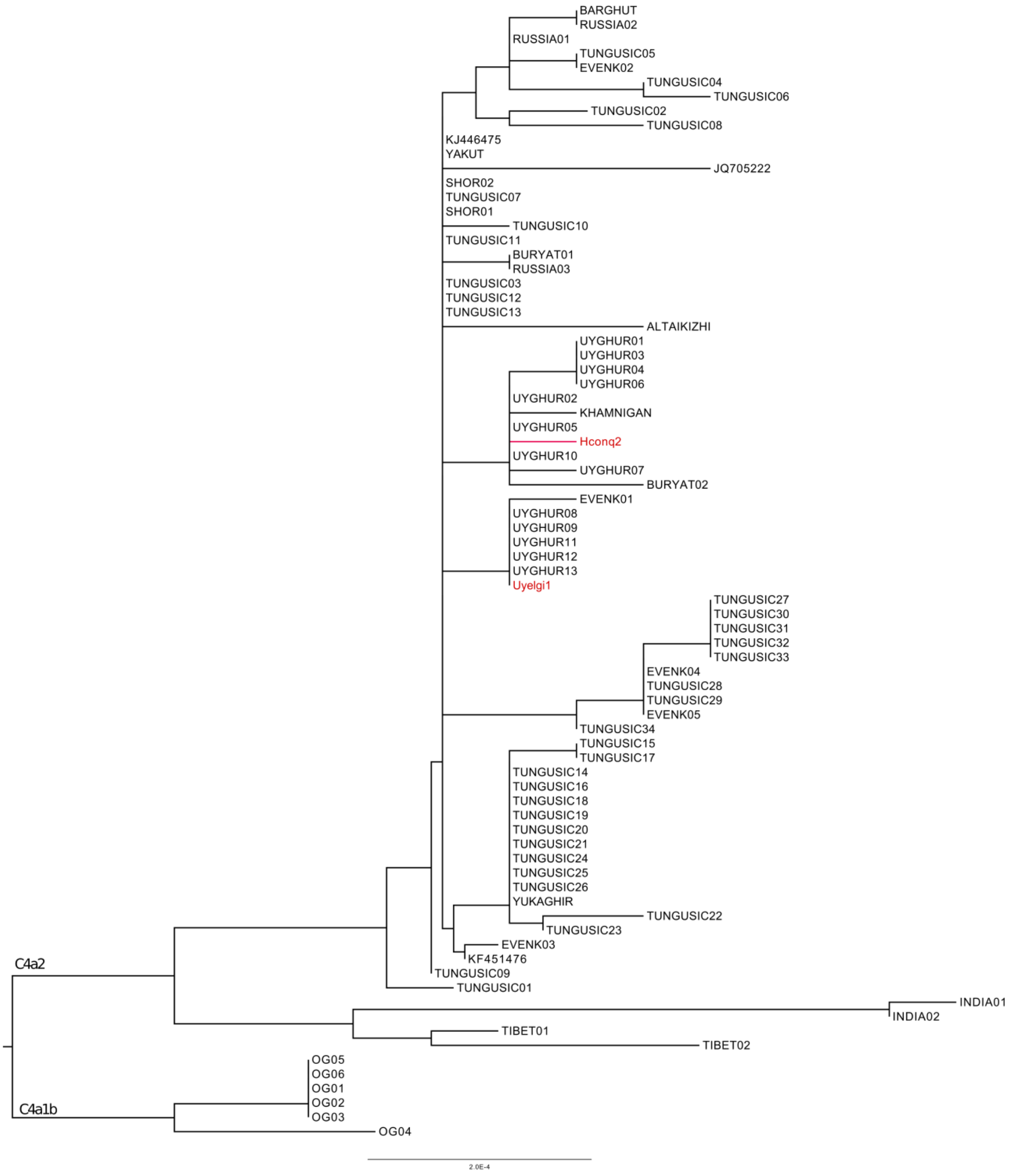

**Fig. S4d Phylogenetic tree of haplogroup C4a2a1.**

The subhaplogroup C4a2a1 can be found in the Kurgan 29 Grave 1, 2 (Uyelgi1) from the newest horizon of the Uyelgi site (10-11$^{th}$ centuries), and in a Hungarian Conqueror grave (Hconq2) from M43-Makó Igási járandó site, (10$^{th}$ century). Although the distribution of the subhaplogroup can be attested to a relatively definable area of Central and Eastern Siberia, the individuals in question are separated by a number of samples from different regions impeding their direct connection (for the abbreviation and further information see Supplementary Table S7).

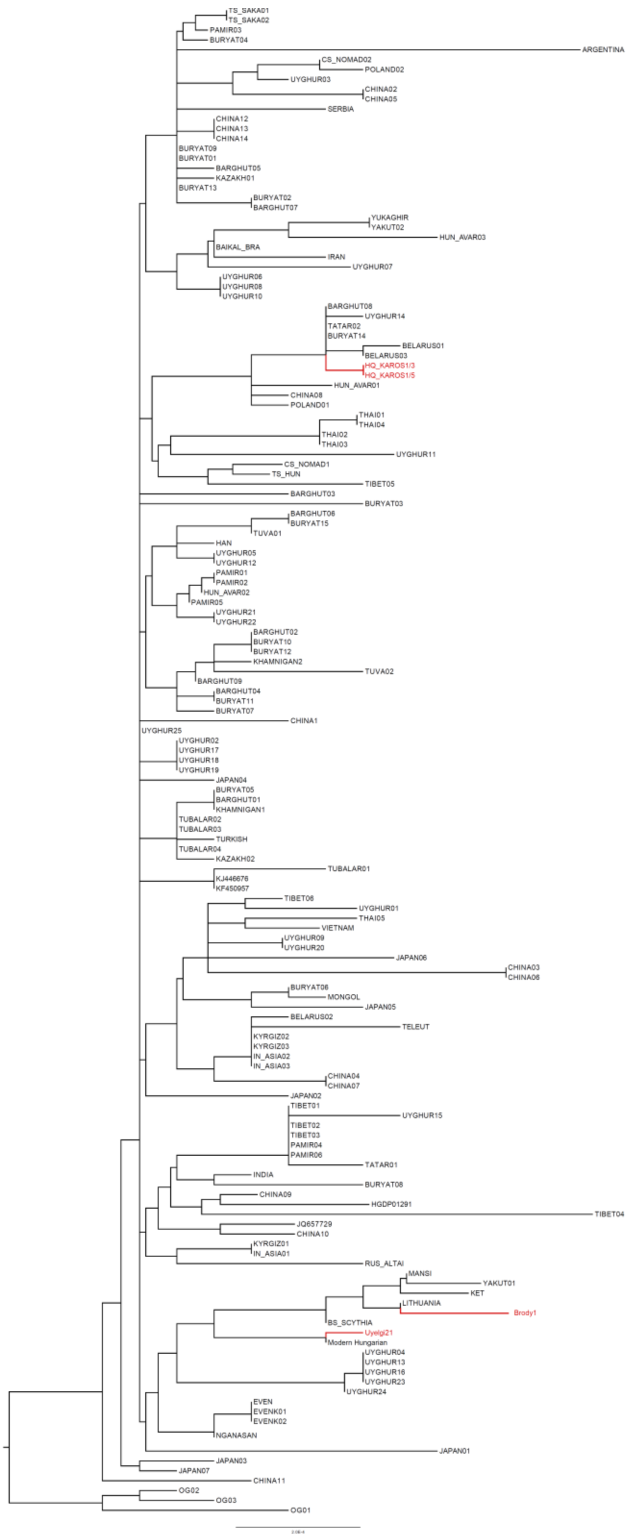

**Fig. S4e Phylogenetic tree of haplogroup D4j.**

The subhaplogroup D4j can be found in Uylegi sample from the oldest horizon, 9th century (Kurgan 11, Grave 5), furthermore D4j2 subhaplogroup in the Brody site, 3rd-4th centuries (Kurgan 8, Grave 2). The investigated sample (Brody1) is located together with individual from Lithuania, furthermore with Mansi, Yakut and Ket individuals on one branch. Interestingly, The Uyelgi21 sample with D4j haplogroup clustered together with a present-day Hungarian individual (for the abbreviation and further information see Supplementary Table S7).

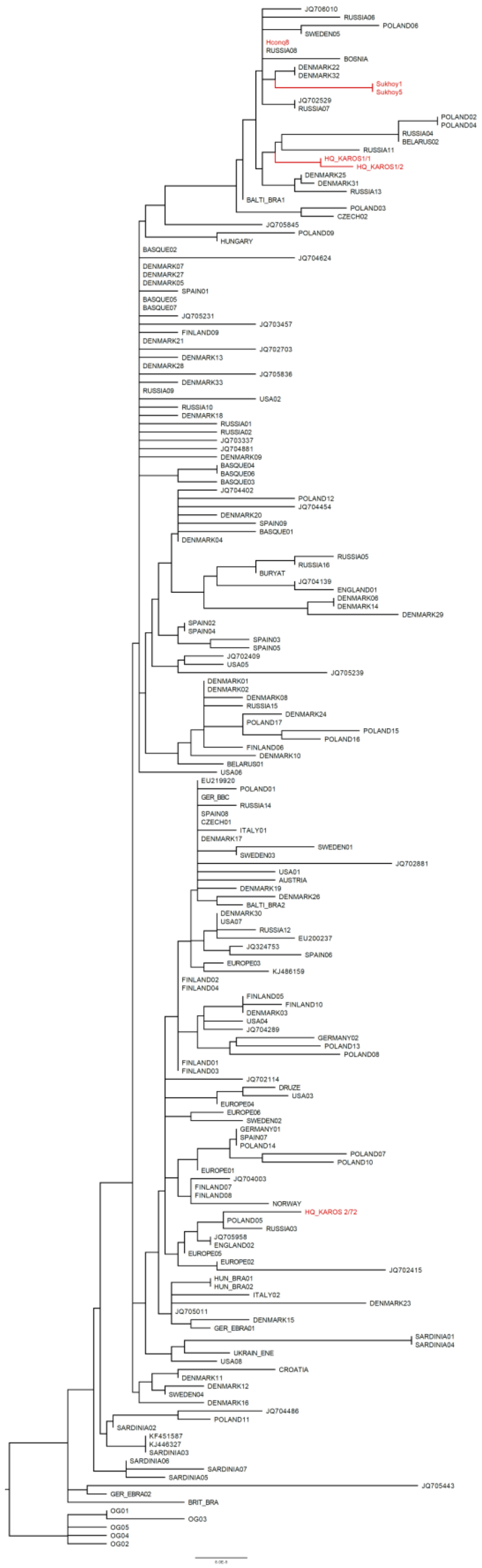

**Fig. S4f Phylogenetic tree of haplogroup H1b2.**

The subhaplogroup H1b2 can be found in two graves from Sukhoy Log site, from the 7–8th centuries (Sukhoy1 and Sukhoy5) and in three Hungarian Conqueror graves from

Nyíregyháza (Hconq8) and Karos sites, 9–10th centuries (Karos1/1 and Karos1/2)[17]. Although the distribution of the subhaplogroup on the main branch containing the mentioned samples can be attested to a relatively definable area of Northern and Eastern Europe, the individuals in question are separated by a number of samples from different regions impeding their direct connection, except the identical (Sukhoy1 and Sukhoy5) and similar (Karos1/1 and Karos1/2) pairs which may represent close maternal kinship within the Sukhoy Log and Karos sites (for the abbreviation and further information see Supplementary Table S7).

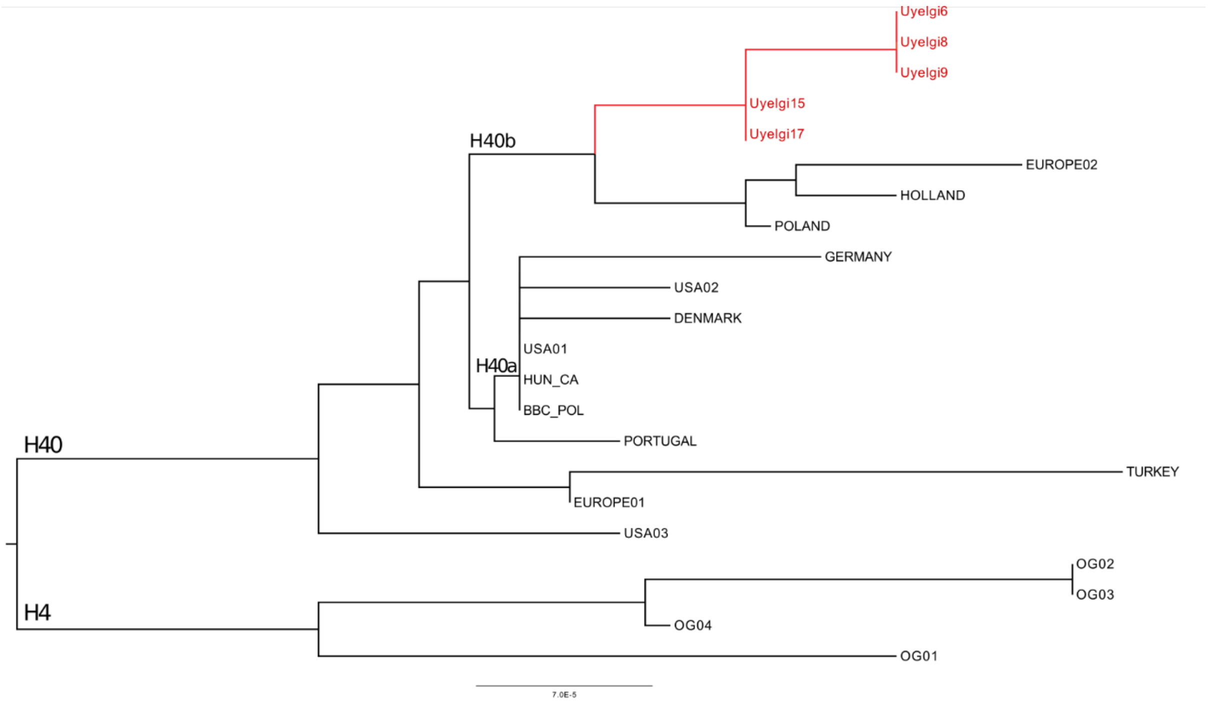

**Fig. S4g Phylogenetic tree of haplogroup H40b.**

The subhaplogroup H40b can be found in five individuals from two horizons of Uyelgi site: Grave 5 and Grave 7 of Kurgan 9 (Uyelgi15 and Uyelgi17) from the middle horizon (910th centuries) and in Kurgan 28 Graves 5, 6 (Uyelgi6 and Uyelgi9) and Kurgan 29 Grave 1 (Uyelgi8) from the newest horizon (10-11th centuries). The middle horizon has two identical mitochondrial lineages, which are genetically ancestral to the three identical mitogenomes of the newest horizon, connecting the two horizons together (for the abbreviations and further information see Supplementary Table S7).

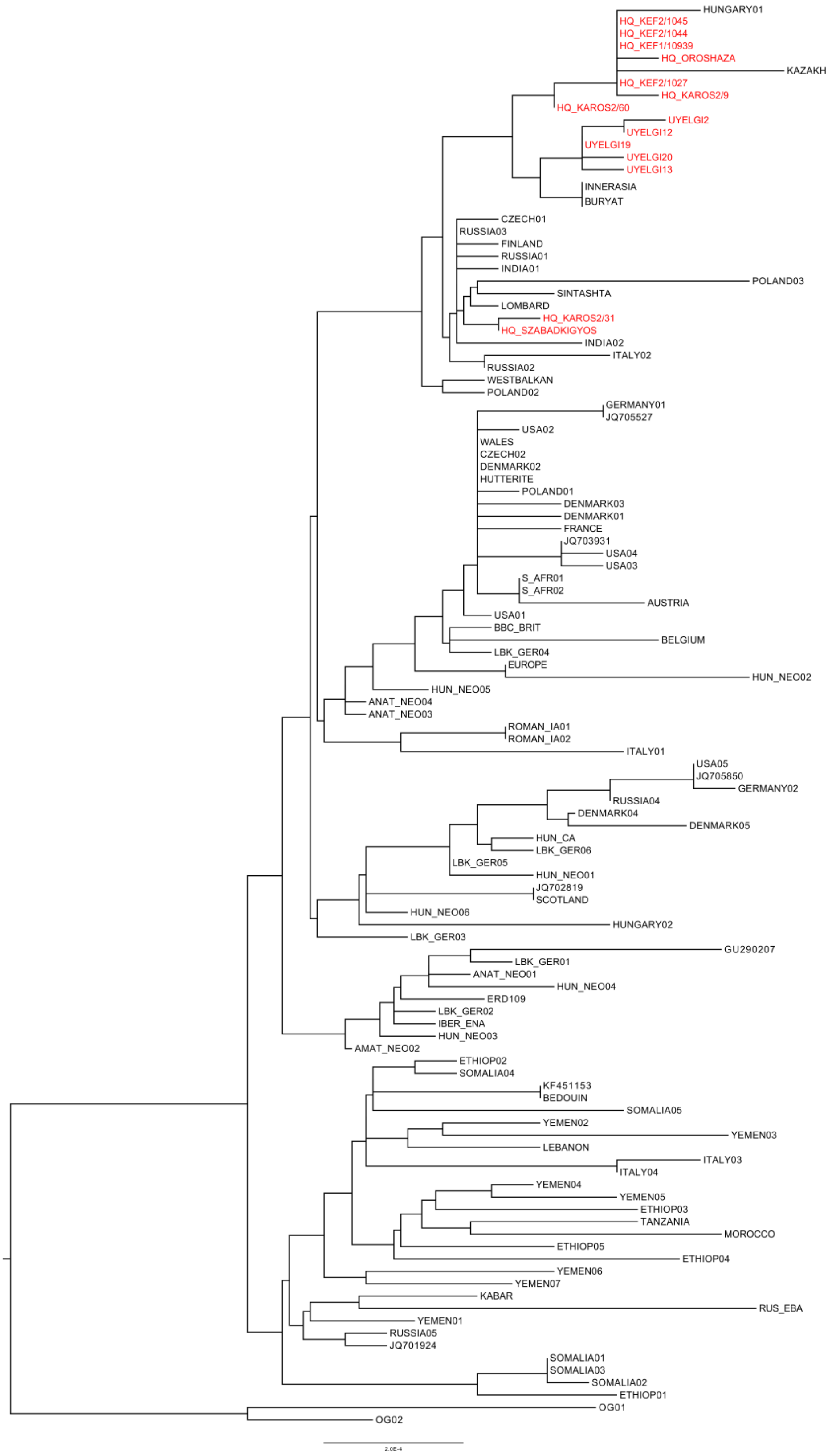

**Fig. S4h Phylogenetic tree of haplogroup N1a1a1a(1a).**

The subhaplogroup N1a1a1a1a was detected in five individuals assigned to two horizons of Uyelgi site: Graves 12 and 16 of Kurgan 32 (Uyelgi19 and Uyelgi20) from the oldest (9$^{th}$ century) horizon, and Grave 3 of Kurgan 36 (Uyelgi13), Graves 5 and 6 of Kurgan 28

(Uyelgi2 and Uyelgi12) from the newest horizon (10–11th centuries), furthermore, in nine Hungarian Conquerors from various cemeteries in 9–10th centuries Carpathian Basin (Karos site: Karos2/31; Karos2/60; Karos2/9; Kenézlő-Fazekaszug site: KeF2/1044; KeF2/1045; KeF1/10939; KeF2/1027; Szabadkígyós-Pálliget; Orosháza-Görbicstanya) and in one modern Hungarian. Two Hungarian Conqueror maternal lineages (Karos2/31 and Szabadkígyós-Pálliget) cannot be attested to special geographic origin, since their positions are too basal, and they have no proximate relatives, which impede further analysis in this level. The Hungarian Conqueror individuals: KeF2/1045, KeF2/1044, Kef1/10939, Orosháza, Kef2/1027, Karos2/9 and Karos2/60 (along with the modern Hungarian) form another relatively compact branch, are interposed with a Kazakh sample. The Uyelgi branch is very compact, clearly connects the oldest and newest horizons together, and clusters with Hungarian Conquerors in one main branch but samples from the two regions are separated in sub branches (for the abbreviations and further information see Supplementary Table S7).

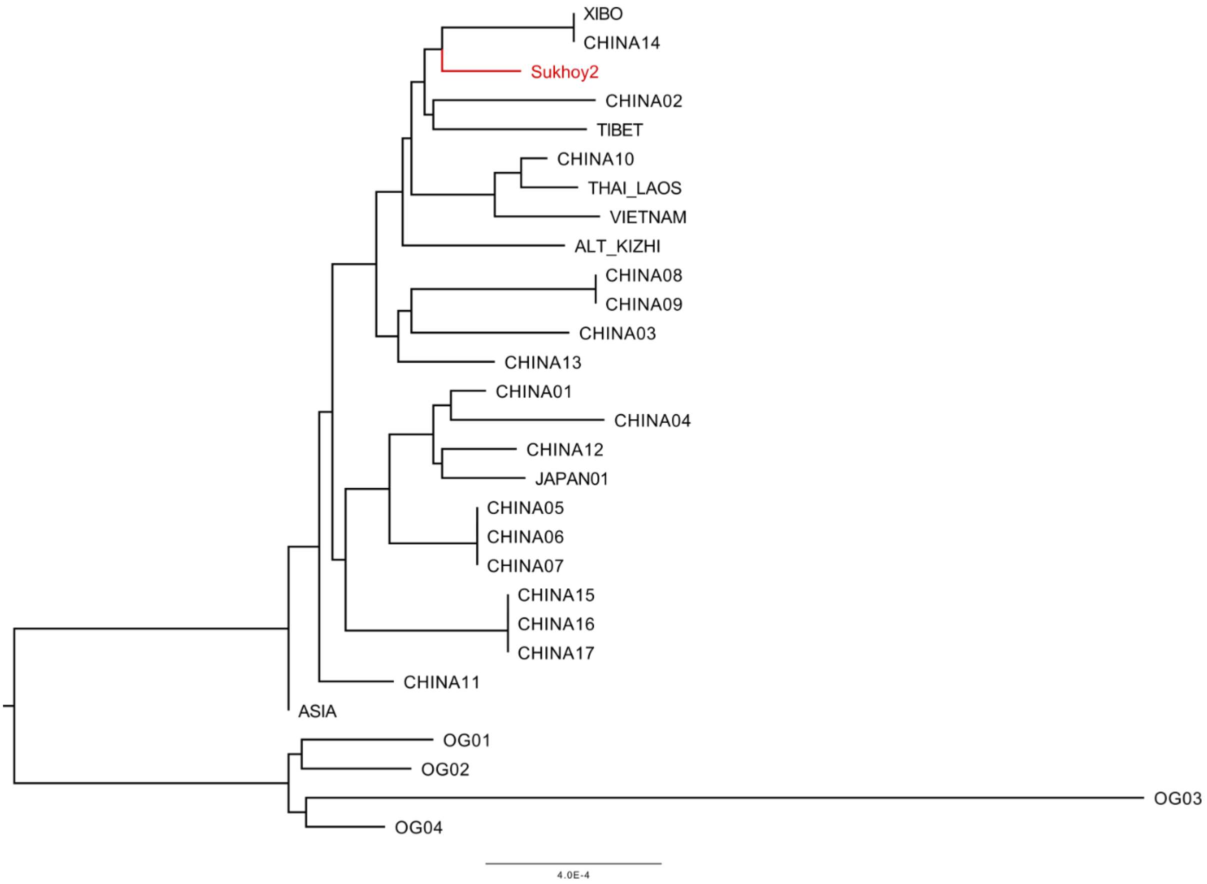

**Fig. S4i Phylogenetic tree of haplogroup R11b1b.**

The subhaplogroup R11b1 can be found in individual Sukhoy2 from 7–8th centuries Sukhoy Log site. The overall structure of the phylogenetic tree suggests a Central-Eastern Asian origin and distribution of this lineage; however, the deep divergences are preventing further analyses. Interestingly, the Sukhoy2 sample clusters together with East Asian Xibo and Chinese mtDNA lineages (for the abbreviations and further information see Supplementary Table S7).

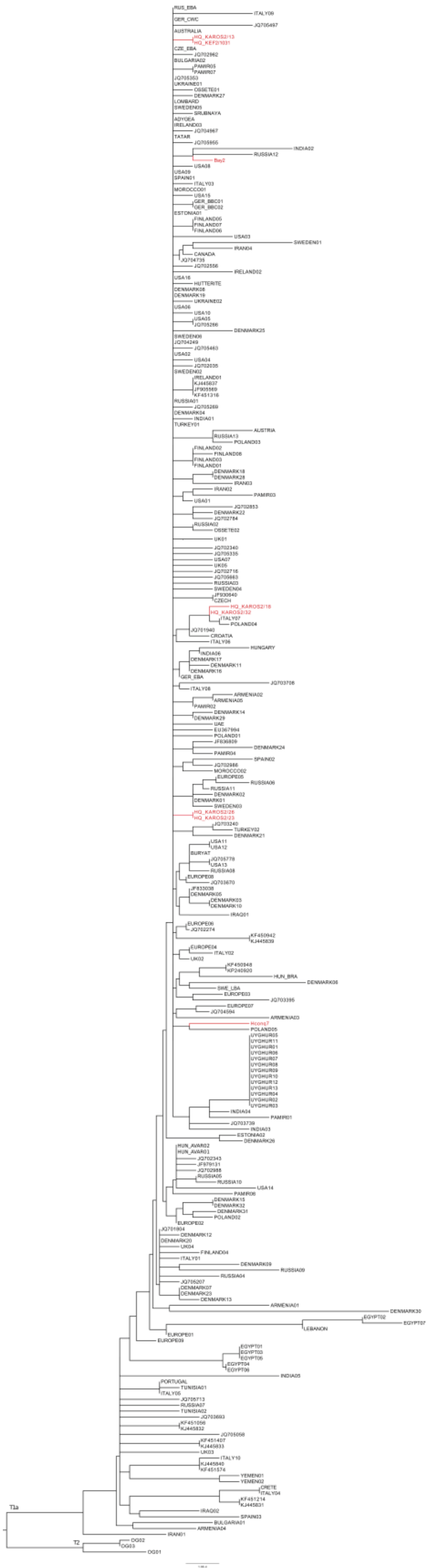

**Fig. S4j Phylogenetic tree of haplogroup T1a1.**

The subhaplogroup T1a1 can be found in the 10th century Bayanovo cemetery (Bay2). This haplogroup are widespread with many identical haplotypes and without any stronger connection to geographical locations (for the abbreviation and further information see Supplementary Table S7).

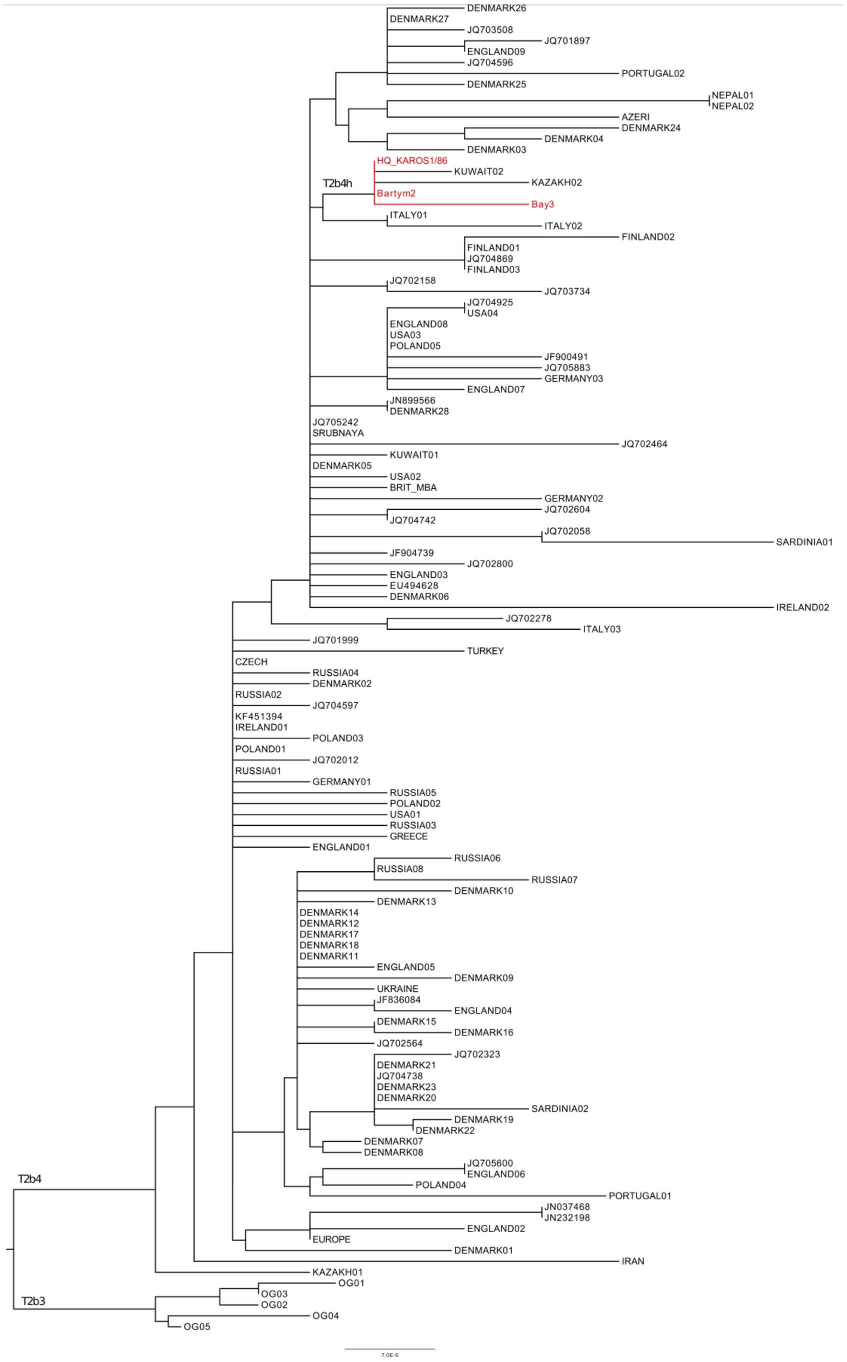

**Fig. S4k Phylogenetic tree of haplogroup T2b4h.**

The subhaplogroup T2b4h can be found in a cemeteries Bartym (Bartym2, 5-6$^{th}$ centuries) and Bayanovo (Bay3, 10$^{th}$ century) which are located on one branch together with a Hungarian Conqueror individual from Karos site (Karos1/86, 9–10$^{th}$ centuries). The phylogenetic proximity of the samples is apparent, the Karos and the Bartym individuals have even identical mtDNA sequences, and their lineage is maternally ancestral to the Bayanovo samples mtDNA and to a Kazakh and Kuwait samples (for the abbreviations and further information see Supplementary Table S7).

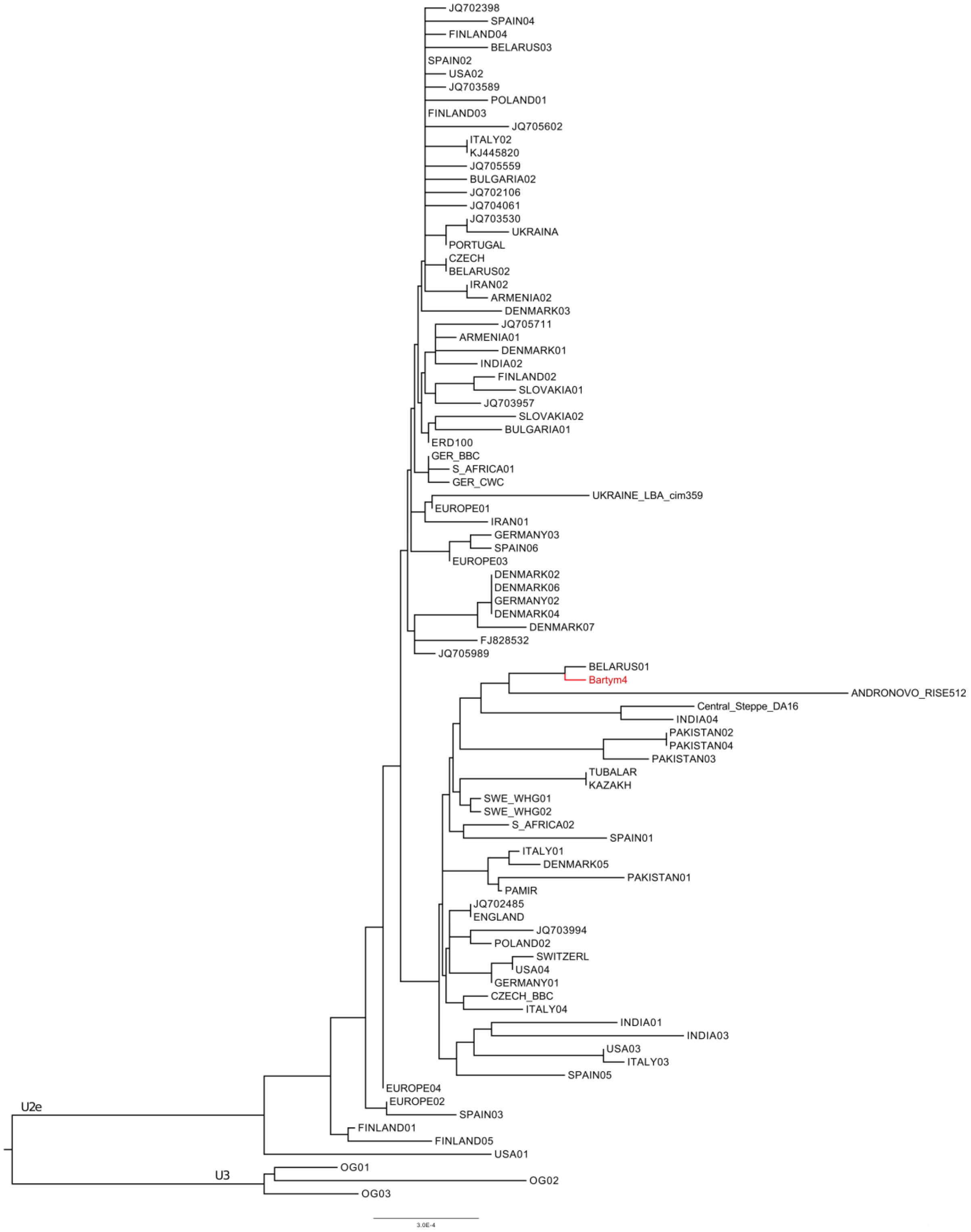

**Fig. S4l Phylogenetic tree of haplogroup U2e1.**

The subhaplogroup U2e1 can be found in a Bartym cemetery (Bartym4, 5-6th centuries). The Bartym individual's position on the tree highly supports local characteristics for his mtDNA (for the abbreviations and further information see Supplementary Table S7).

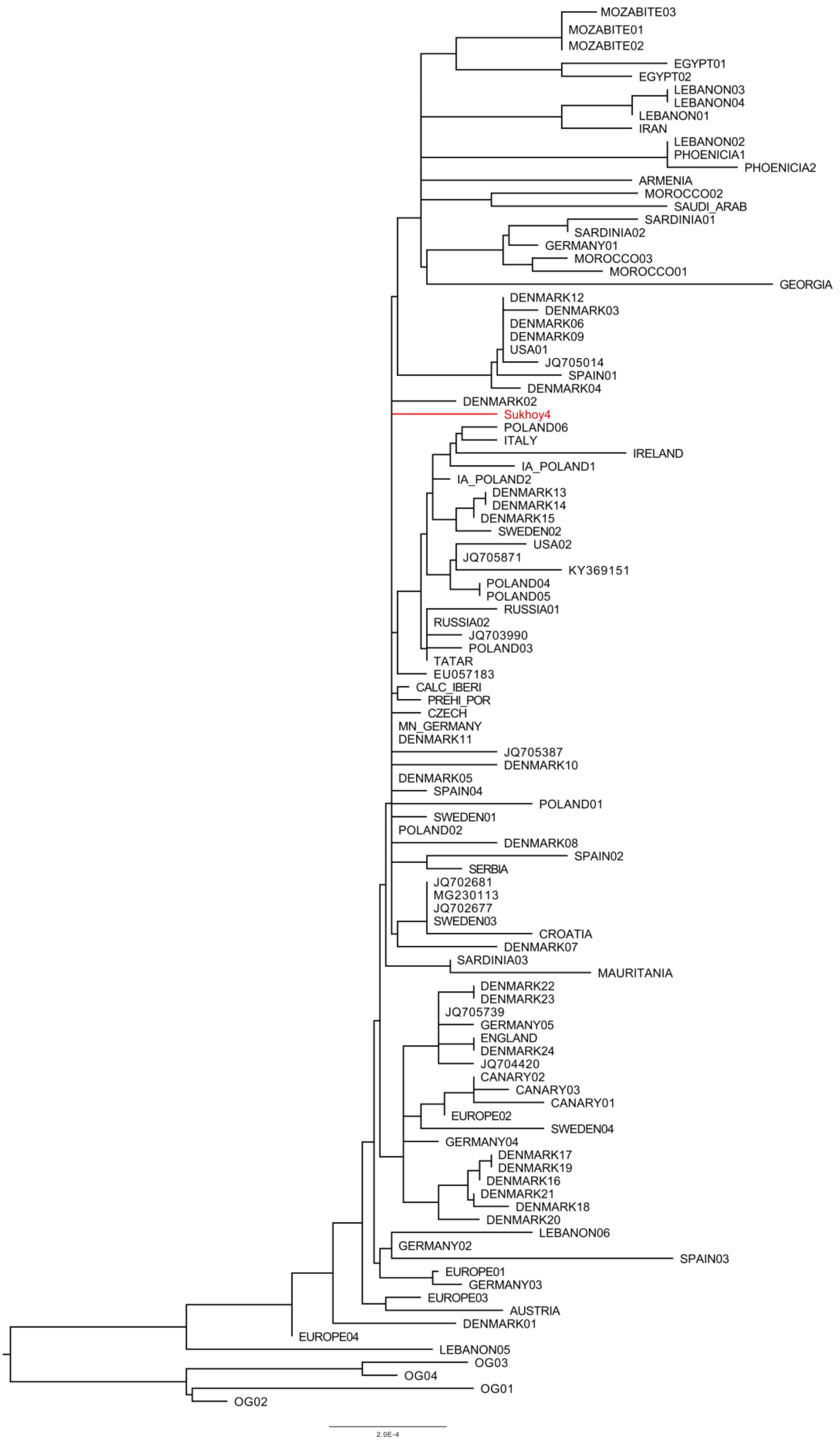

**Fig. S4m Phylogenetic tree of haplogroup U3a1.**

The subhaplogroup U3a1 can be found by grave from Sukhoy Log site, 7-8th centuries (Sukhoy4). The surroundings of the sample in the phylogenetic tree points to a rather North European characteristic, which might indicate either the local origin (as the result of locality due to a slight eastward allocation of the lineage), or some direct Northern European descent (for the abbreviations and further information see Supplementary Table S7).

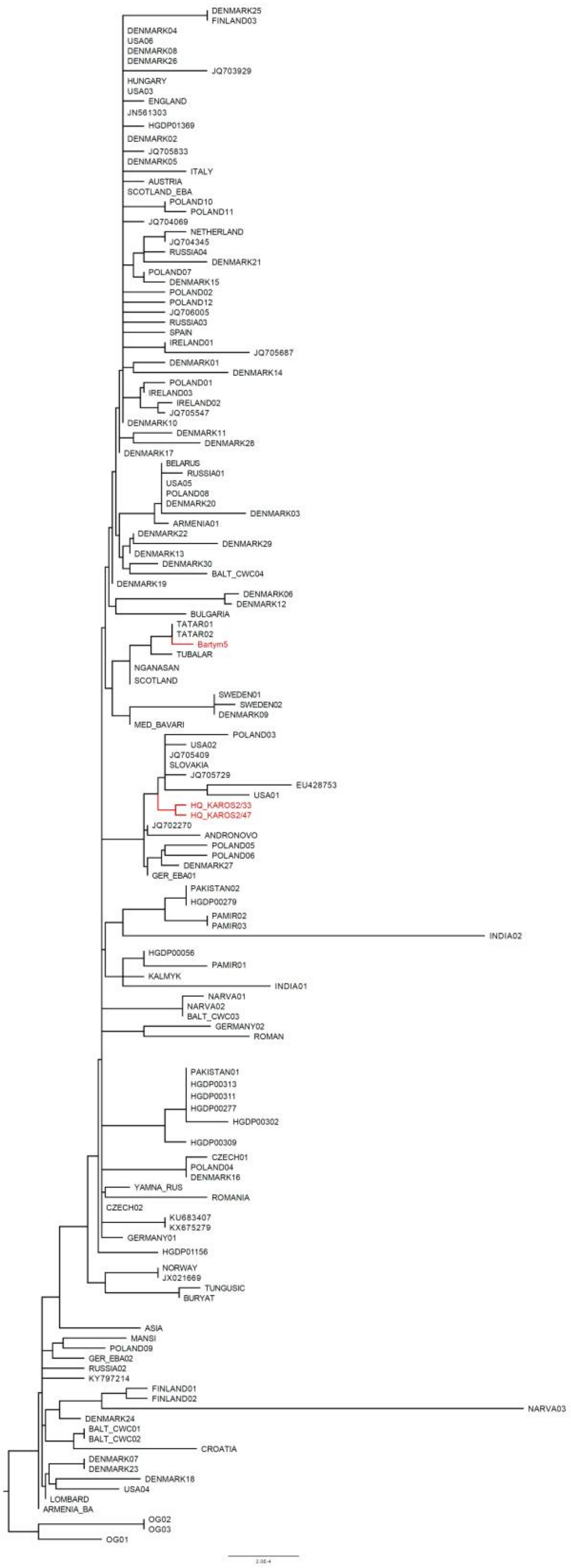

**Fig. S4n Phylogenetic tree of haplogroup U4a1d.**

The subhaplogroup U4a1d can be found in grave from Bartym site, 5–6$^{th}$ century (Bartym5). The origin of the subhaplogroup can be attested to Europe with the appearance of some sporadic Asian lineages. The Bartym5 individual's proximity to Siberian and Pontic steppe samples presumes the origin of the lineage from this region (for the abbreviations and further information see Supplementary Table S7).

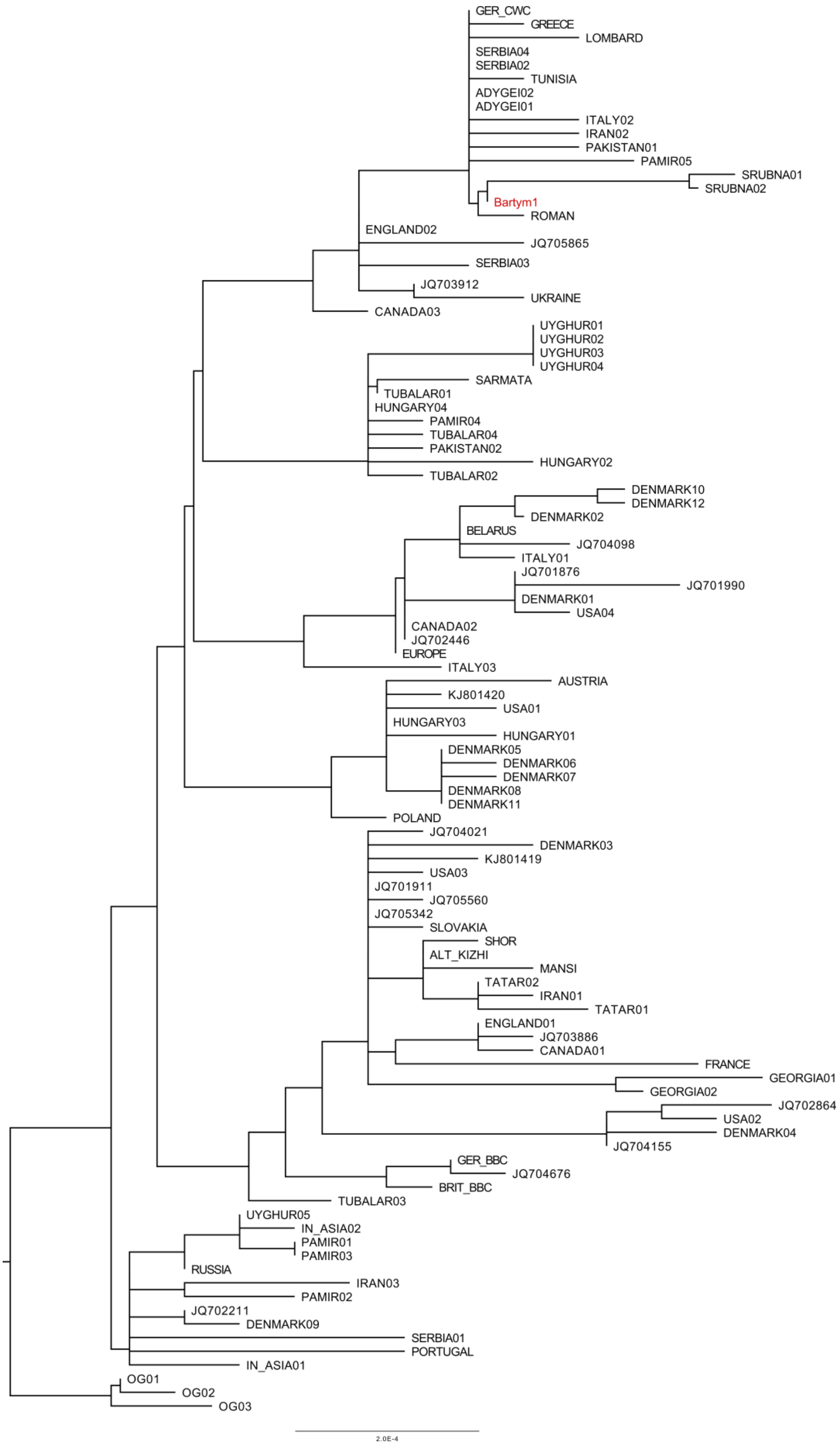

**Fig. S4o Phylogenetic tree of haplogroup U4b1a1a1.**

The subhaplogroup U4b1a1a1 can be found in grave from Bartym site, 5–6$^{th}$ centuries (Bartym1). The Bartym1 sample is located together with two individuals related to Bronze Age Srubnaya culture from Republic of Bashkorstan, Near Usmanovo village, Kazburun[18], where the lineage of Bartym1 clustered with two Srubnaya culture related individuals mtDNA (for the abbreviation and further information see Supplementary Table S7).

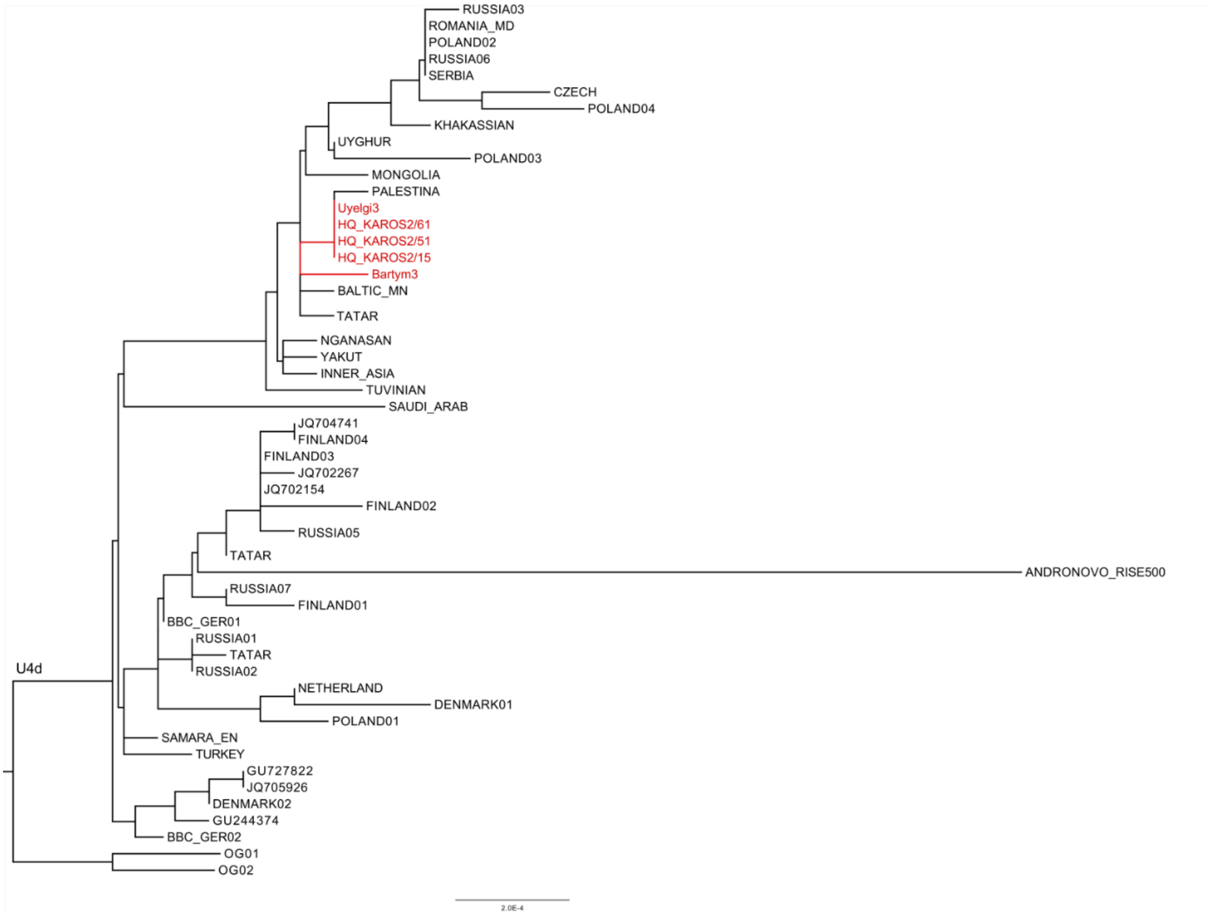

**Fig. S4p Phylogenetic tree of haplogroup U4d2.**

The subhaplogroup U4d2 can be found in cemeteries Uyelgi (Uyelgi3, 10-11$^{th}$ centuries) and Bartym (Bartym3, 5-6$^{th}$ centuries). The mtDNA of Uyelgi sample is identical to the mitogenomes of three Hungarian Conqueror individuals from Karos cemetery (9-10$^{th}$ centuries), which represents close or even direct maternal relationship between these individuals. The individual from Bartym located on the same main branch only may show relatedness to the geographic area of Ural region, but no further statements can be made (for the abbreviations and further information see Supplementary Table S7).

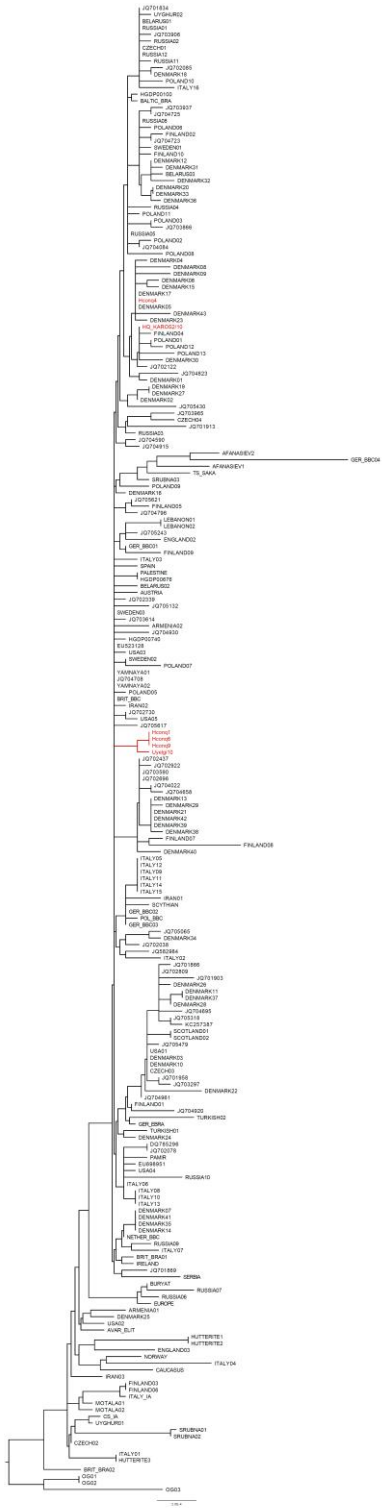

**Fig. S4q Phylogenetic tree of haplogroup U5a1a1.**

The subhaplogroup U5a1a1 was detected in the middle horizon of Uyelgi site, 9–10th centuries (Uyelgi10) and in five Hungarian Conqueror samples Hconq1 and Hconq6 both from Balatonújlak-Erdődűlő, Hconq4 from Harta-Freifelt, Hconq9 from Makó-Igási járandó and Karos2/10). The Uyelgi10 and three Hungarian Conqueror samples (Hconq1, Hconq6 and Hconq9) group together, assuming close relationship between them. The remaining two Hungarian Conquerors (Hconq4 and Karos2/10) does not show any direct connection to the others, but their origins can be more confidently attested to Northern Europe (for the abbreviations and further information see Supplementary Table S7).

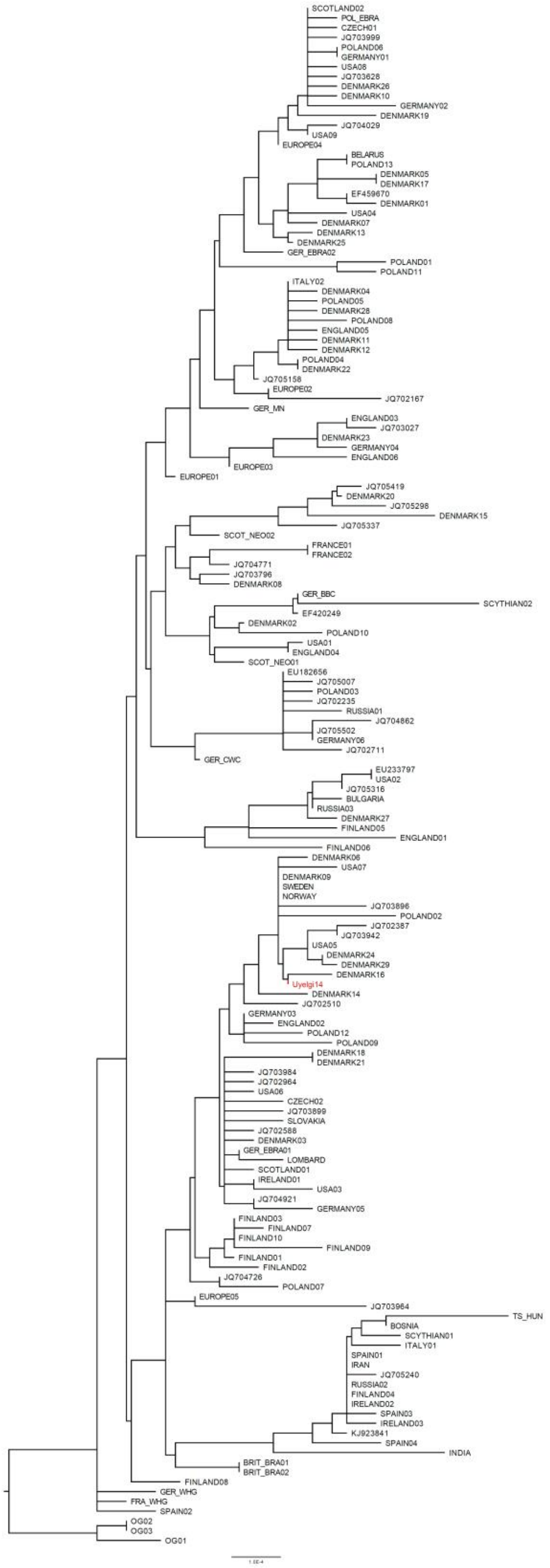

**Fig. S4r Phylogenetic tree of haplogroup U5b2a1a1.**

The subhaplogroup U5b2a1a1 can be found in grave from the newest horizon of Uyelgi site, 10-11th centuries (Uyelgi14). The distribution of the subhaplogroup can be attested confidently to Central and Northern Europe, where the Uyelgi14 individual is the only Eastern representative of this lineage (for the abbreviations and further information see Supplementary Table S7).

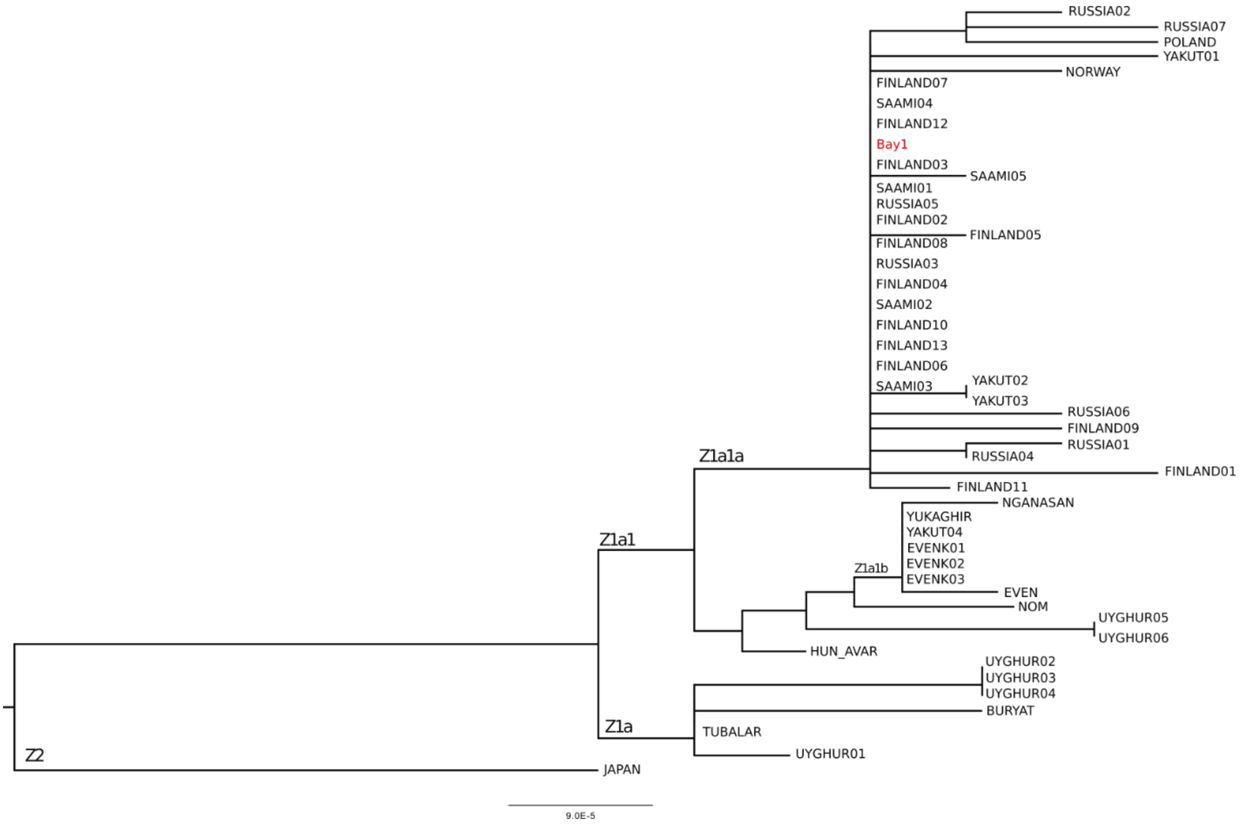

**Fig. S4s Phylogenetic tree of haplogroup Z1a(1a).**

The subhaplogroup Z1a1a can be found in Bayanovo cemetery (Bay1, 10$^{th}$ century). The mitochondrial lineage of Bayanovo individual is identical to many modern mostly Scandinavian individuals´ mitogenomes and seems to be a possibly genetic ancestor of these Western (Scandinavian) lineages (for the abbreviations and further information see Supplementary Table S7). The subhaplogroup Z1a1a appeared most recently (1.6-0.8 kya) presumably by Kets in Volga Ural region and/or by Finn-Saami populations[19].

#### 5. Supplementary figures and descriptions of mitochondrial population genetic and Y-chromosomal network results

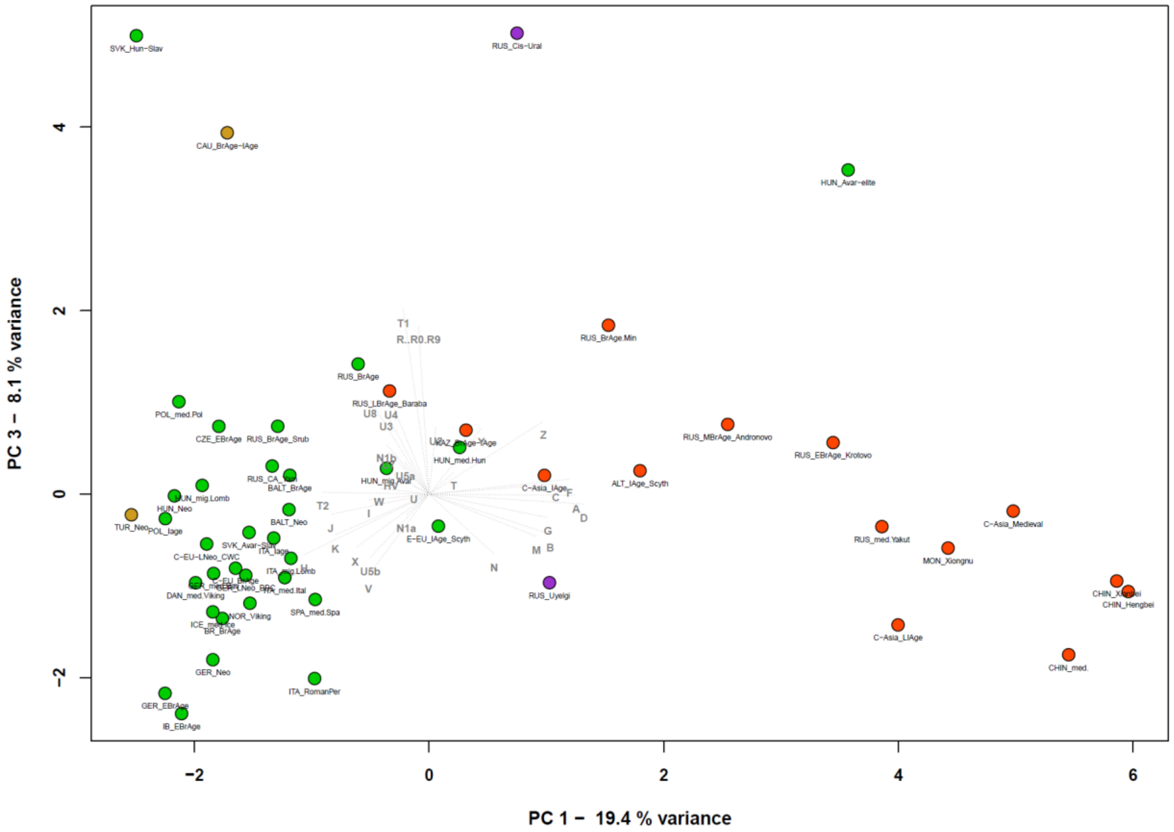

**Fig. S5 PCA plot with 50 ancient populations, representing first-third principal components.**

PCA plot revealed predominant differences between Asian and European populations. Even though the Asian populations were relatively far from each other, they firmly spanned on the right side of plot along the PC1 and PC3. Investigated population from Uyelgi (RUS_Uyelgi) clustered together with the Iron age population from Central Asian Steppe (C-Asia_IAge), the Scythians from Altai region (ALT_IAge_Scyth), East European Iron Age Scythians (E-EU_IAge_Scyth ) and Medieval population of Hungarian conquerors (HUN_med.HUN). Despite of the separation of populations from Cis-Ural region (RUS_Cis-Ural) along the PC3, the Caucasian Bronze Age and Iron Age populations (CAU_BrAge-IAge) as well as Hungarian-Slavic population from Slovakia (SVK_Hun-Slav) and the Avar-elite population from Hungary (HUN_Avar-elite) are also detached together with this group. The far-off position of Cis-Ural population along PC3 is inflicted by presence of both haplogroups R and T1, which haplogroups have the highest influence on this component (for the frequencies of haplogroups and the reference see Supplementary Table S4).

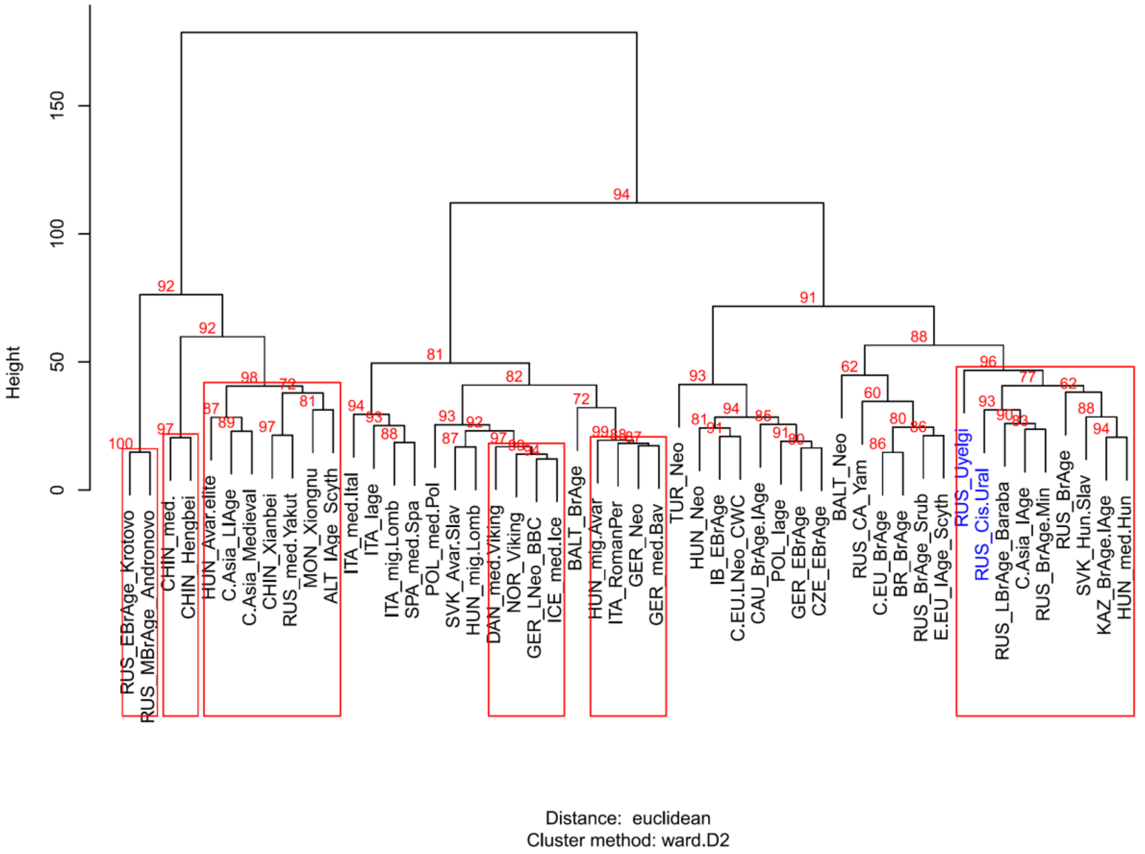

**Fig. S6 Ward type clustering of the 50 ancient populations.**

The segregation and clustering of European and Asian ancient populations are displayed on the Ward type clustering tree. The investigated population from Uyelgi (RUS_Uyelgi) and Cis-Ural RUS_Cis-Ural) is situated on the mixed European–Asian branches of Ward-tree, together with Hungarian conquerors (HUN_med.Hun) in the same main branch. The Cis-Ural (RUS_Cis-Ural) clustered furthermore in a sub-branch with Late Bronze Age Siberian Baraba population (RUS_LBrAge_Baraba), Central Asian population from Iron age (C.Asia_IAge) and Bronze Age population from Minusinsk Depression (RUS_BrAge.Min) (for the frequencies of haplogroups and the reference see Supplementary Table S3).

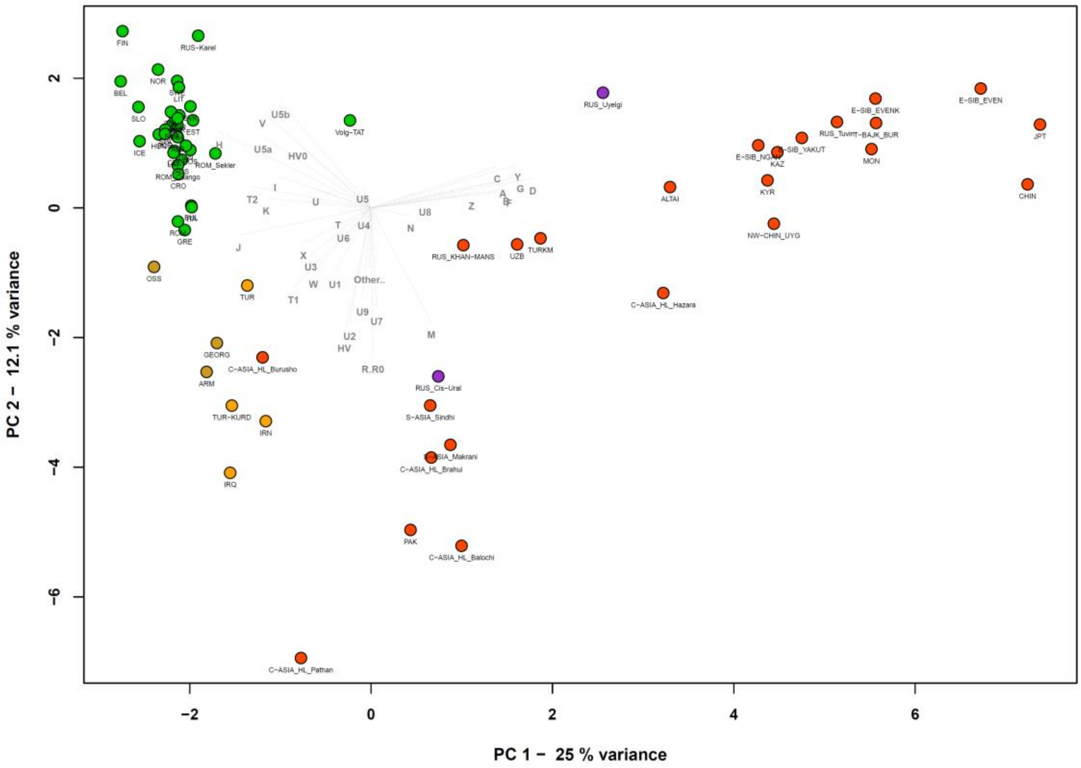

a)

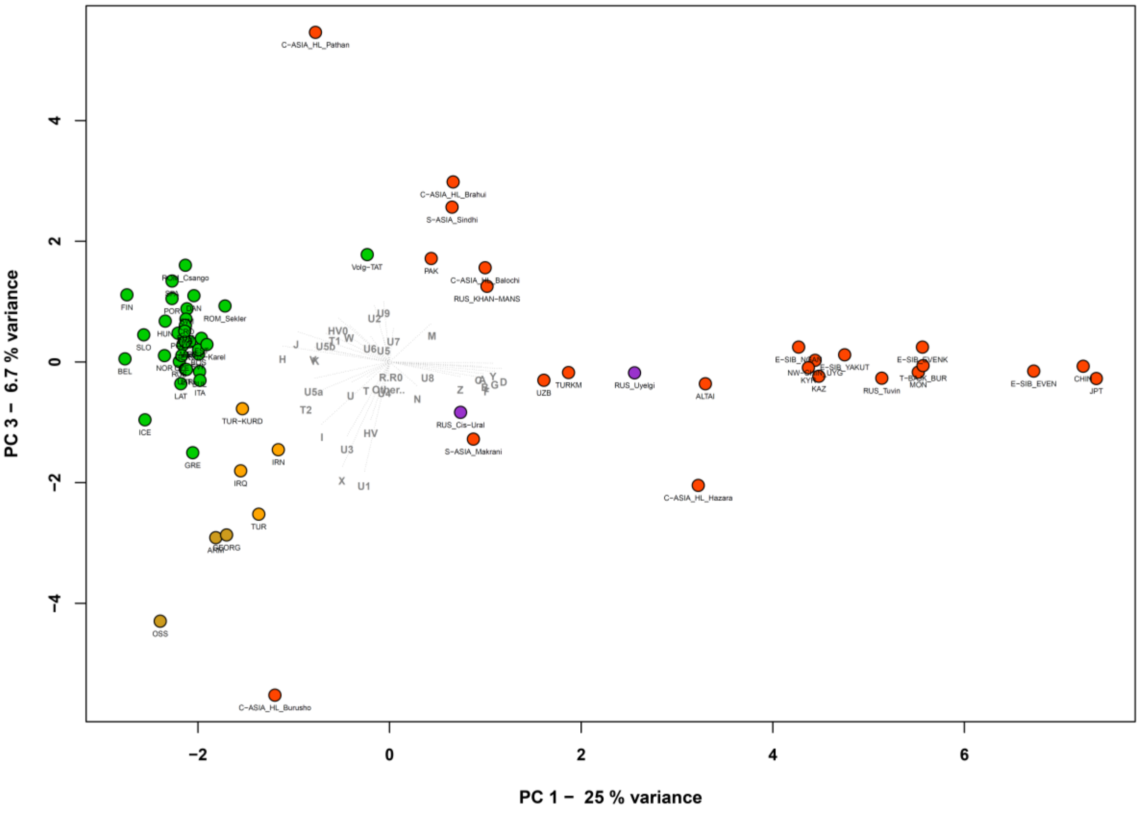

b)

**Fig. S7 PCA plots with 64 modern and two investigated ancient populations from Ural region, representing first two principal components (a) and first-third principal components (b); (43.8 % of the total variance is shown).**

The PCA plots show the clear separation of Asian and European populations, moreover the populations form Near East and Central-Asian ethnic groups segregated along the PC2. The Uyelgi population (RUS_Uyelgi) is standing along the PC1 and PC2 between East- and Central-Asian populations, and along PC3 locates relatively near to populations from Turkmenistan (TURKM), Uzbekistan (UZB) and Altai region (ALTAI), and to Central Asian ethnic group Hazara (C-ASIA_HL_Hazara); whereas the Cis-Ural (RUS_Cis-Ural) is near to ethnic groups from Central Asia along all three components (for the frequencies of haplogroups and the reference see Supplementary Table S4).

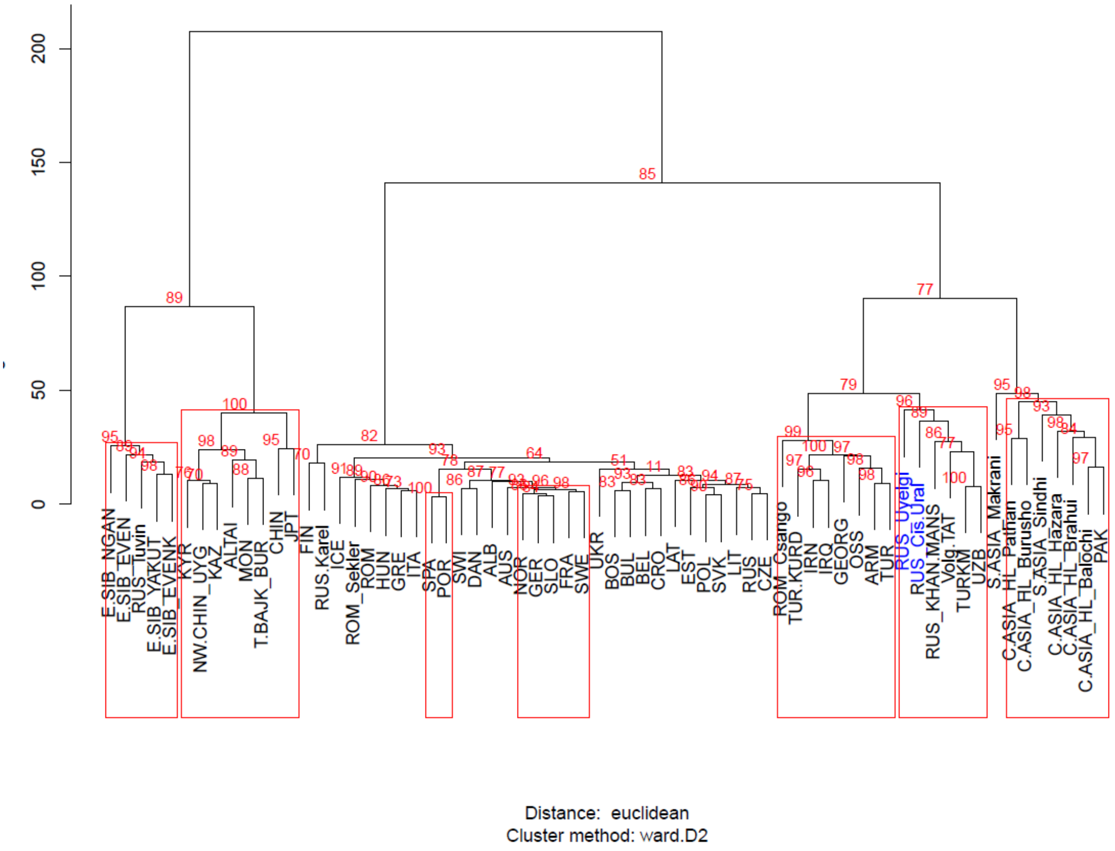

**Fig. S8 Ward type clustering of 64 modern and two investigated ancient populations from Ural region.**

The ward clustering tree presents separation of East- and Central-Asian, European, Caucasian – Near Eastern and Central-South Asian populations in a three major branches. The two medieval populations Uyelgi (RUS_Uyelgi) and Cis-Ural (RUS_Cis-Ural) are situated on the Central-South Asian and Near Eastern branch, in the same sub-branch together with Central-Asian and Finno-Ugric populations (Khanty and Mansi, RUS_KHAN.MANS) ang Volga Tatars from Russia (for the frequencies of haplogroups and the reference see Supplementary Table S4).

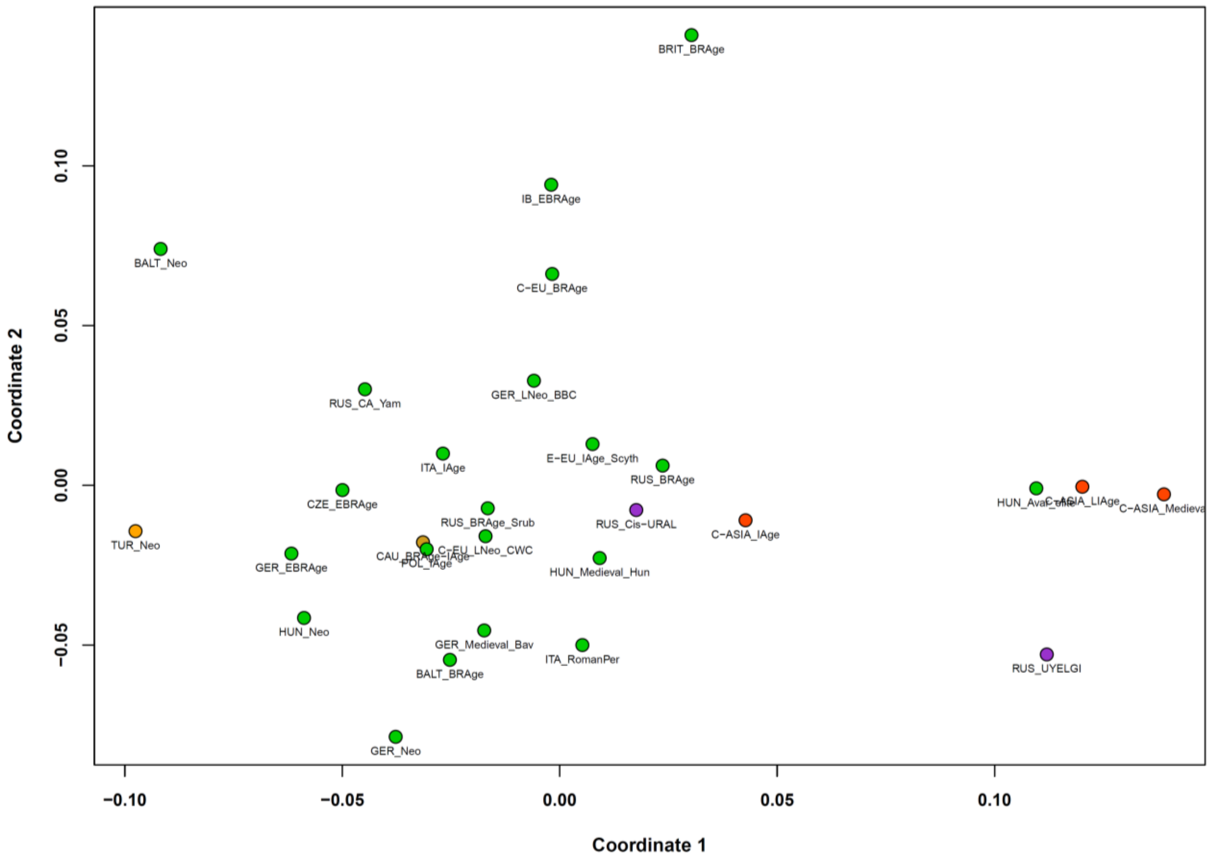

Coordinate 2
Coordinate 1
0.10
0.05
0.00
−0.05
−0.10
−0.05
0.00
0.05
0.10
BRIT_BRAge
IB_EBRAge
BALT_Neo
C−EU_BRAge
RUS_CA_Yam
GER_LNeo_BBC
ITA_IAge
E−EU_IAge_Scyth
RUS_BRAge
CZE_EBRAge
RUS_BRAge_Srub
RUS_Cis−URAL
C−ASIA_IAge
C−ASIA_LIAge
TUR_Neo
GER_EBRAge
C−EU_LNeo_CWC
POL_IAge
HUN_Medieval_Hun
HUN_Neo
GER_Medieval_Bav
ITA_RomanPer
BALT_BRAge
RUS_UYELGI
GER_Neo

**a)**

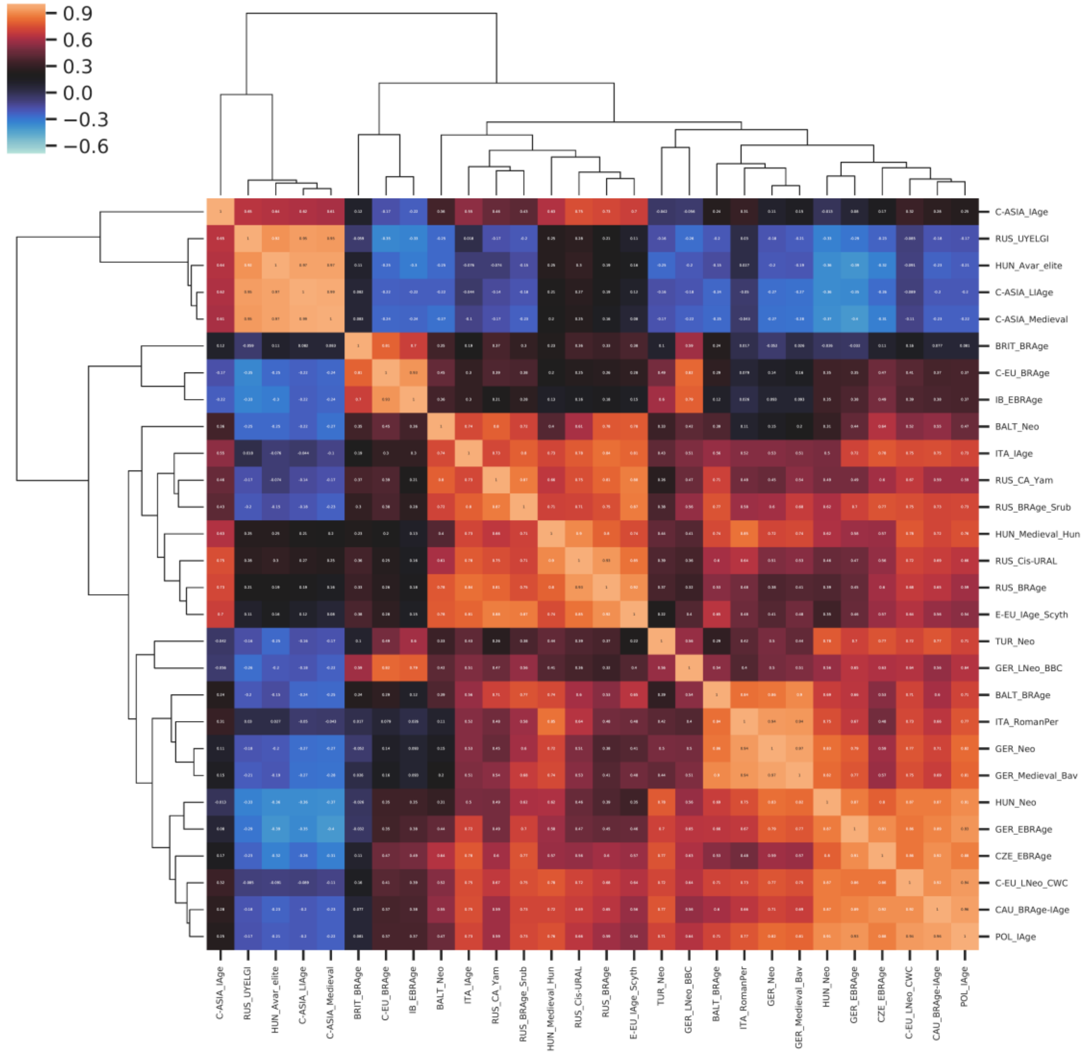

**b)**

**Fig. S9 MDS plot of 28 ancient populations (stress value = 0.1638) (a) and the Spearman´s correlation heatmap of the $F_{ST}$ values (b)**

a) The Cis-Ural (RUS_Cis-Ural) population showed affinities to the populations of medieval Hungarian Conquerors (HUN_Medieval_Hun), Russian Bronze Age (RUS_BRAge), Scythians from Europe (E-EU_IAge_Scyth) and Iron Age populations from Central Asian Steppe (C-ASIA_IAge) along coordinates 1 and 2 and is situated between European and Asian populations, which reflects on the $F_{ST}$ values. The Uyelgi is standing relatively far from any ancient populations, which is might be caused by significant genetic distances from all ancient populations (for the $F_{ST}$ values, p values, linearized Slatkin values and references see Supplementary Table S5).

b) Based on Spearman´s correlation heatmap the Uyelgi (RUS_Uyelgi) clustered together with Central Asian populations of Iron Age (C-ASIA_IAge), Late Iron Age (C-ASIA_LIAge) and medieval period (C-ASIA_Medieval), as well as with Avar Elite from Hungary, while the Cis-Ural (RUS_Cis-Ural) clustered with Hungarian Conquerors (HUN_Medieval_Hun), Russian Bronze Age population (RUS_BRAge) and Scythians from Europe (E-EU_IAge_Scyth).

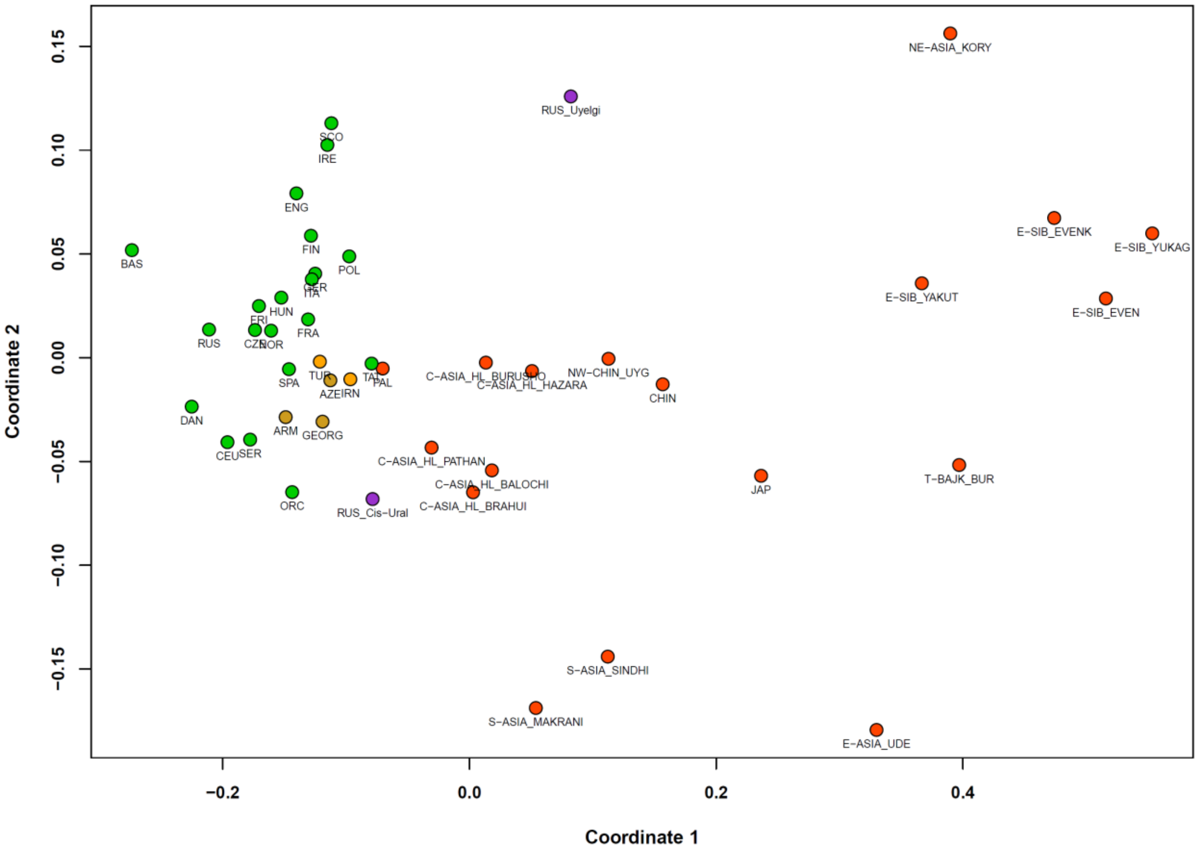

**Fig. S10 MDS plot of 43 modern populations and two investigated ancient populations from Ural region (stress value = 0.06749).**

The MDS plot shows differentiation of European and Asian populations, where Caucasian and Near-Eastern populations are between Europeans and Asians but closer to European populations, and reflects on the $F_{ST}$ values, where investigated population of Cis-Ural (RUS_Cis-Ural) is closer to the populations of Central Asian Highland (Brahui, Balochi and Pathan) (C-ASIA_HL_BRAHUI; C-ASIA_HL_BALOCHI; C-ASIA_HL_PATHAN) and the European and Caucasian populations (e.g. ORC, CEU, SER, ARM, GEORG), while Uyelgi (RUS_Uyelgi) is between European and Asian populations along coordinate 1, but it diverges along coordinate 2 together with Northeast Asian Koryak population (NE-ASIA_KORY) (for the $F_{ST}$ values, p values, linearized Slatkin values and references see Supplementary Table S6).

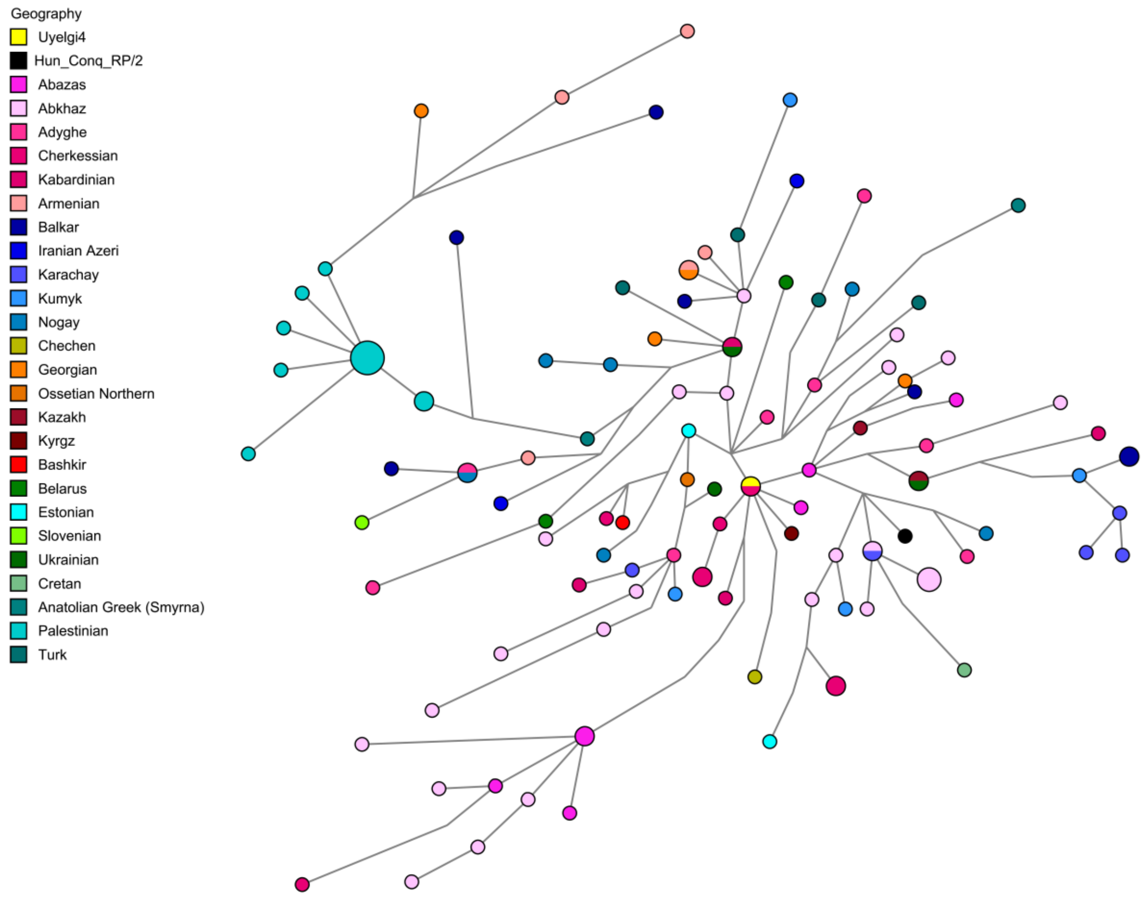

**Fig. S11 Median Joining Network analysis of G2a2-U1 Y-haplogroup based on 14 STRs.** The Uyelgi4 sample (marked with yellow) has identical all investigated 14 STR loci with a Cherkessian individual from Caucasus region. One-step-neighbours to Uyelgi4 are further Cherkessian and two Abazas individuals from Caucasus region and one Kyrgyz individual from Central Asia. The Hungarian Conqueror sample from Rétközberencs-Paromdomb (RP/2) surrounded with Caucasian samples is three-steps away from the Uyelgi4 sample (for the population information, STR data and references see Supplementary Table S9).

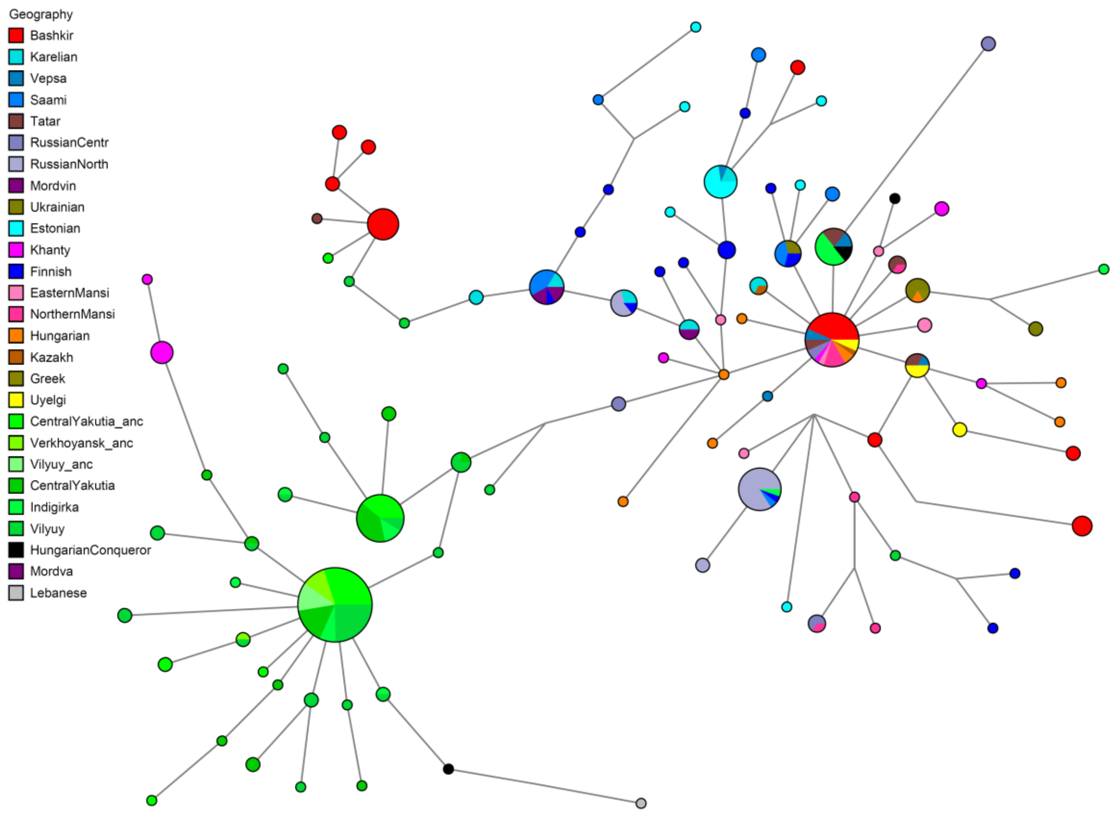

**Fig. S12 Median Joining Network analysis of N1a1-M46 Y- haplogroup using 12 STRs.** Two Uyelgi samples (Uyelgi1 and Uyelgi5) share identical 12 STRs with further 28 modern individuals, which is visible on the reduced N1a1-M46 Network tree using 12 STR data: 13 Bashkirs and two Tatars from Volga-Ural region, five Mansi and one Khanty individuals from Western Siberia, one Kazakh from Inner Asia, two Central-Russian and two Vepsa individuals from northeast Europe, and two Hungarians from Carpathian Basin. One-step-neighbour to this group are three Uyelgi samples (Uyelgi7, Uyelgi15 and Uyelgi16), two Tatars from Volga-Ural region and one Vepsa individual from northeast Europe, as well as seven Indigirka Individuals from East Asia, further three Tatars from Volga-Ural region, two Vepsa individuals from northeast Europe and two Hungarian Conquerors from Carpathian Basin (Bodrogszerdahely-Bálványhegy/grave 3 and Karos-Eperjesszög II/grave 29). Another Hungarian Conqueror sample (Karos-Eperjesszög II/grave 14) is two-steps away from the Uyelgi1 and Uyelgi5 among others above mentioned samples (for the population information, STR data and references see Supplementary Table S8).
